# Asynchronous Transitions from High-Risk Hepatoblastoma to Carcinoma

**DOI:** 10.1101/2024.12.24.630261

**Authors:** Xinjian Yu, Stephen Sarabia, Martin Urbicain, Sonal Somvanshi, Roma Patel, Priyanka Rao, Tuan M Tran, Yen-Ping Yeh, Keng-Shih Chang, Yi-Tzu Lo, Jessica Epps, Kathleen A. Scorsone, Hua-Sheng Chiu, Emporia Faith Hollingsworth, Cintia R. Perez, Mohammad Javad Najaf Panah, Barry Zorman, Jordan Muscal, Milton J Finegold, John A. Goss, Rita Alaggio, Angshumoy Roy, Kevin E. Fisher, Andras Heczey, Sarah Woodfield, Sanjeev Vasudevan, Kalyani Patel, Ting-Wen Chen, Dolores Lopez-Terrada, Pavel Sumazin

**Author notes:** Correspondence to. Lead corresponding author: Pavel Sumazin, Texas Children’s Hospital, Feigin Tower, 1102 Bates Avenue, FC1220.09, Houston, TX 77030, USA. Equal contribution (Xinjian Yu, Stephen Sarabia, Martin Urbicain). Author Contributions: D. Lopez-Terrada and P. Sumazin conceived the work. S. Sarabia, M. Urbicain, S. Somvanshi, R. Patel, P. Rao, T.M. Tran, Y.P. Yeh, K.S. Chang, Y.T. Lo, J. Epps, K.A. Scorsone, H.S. Chiu, E. F. Hollingsworth, C.R. Perez, M.J.N. Panah, B. Zorman, M. Finegold, J.A. Goss, R. Alaggio, A. Roy, K.E. Fisher, A. Heczey, S. Woodfield, S. Vasudevan, K. Patel, T.W. Chen performed assays and analyses. X. Yu., D. Lopez-Terrada, and P. Sumazin wrote the paper. Data availability statement: All data generated in this project, including raw sequencing and CELL files resulting from whole exome sequencing, targeted panel (Texas Children’s Pediatric Solid Tumor Cancer Mutation Panel), Affymetrix OncoScan, and single-nuclei RNA sequencing, single-cell RNA sequencing, and single-nuclei DNA sequencing profiles, are freely available at the European Nucleotide Archive, accession PRJEB82511. Human ethics: All human participants were fully consented under Texas Children’s Hospital IRB H-13999.

## Abstract

**Background:** Pediatric hepatocellular tumors are most commonly classified as hepatoblastoma (HB) or hepatocellular carcinoma (HCC), yet a subset exhibits mixed histological and molecular features of both. These tumors, previously described as hepatoblastomas with carcinoma features (HBCs), include cases provisionally designated by the World Health Organization as hepatocellular neoplasm–not otherwise specified (HCN-NOS). Their biology remains poorly understood, with unresolved questions about their cellular composition and outcomes. In particular, it is unclear whether HBCs comprise hybrid cell populations with combined HB and HCC characteristics (HBC cells) or admixtures of distinct HB and HCC cells.

**Methods:** We performed multi-omics profiling—including snRNA-seq, snDNA-seq, and multi-region longitudinal bulk RNA and DNA sequencing—to characterize HBC composition, evolution, and treatment response.

**Results:** HBCs comprise heterogeneous mixtures of HB-like, HBC-like, and HCC-like molecular cell types. HBC outcomes are significantly worse than HB, and HBC cells are more chemoresistant than HB cells, with resistance shaped by their cell identity, genetic alterations, and embryonic differentiation stage. HBC cells originate from HB cells that were arrested at early hepatic stem cell development stages because of aberrant WNT signaling activation. Inhibition of WNT signaling promoted differentiation and enhanced sensitivity to chemotherapy. Furthermore, each analyzed HBC reflected a dynamic process of multiple HB-to-HBC and HBC-to-HCC transitions, underscoring their evolutionary complexity.

**Conclusions:** Multi-omics profiling of HBCs revealed key findings about their biology and composition, demonstrating that they originate from HB precursors at early hepatic stem cell development stages and that their differentiation arrest depends on sustained aberrant WNT-signaling activity.

## INTRODUCTION

HB and HCC are the most frequent primary hepatocellular tumors diagnosed in children [1]. However, some hepatocellular tumors demonstrate combined HB-HCC molecular and histological features. Some of these tumors are recognized under the provisional HCN NOS diagnostic category [2]. Initially, HCN NOS were identified based on histopathology features alone [3, 4]. However, recent approaches combined immunohistochemistry and molecular biomarkers to diagnose HCN NOS and related tumors under the broader hepatoblastoma with carcinoma features (HBC) category [5]. Consequently, we distinguish between three types of childhood hepatocellular tumors: HB, HBC, and HCC [5, 6].

HBC histology suggested classification into three subtypes: biphasic, which contain homogenous HB and HCC regions; equivocal, which are composed of intermingled cells with HB and HCC features; and high-grade focal, which, before and after chemotherapy, contain small areas displaying high-grade HB-HCC features—including atypia and pleomorphism—enveloped by HB cells. The molecular profiles of all HBC subtypes revealed recurrent genetic alterations alongside the activation of both canonical HB and HCC pathways. Because HBs tend to be more chemoreceptive, post-treatment HBCs are often enriched with HBC and HCC-like regions (Figure S1) [5].

A key unresolved question is whether HBCs are solely composed of a mixture of HB and HCC cells or whether they also include cells with HBC-specific DNA alterations and gene expression programs (HBC cells). Moreover, the incidence rate of HBC remains unknown, and the outcomes have been reported to be comparable to either those of HB or HCC [7, 8]. In summary, HBC biology and etiology remain poorly understood, and strategies for their treatment are being investigated [9]. In this study, we characterized the cellular composition of HBCs and the phylogenetic relationships among their constituent subclones. Our research offers valuable insights into HBC etiology, their responses to therapies, and the conditions necessary for HBC initiation.

## RESULTS

### HBC outcomes are poor

To compare HBC patient outcomes to outcomes of HBs, we have collected clinical data for 42 HBCs, including 41 with recorded events for at least 2 years post-diagnosis (Table S1). HBCs diagnoses were determined through a multi-institutional consensus based on histological and molecular features. Studies that clinically characterized HBs at scale include the Children’s Hepatic Tumors International Collaboration with 1605 HBs with at least 2 years of follow-up post-diagnosis [10]; Cairo et al. (2008 and 2020) with 85 and 174 HBs, respectively [11, 12]; Sumazin et al. (2017) with 69 HBs [13]; Nagae et al. (2021) with 139 HBs [14]; and Hirsch et al. (2021) with 86 HBs [15]. A comparison of event and outcomes data suggests that HBCs have significantly worse event-free survival (Figure 1A) and overall survival (Figure 1B) than HBs, with five-year overall survival rates of 43% for HBCs and 79% for HBs on average.

**Figure 1.**
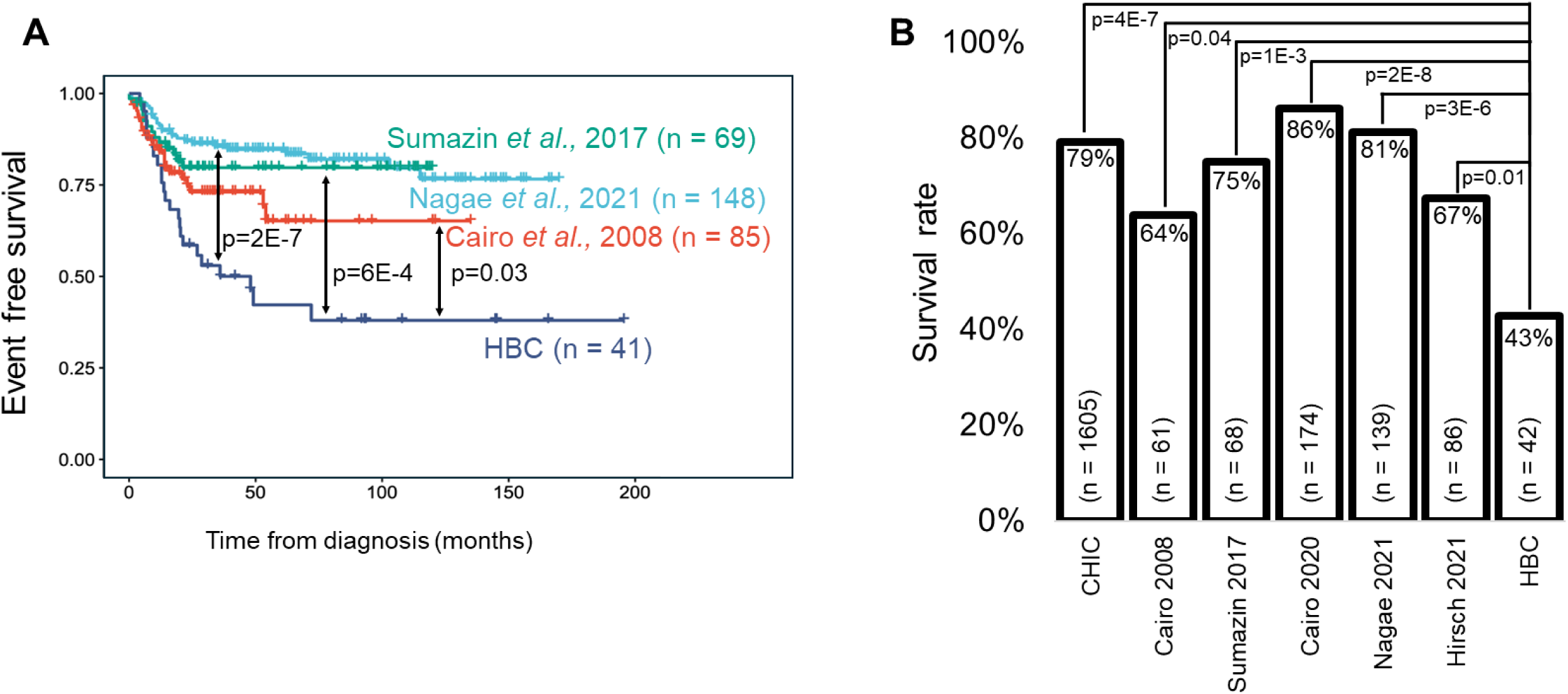
HBCs exhibit poor clinical outcomes. (**A**) Event-free survival analysis for our HBC cohort compared with three HB cohorts with recorded events; p-value by log-rank test. (**B**) Overall survival rates—based on at least two years of post-diagnosis surveillance—were compared between our HBC cohort and six HB cohorts with available patient outcome data; p-values by Fisher’s exact test. CHIC: Children’s Hepatic Tumors International Collaboration.

### HBC resections contain HBC cells

To determine whether HBC tumors contain cells with both HB and HCC molecular features (HBC cells), we developed a gene expression-based classifier that scores cells for HB and HCC features, allowing clear distinction between HBs and HCCs (Figure S2A and Table S2). Non-cancer cells were negative for both feature sets (Figure S2B), whereas HBC cells were expected to score highly for both HB and HCC features. This HB-HCC classifier accurately identified HB and HCC cells in samples that were profiled at single-cell (Figures 2A and S2B and Table S3) and bulk resolutions (Figure S2C and Table S4).

**Figure 2.**
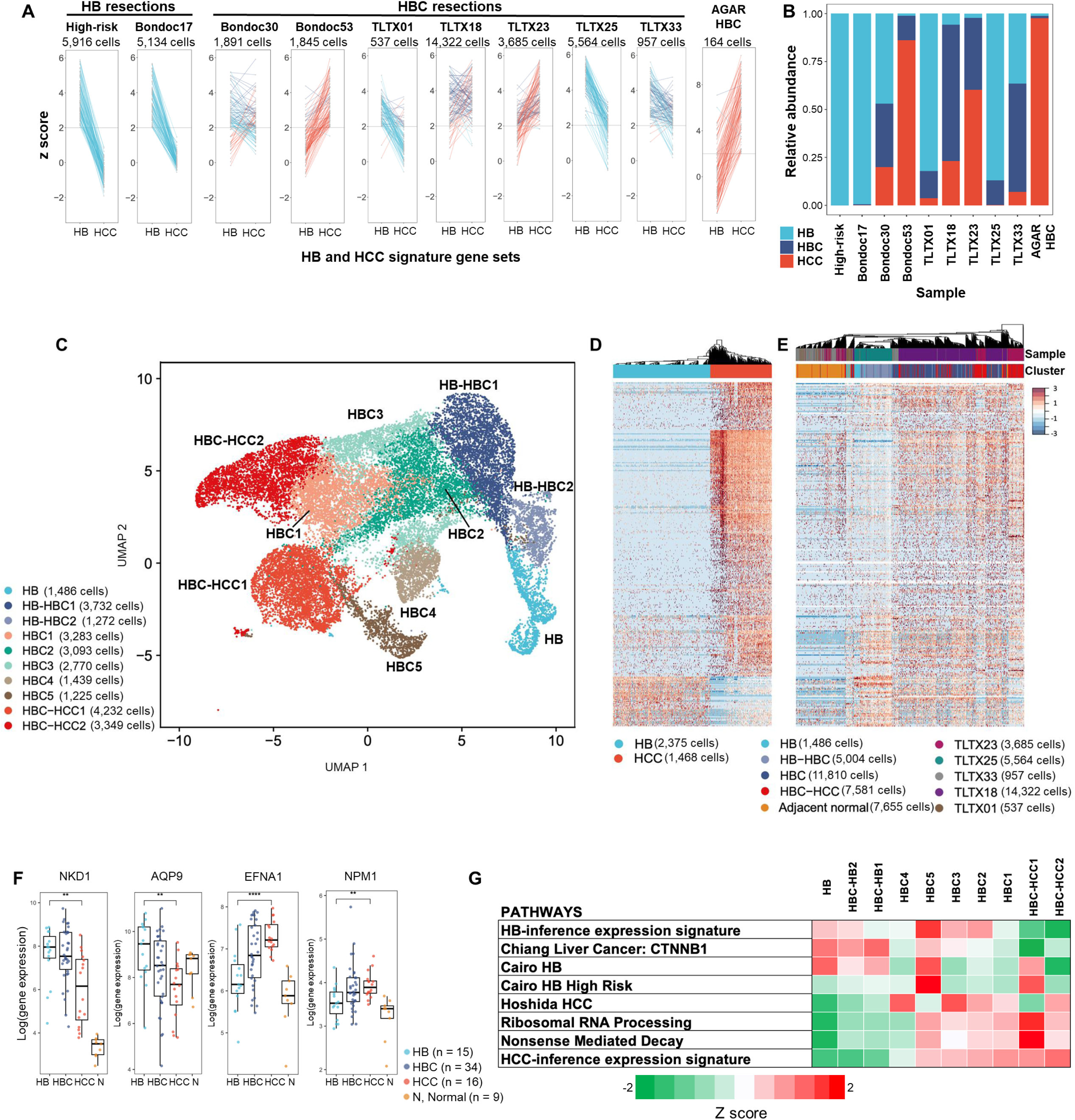
HBC tumors comprise HB, HBC, and HCC cells. (**A**) Upregulation of HB and HCC signature gene sets, represented by z-scores, in cells from two HB resections, seven HBC resections, and two AGAR-trial HBC biopsies. For clarity, data for 100 randomly selected cells from each sample are shown. HB and HCC scores for each cell are connected by a line, with high scores for HB, HCC, or both gene sets indicating HB, HCC, and HBC cells, respectively. These samples were not used for biomarker discovery. (**B**) Estimated cellular composition of each sample. A z-score > 2 or a z-score difference > 2 indicates significant evidence for HB or HCC identity, whereas cells with significant and comparable scores for both sets were classified as HBCs. (**C**) UMAP of 25,881 tumor cells from five HBC resections (TLTX samples in panels A and B) illustrating transcriptomic clusters corresponding to distinct cancer cell types. HB–HBC and HBC–HCC clusters include HBC cells transcriptionally proximal to HB and HCC, respectively. (**D**) Unsupervised hierarchical two-dimensional clustering of cell profiles from HB and HCC resections; columns and rows represent cells and genes, respectively. (**E**) Unsupervised hierarchical one-dimensional clustering of cells from the five HBC resections (TLTX samples), with genes ordered as in panel D. Annotations indicate the sample of origin and cluster type. (**F**) Expression of selected HB and HCC biomarkers in HB, HBC, HCC, and non-cancer liver (normal) samples profiled using the NanoString nCounter assay. *** p < 0.0001; ** p < 0.001; * p < 0.01. (**G**) Differentially expressed pathways across clusters shown in panel C.

We applied this classifier to single-cell transcriptomic profiles from resection samples of two HBCs profiled by Bondoc et al. [16], four biphasic HBCs (TLTX01, TLTX23, TLTX25, and TLTX33), and one equivocal HBC (TLTX18). As supplementary controls, we also analyzed profiles of one HB sample and two patient-derived xenografts (PDXs) from Bondoc et al., a resection of a high-risk HB, and two HBC biopsies from the AGAR trial (Figures 2A and S2D,E) [17]. Because AGAR patients underwent intensive therapy and HBs are less chemoresistant, AGAR HBCs were expected to contain few HB cells [18].

Across all HBC resections—including biphasic tumors expected to contain histologically homogeneous regions—we detected HBC cells (Figures 2A,B and S2D,E; Table S5). As anticipated, HB samples consisted exclusively of HB cells, whereas AGAR HBCs were dominated by HCC cells (93%). Notably, HBC cells accounted for more than half (51%) of all malignant cells in HBC tumors.

### Both HB and HCC signature genes are upregulated in HBC cells

An unresolved question in HBC biology is why some bulk RNA-expression profiles of HBC tumors cluster with those of HBs or HCCs, while others form an intermediate cluster characterized by concurrent upregulation of both HB- and HCC-specific genes [5]. Our analysis indicated that some HBC cells more closely resemble HBs, others resemble HCCs, and some are transcriptionally distinct from both (Figures 2C-E and Table S6). Consequently, HB and HCC cells from each HBC tumor clustered with their respective counterparts regardless of tumor origin, while the HBC cells exhibited a transcriptomic spectrum from HB to HCC.

Given their transcriptomic range, we found that the bulk RNA-expression profile of an HBC tumor reflects its cellular composition and dictates whether it resembles HBs, HCCs, or neither (Figure S2F). For instance, in TLTX18, 71% of cells were classified as HBCs, 23% as HCCs, and 6% as HBs. In hierarchical clustering (Figure 2E), cells were clustered with other cells of their type. TLTX18, however, was clustered with other intermediate HBC tumors (Figure S2F). In contrast, TLTX25, composed predominantly of HB cells, clustered with HBs, while AGAR ESFT, F HBCs (93% HCC cells) clustered with HCCs (Figure S2F). Genes and pathways that were differentially expressed across tumor types had intermediate expression patterns in HBC cells and tumors (Figures 2F,G and Table S7).

### High-risk HB, HBC, and HCC cells are less differentiated

Genes associated with stem cells and pluripotency are recurrently upregulated and genomically altered in HBCs [5]. To investigate the embryonic differentiation states of HBCs, we developed a model that infers the differentiation stage of liver cells based on single-cell RNA sequencing (scRNA-seq) profiles of human embryonic liver cells at 5 and 19 weeks post-conception (Figure S3A) [19]. This model estimates the differentiation state of individual cells and bulk samples from their RNA expression profiles.

Our analyses revealed clear developmental distinctions among tumor types. Pre-treatment HBs were predominantly composed of cells at later embryonic differentiation stages, whereas pediatric HCCs were enriched in cells at earlier embryonic stages. In contrast, post-treatment HBs and pediatric HCCs contained the most embryonically undifferentiated cell populations, suggesting a depletion of HB cells that are at later embryonic differentiation stages post therapy (Figure S3F and Table S9). Indeed, comparative analysis of scRNA-seq–profiled HBs [20] suggested that low-risk tumors are enriched in cells at later differentiation stages (Figure S3G; Table S9) and that HBCs are significantly less differentiated than low-risk HBs (Figures S3F–H; Table S9; p = 0.02, t-test). Likewise, HB differentiation scores and the expression of LIN28B—a known marker of early developmental states [13, 23]—were correlated with established clinical risk categories (Figures 3A,B and Table S10).

**Figure 3.**
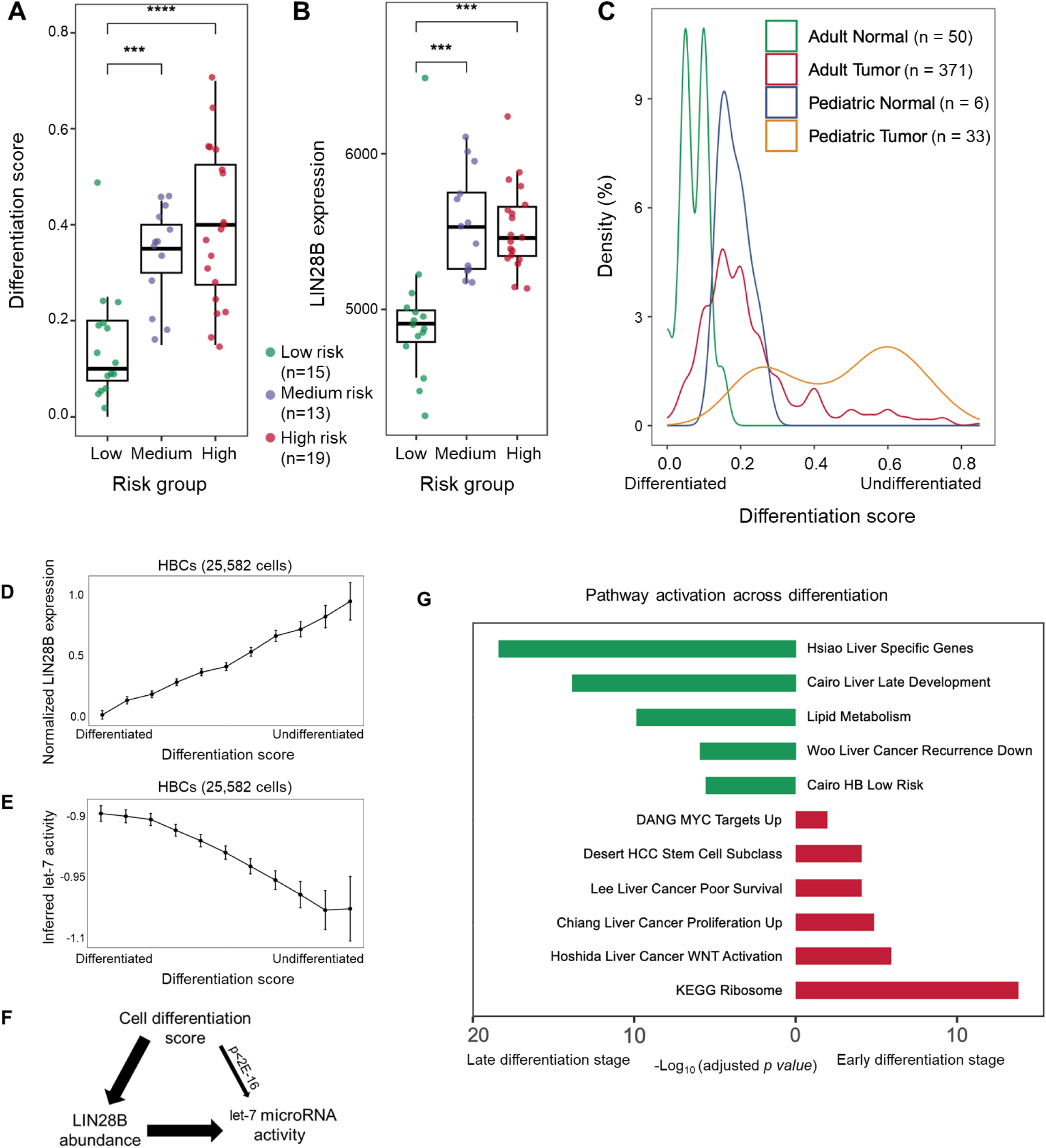
Tumor embryonic differentiation stages predict clinical risk. (**A**) Tumor embryonic differentiation scores and (**B**) LIN28B expression levels predict HB risk; **** p < 0.0001; *** p < 0.001; error bars represent S.E.M. (**C**) Distribution of embryonic differentiation scores across pediatric and adult liver cancer and non-cancer samples. (**D**) LIN28B expression and (**E**) inferred activity of the let-7 miRNA family plotted by embryonic differentiation stage. (**F**) Data processing inequality (DPI) analysis of the relationships between embryonic differentiation, let-7 activity, and LIN28B expression indicates that LIN28B mediates the association between embryonic differentiation and let-7 activity. (**G**) Pathways enriched in early (differentiation score ≥ 0.45, red) and late (≤ 0.2, green) embryonic differentiation stages in HBC resections.

Comparative analyses across pediatric and adult non-cancerous and cancerous samples demonstrated that adult HCCs [21] and fetal liver tissues are enriched for cells at early embryonic stages relative to adult liver (p<2E−16 and p=7E−4, respectively, t-test). However, pediatric liver cancers were more undifferentiated than adult cancers and non-cancer liver samples, displaying a bimodal distribution of differentiation scores spanning from adult-HCC-like to highly undifferentiated states (Figure 3C; Table S10; p=7E−10, t-test). Furthermore, inferred proliferative activity indicated that undifferentiated HBC cells tend to be more quiescent (Figure S3I), a feature associated with chemoresistance [22]. Together, these findings suggest that enrichment for cells at early embryonic differentiation stages is linked to increased chemoresistance in both HBs and HBCs.

We next examined genes and pathways associated with embryonic differentiation, including LIN28B (Figure 3B). LIN28B expression correlated with differentiation stage in embryonic liver, HBs, and HBCs (Figures 3D and S3J). Consistent with its established role [24–26], LIN28B expression was inversely correlated with let-7 microRNA activity in HBC cells (Figure 3E). By suppressing let-7 biogenesis, LIN28B maintains the stem-like properties of hepatic stem cells and hepatoblasts during liver development [27]. Information-theoretic analyses further revealed that the relationship between embryonic differentiation and let-7 activity depends on LIN28B abundance, suggesting that LIN28B modulates let-7 activity in HBCs (Figure 3F; Table S11). Finally, pathway-level analyses showed that HCC- and high-risk– associated pathways were enriched in undifferentiated cells, whereas mature liver–associated pathways were activated in differentiated cells (Figures 3G and S3K–L; Table S12).

### Aberrant WNT-signaling activation regulates HBC embryonic differentiation

We hypothesized that aberrant activation of WNT signaling contributes to the undifferentiated state of HBCs. DNA alterations in WNT-pathway genes drive constitutive WNT activation in nearly all HBs and HBCs [28–30], and the expression of DKK1—a marker of WNT-signaling activity—is a prognostic biomarker that is correlated with LIN28B expression [13]. Consistent with its role in embryonic liver development, WNT signaling activity declines sharply between 5- and 9-weeks post-conception, as reflected by decreasing DKK1 expression (Figure S4A). Although this decline is also observed in HB and HBC cells, DKK1 expression remains significantly elevated in HBCs—even in relatively differentiated HBC cells (Figures S4B,C).

To directly test the role of WNT signaling in regulating cancer cell differentiation, we treated the undifferentiated, patient-derived HBC cell line HB17 [9] with the WNT-inhibitor ICG-001 [31]. While HB17 cells typically differentiate slowly in culture, treatment with ICG-001 at low toxicity significantly reduced both DKK1 and LIN28B expression (Figure S4D, Table S13). These results were replicated with a second inhibitor (WNT-C59) and confirmed using a WNT-reporter assay (Figures S4E,F).

To assess WNT inhibition effects at the single-cell level, we performed scRNA-seq profiling of HB17 cells treated with ICG-001 (5 μM) for either (i) 10 days or (ii) 5 days followed by 5 days without the inhibitor. Comparative transcriptomic analysis revealed a pronounced shift in the single-cell landscape following treatment (Figures 4A and S5A). Notably, WNT inhibition induced differentiation: treated HB17 cells transitioned toward more differentiated transcriptional states at a significantly higher rate than placebo-treated cells (Figure 4B, Table S14).

**Figure 4.**
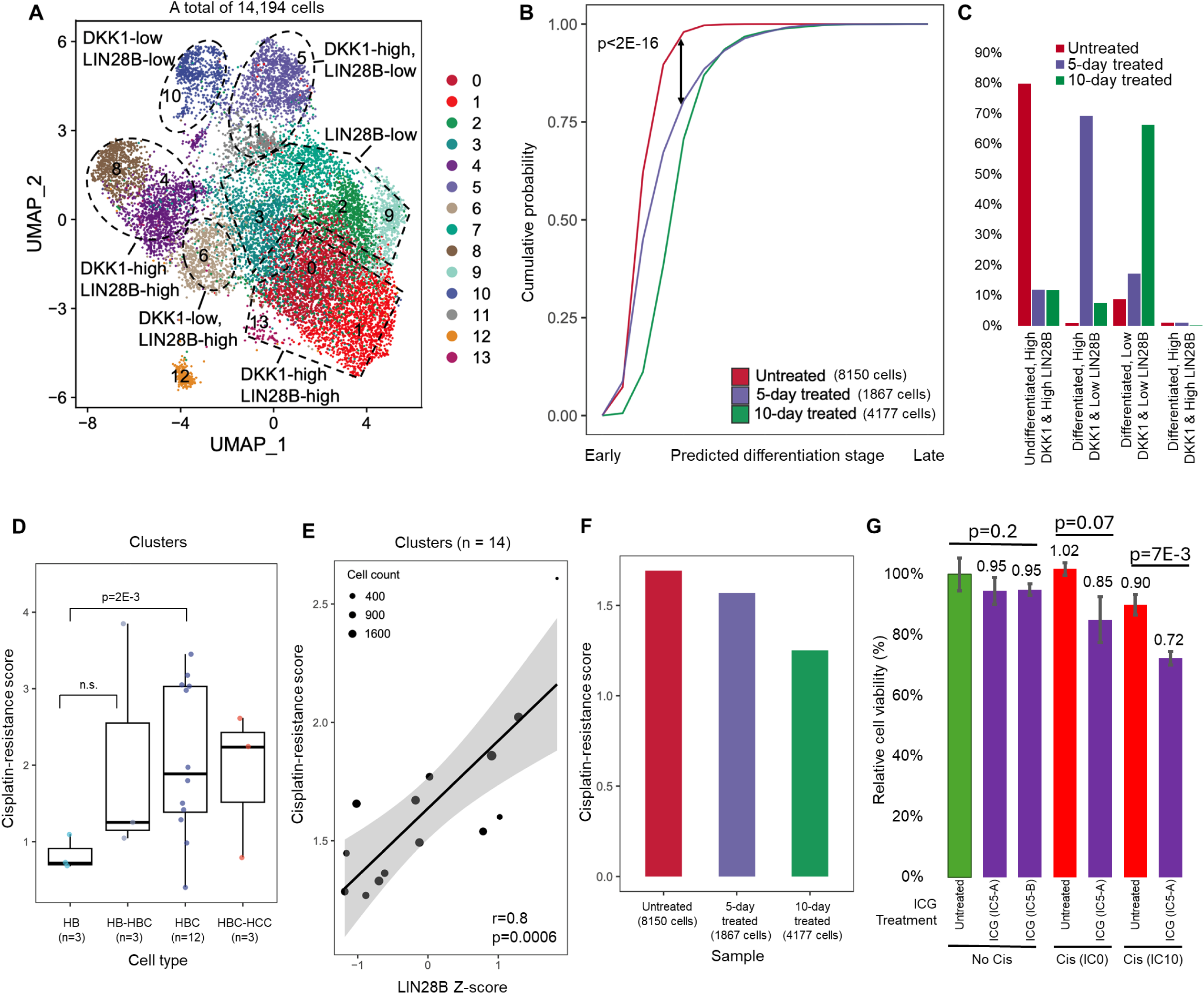
WNT signaling regulates liver differentiation and chemoresistance. (**A**) UMAP of scRNA-seq profiles from 14,194 patient-derived HB17 cells showing populations with variable expression of the embryonic differentiation marker LIN28B and the WNT-activity marker DKK1. Cells were cultured for ten days without treatment, treated with the WNT inhibitor ICG-001 for five days followed by five days without the inhibitor, or treated continuously for ten days; see Figure S5. (**B**) Cumulative distributions of inferred embryonic differentiation stages of HB17 cells under each treatment condition; p-value by chi-squared test. (**C**) Proportion of HB17 cells by differentiation stage and LIN28B/DKK1 expression across treatment conditions. Seventy percent of cells treated for five days and then rested regained aberrant WNT signaling but remained differentiated; fewer than 3% of differentiated cells exhibited high LIN28B and DKK1 expression, indicating a low misclassification rate. (**D**) Inferred cisplatin resistance across clusters from five HBC resections (TLTX samples in Figure 2); p-values by t-test. (**E**) Correlation between inferred cisplatin resistance and LIN28B expression in HB17 clusters across the three treatment conditions. Circle sizes reflect cluster sizes; Pearson correlation coefficient (r) and p-value shown. (**F**) Inferred cisplatin resistance in untreated and ICG-001–treated HB17 cells. (**G**) Relative HB17 cell viability following low-dose ICG-001 (IC₅), cisplatin (IC₀ or IC₁₀), or combination treatments compared with untreated controls. Cells were treated with ICG-001 for five days before resting (IC₅-A) or rested before treatment (IC₅-B). S.E.M. shown; Cis, cisplatin.

Differentiation scores were tightly correlated with LIN28B expression levels (Figures 4A,C), although LIN28B was not used to infer differentiation. After 10 days, ∼80% of placebo-treated cells retained high DKK1 and LIN28B expression and remained undifferentiated. In contrast, ∼70% of ICG-001–treated cells exhibited low DKK1 and low LIN28B expression, achieving differentiation scores comparable to fetal liver (Figure S3E). Interestingly, cells treated for 5 days and then cultured drug-free for 5 additional days predominantly exhibited high DKK1 but low LIN28B expression, consistent with a more differentiated, post-inhibition state. This suggests that WNT inhibition promotes differentiation, and that once differentiation is initiated, it is not reversed even after WNT activity recovers (Figures 4C, S5A–C, Table S15). As a control, few cells with high DKK1 and LIN28B expression were inferred to be differentiated, indicating that the observed differentiation effects were robust and not attributable to noise.

### WNT-signaling inhibition sensitizes HBC cells to cisplatin by promoting cell differentiation

Our analysis linked aberrant WNT-signaling activation to the maintenance of embryonically undifferentiated states in HBC cells and showed that WNT inhibition promotes their differentiation. Because undifferentiated liver cancer cells are more chemoresistant, we next evaluated whether WNT-signaling inhibition could reduce resistance to cisplatin, a standard component of pediatric liver cancer therapy [32, 33].

Towards this goal, we developed a model to infer cisplatin resistance from bulk RNA expression profiles coupled with observed resistance of pediatric liver cancer cell lines [15] (Figure S6A). The model accurately estimated cisplatin resistance from scRNA-seq profiles of treated patient-derived tumor spheroids [20] (Figure S6B), predicted HB risk categories (Figure S6C, Table S16), and identified HCC cells as significantly more resistant to cisplatin than HB cells (Figure S6D). It also detected an increase in mean cisplatin resistance following treatment in a partially chemoresistant HBC PDX (Figure S6E). We found that, on average, HBC cell clusters exhibited greater cisplatin resistance than their HB counterparts in our snRNA-seq data (Figure 4D, Table S17; p=0.002). Moreover, LIN28B expression significantly correlated with cisplatin-resistance scores in ICG-001–treated HB17 cells, suggesting that differentiation reduces resistance to cisplatin (Figure 4E, Table S17). Consistently, average chemoresistance scores decreased following ICG-001 treatment (Figure 4F).

To test whether cell differentiation enhances chemosensitivity, we treated HB17 cells with ICG-001 (5 μM, IC₅) using a 5-day-on/5-day-off regimen, followed by cisplatin at non-toxic (IC₀) or low-toxicity (IC₁₀) doses. Co-treatment with ICG-001 and cisplatin significantly reduced cell viability compared to cisplatin alone, with IC₁₀ cisplatin reducing viability by 28% after treatment with IC₅ ICG-001 (p=7E-3, Figure 4G). These findings indicate that WNT-signaling inhibition promotes differentiation and increases cisplatin sensitivity in HBC cells.

### HBCs are derived from HBs at early hepatic stem cell development stages and transition to HCC

Having established that HBC tumors contain cells with HB, HBC, and HCC molecular features, we next investigated the phylogenetic relationship between these cancer cell types. To reconstruct tumor phylogenies, we integrated data from multiple assays, including multi-region longitudinal whole-exome sequencing (WES) [34], Affymetrix OncoScan profiles [5], single-nuclei DNA-seq (snDNA-seq) [35, 36], Texas Children’s Pediatric Solid Tumor Cancer Mutation Panels [37, 38], NanoString bulk RNA expression profiles [5], and snRNA-seq [39]. Samples and patients are listed in Table S1. TLTX36 regions collected before and after treatment were analyzed at bulk resolution, whereas multiple post-treatment regions from TLTX01, TLTX18, TLTX23, TLTX25, and TLTX33 were profiled using a combination of bulk, snDNA-seq, and snRNA-seq assays.

The inferred TLTX36 phylogeny identified alterations shared across genetic subclones and indicated that TLTX36 HBC cells originated from HB progenitors. The pre-treatment biopsy was primarily composed of embryonically differentiated (fetal) HB cells, whereas post-treatment resections contained a higher proportion of embryonically undifferentiated and genetically less-stable HB and HBC cells (Figure 5A; Tables S19, S20). Quantitative histologic annotations of fetal and embryonal compositions, performed by a team of clinical pathologists (DLT and KP), were consistent with the inferred embryonic differentiation scores (Table S19).

**Figure 5.**
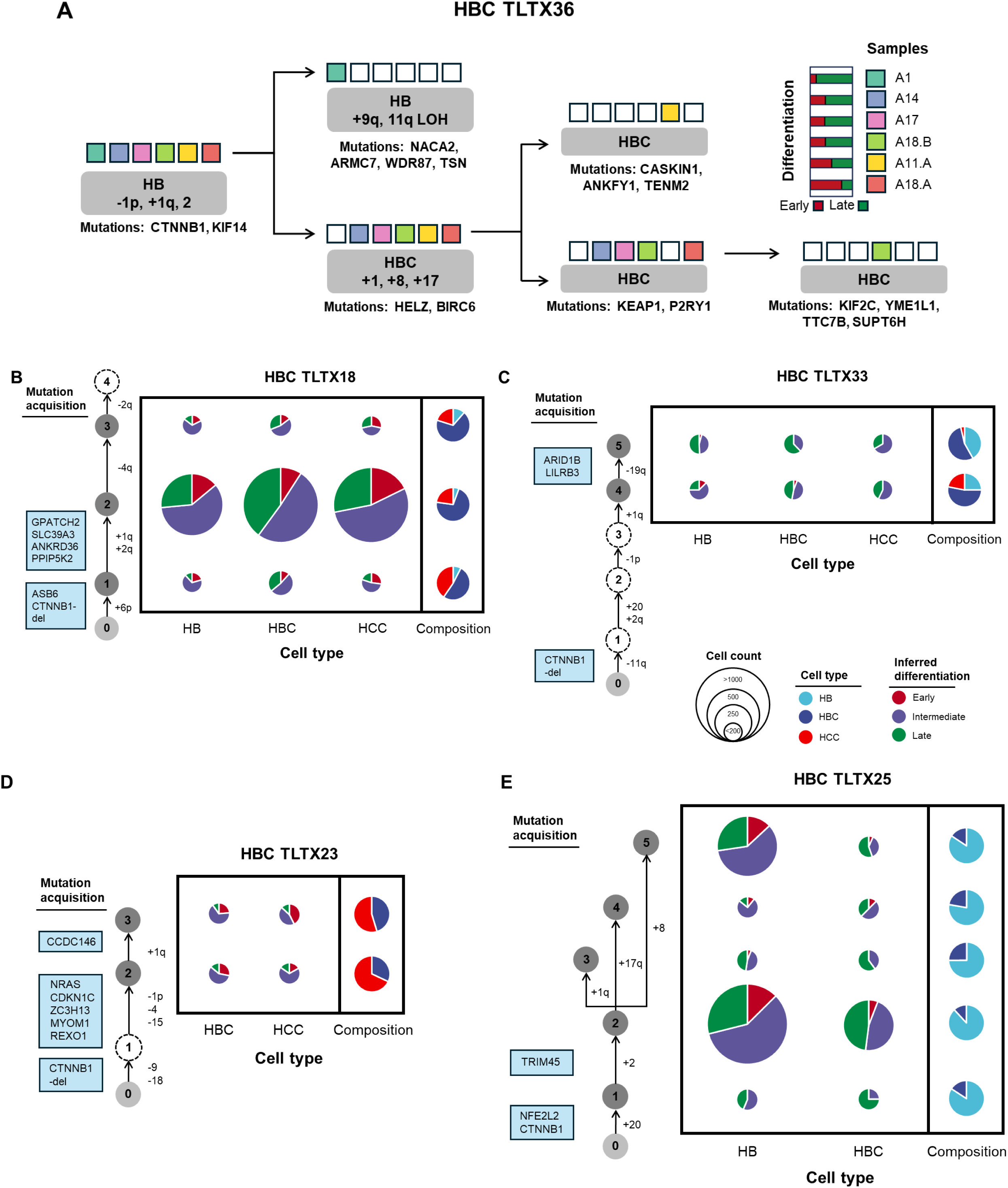
HBC phylogeny and composition reveal multiple transition events during hepatic stem cell development. (**A**) Phylogeny of HBC TLTX36 inferred from whole-exome sequencing of a pre-treatment biopsy (A1) and five regions of a post-treatment resection (A14, A17, A18.B, A11.A, A18.A). Each genetic subclone is annotated by the region in which it was detected. Embryonic differentiation scores for each region were derived from NanoString RNA profiles. LOH, loss of heterozygosity. (**B–E**) Phylogenies, cellular compositions, and differentiation stages of HBC tumors (**B**) TLTX18, (**C**) TLTX33, (**D**) TLTX23, and (**E**) TLTX25 inferred from single-cell RNA and DNA profiles, whole-exome sequencing, and SNV array data across multiple regions. Phylogenies incorporate somatic mutations and copy-number alterations; circle sizes represent cell counts, and colors indicate differentiation stages. Relative tumor-type compositions (HB, HBC, and HCC) are also shown for each genetic subclone, with colors indicating cell types.

Mutations and copy-number alterations in HBC cells were supported by data from OncoScan, snDNA-seq (Figure S7), snRNA-seq (Figure S8), and WES (Figure S9). Cells were grouped into genetic subclones based on shared DNA alterations, and each subclone was further classified by cancer cell type (HB, HBC, or HCC) and embryonic differentiation stage (early, intermediate, or late; Figures 5B–E). TLTX18 and TLTX33 contained four and five distinct genetic subclones, respectively, as determined by WES and snDNA-seq, but of these, only three (TLTX18) and two (TLTX33) were sufficiently represented in snRNA-seq profiles to permit downstream analysis.

The inferred phylogenies for TLTX18 and TLTX33 (Figures 5B,C) demonstrated that each genetic subclone encompassed cells from all three cancer cell types, with subclonal populations enriched in cells at early hepatic stem cell differentiation stages. Although only limited numbers of HB cells from TLTX23 and HCC cells from TLTX25 were profiled by snRNA-seq, these tumors exhibited similar patterns, with each genetic subclone containing undifferentiated cells from two tumor types (Figures 5D,E). Given the low probability that cells of distinct tumor types and differentiation states independently and recurrently acquired identical genetic alterations, these phylogenies indicate that transitions from HB to HBC and from HBC to HCC occur early in embryonic liver differentiation, across genetically distinct subclones, and within each HBC tumor.

## DISCUSSION AND CONCLUSIONS

The provisional HCN NOS diagnostic category was introduced during the *International Pediatric Liver Tumors Consensus* at the International Pathology Symposium in 2014 [2] and is recognized by the *World Health Organization*. HCN NOS and related tumors, including HBs exhibiting focal atypia or pleomorphism, share common clinical, histological, and molecular features. Collectively, these tumor types are referred to as HBCs [5]. Bulk molecular profiles of these tumors show that both HB- and HCC-specific gene expression and pathway activity are elevated in HBCs. However, HBC bulk profiles did not resolve key questions regarding their cellular composition and intratumoral heterogeneity.

Our single-cell analyses revealed that each HBC tumor contained both HB-like and HCC-like cells with expression patterns similar to those found in HB and HCC tumors, respectively. However, most cells in HBCs exhibited intermediate transcriptional states ranging from HB-like to HCC-like (Figure 2E), consistent with the presentation of HBC cells. When clustered hierarchically, some HBC cells more closely resembled HB or HCC, whereas others were transcriptionally distinct enough to form a separate grouping in relative to both HB and HCC transcriptomes.

To further explore the clonal architecture of these tumors, we constructed tumor maps that integrated inferences about clonal composition and phylogenetic relationships among genetic subclones, as well as the tumor types and embryonic differentiation states of the cells comprising each subclone (Figure 5). These maps revealed multiple asynchronous transitions between HB and HBC, and between HBC and HCC cells, within each tumor. This analysis revealed that each genetic subclone comprised a mixture of molecular cancer cell types—HB, HBC, and HCC—and contained cells arrested at early stages of hepatoblast differentiation.

Given the improbability of repeated genetic convergence or reversal of embryonic differentiation across all subclones and tumors, we inferred that each HBC tumor arises through multiple asynchronous transitions originating from HB cells at very early differentiation stages. Furthermore, inhibition of aberrant WNT signaling promoted HBC differentiation, indicating that this pathway may play a key role in maintaining the undifferentiated state of HBC cells. We conclude that the phenotypic heterogeneity of HBCs reflects both genetic instability and variability in embryonic differentiation and cell states. Notably, 84% of HBCs harbored mutations commonly associated with HCC that preceded HB-to-HBC transitions, suggesting that these alterations may facilitate such transitions (Table S1).

Our study has several limitations. First, although our cohort included 42 HBCs, only a subset was analyzed in depth using single-cell resolution assays; thus, these findings should be validated in a larger cohort. Second, nearly all molecular profiles were derived from post-treatment resections, in which HB cells were expected to be underrepresented due to their lower chemoresistance. Large-scale comparisons of pre- and post-treatment samples are needed to assess the impact of chemotherapy on tumor composition and cell states. Finally, while we demonstrate that HBC tumors arise through asynchronous transition events, the specific molecular drivers underlying these transitions remain to be identified.

## MATERIALS AND METHODS

Below, we summarize our materials and methods, with details given in Supplementary Materials.

### snRNA-seq, snDNA-seq, and scRNA-seq protocol, preprocessing, and analysis

snRNA-seq assays were performed as previously described [39] using frozen tumor resections after adjuvant chemotherapy. Tissues were profiled, analyzed, integrated, and annotated [34, 40].

### Whole exome and targeted sequencing and analysis

A total of 52 samples were profiled by both whole-exome sequencing and the Texas Children’s Hospital’s Solid Tumor Comprehensive Panel [41], producing inferred mutations and copy number alterations.

### Inferring CNVs by Affymetrix OncoScan

Genome-wide copy numbers were estimated using DNA isolated from formalin-fixed paraffin-embedded (FFPE) tumor tissue using the Affymetrix OncoScan, as previously reported [5].

### RNA expression profiling by NanoString

We profiled the gene expression of each FFPE tissue region using a custom 814-probe NanoString nCounter panel (Tables S21 and S22), as previously reported [5].

### Tumor histopathology and patient characterization

We characterized HBCs with review board approval. Histological reviews of glass slides, including diagnostic biopsies and resection, transplant, and metastasis samples, were performed by MJF, DLT, RA, and KP, who confirmed the features in committee and selected representative tumor areas for molecular evaluation. HBC NOS and HBs demonstrating focal atypia or pleomorphism were identified as HBCs. Representative tumor areas were selected from post-chemotherapy resections based on displays of heterogeneity, atypia, or pleomorphism; see representative pre- and post-treatment HBC regions (Figures S1A-B) and representative slides from each cancer subtype (Figure S10).

### HB17 Culturing and drug-response assays

The HB17 patient-derived cell line was cultured, treated, and evaluated as previously described [9, 42, 43]. Cells were treated with either ICG-001 or WNT-C59, and the responses to DKK1 and CTNNB1 knockdowns in isolation or in combination with ICG-001 or WNT-C59 were evaluated (Figures S4D-G).

### Reconstructing evolutionary trees from WES and single-cell resolution profiles

TLTX36, TLTX18, TLTX23, TLTX33, and TLTX25 tumor phylogenies were reconstructed based on the WES, snDNA-seq, and snRNA-seq profiles of tumor and tumor-adjacent regions, as well as blood samples as controls [34].

### HB vs. HCC subtype classification

HB and HCC biomarker sets included genes that are differentially expressed between HB and HCC cells and versus non-tumor cells. Samples used in classifier training include cells from (A) three HB and eight adjacent normal hepatocyte samples [20], (B) HCC and adjacent normal samples from two pediatric AGAR-trial patients [17], (C) one HB sample, and (D) one HCC sample (Figure S2A). The classifier’s performance was tested on samples not included in training: cells from (A) a high-risk HB resection, (B) an HB sample and its corresponding PDX [16] (Figure S2E), and (C) bulk RNA profiles (Figure S2C). The classifier was used to evaluate cells in (A) two HBC samples and a PDX [16], (B) our five HBCs (TLTX tumors), and (C) two AGAR-trial HBCs (Figure 2A).

### Liver cell embryonic differentiation inference

We trained a random-forest classifier to predict embryonic liver differentiation using scRNA-seq profiles of liver cells from 5W and 19W post conception [19]. Cells from intermediate time points (6W-16W) and embryonic mouse liver samples were used to evaluate the method.

### Cisplatin-resistance prediction

Our classifier was trained to produce resistance scores using RNA expression and cisplatin resistance estimates in nine cancer cell lines (Figure S6A, training p=2E−4) [15]. Resistance scores were evaluated on low- and high-risk HBs [13] (Figure S6C, p<0.01), our HB and HCC samples (Figure S6D), an HB PDX before and after treatment with cisplatin (Figure S6E), and four HB organoids [20] (Figure S6B, p<0.04). When evaluating the score on HBC clusters, each cluster was evaluated independently (Figure 2C).

### Evaluation of HBC proliferation vs. quiescence

We estimated the proliferative activity of HBC tumor cells across differentiation stages using cell cycle S-related genes identified by Whitfield et al [44].

### Estimating miRNA activity and data processing inequality from snRNA-seq data

We used predicted miRNA targets to estimate *let-7*-family miRNA activity. Pairwise normalized mutual information (MI) was calculated between (i) inferred *let-7* activity, (ii) LIN28B expression profiles, and (iii) cell differentiation scores.

## Supporting information

All tables

## SUPPLEMENTARY METHODS

### snRNA-seq protocol

To prepare nuclei, we added cold lysis buffer (NST-DAPI buffer with 0.1-0.2 U/μL murine RNase inhibitor) to frozen tumor tissues and cut the tissues into tiny pieces with scalpels. We used 1.5 mm beads to isolate nuclei by using a bead blaster at 4.0 m/s for three cycles of 20s on, with 10s pauses between cycles. We incubated the nuclear extract on ice for 5 min and passed it through a fluorescence-activated cell sorting (FACS) filter. We sorted the cells using FACS (BD FACSMelody) into 6 to 10 μL RNase inhibitor in 1.5 mL low-binding tubes coated with phosphate-buffered saline (PBS) + 1% bovine serum albumin (BSA). Suspensions were filtered through a 40-μm mesh, and single nuclei were flow-sorted. The DAPI intensity was used to set gates for both diploid and aneuploid cell populations. After sorting, we centrifuged the samples at 550 *g* for 5 min at 4°C and removed the supernatant (∼20 μL left). Then, we slowly added resuspension buffer (PBS + 1% BSA + 0.2 U/μL RNase inhibitor) without resuspending the nuclei pellets and incubated the samples on ice for 5 to 10 min for buffer exchange. Finally, we centrifuged the samples at 550 *g* for 5 min at 4°C, removed the supernatant (∼40 μL left), counted the cells, and performed snRNA-seq^1^.

### Isolation of single nuclei by FACS

Frozen tumor tissue was lysed using an NST-DAPI lysis buffer as previously described^2^. Cell suspensions were filtered through a 40-μm mesh, and single nuclei were flow-sorted (BD FACSMelody). The DAPI intensity was used to set gates for both diploid and aneuploid cell populations. Single nuclei were sorted and then deposited into individual wells of 384-well plates (Eppendorf, 951020702). The sorting instrument alignment was assessed under a microscope before each experiment to ensure single nuclei were accurately deposited into the center of each well using a flat-bottom 384-well plate (Greiner, 781091).

### Acoustic cell tagmentation procedure

FACS-sorted cells in 384-well plates were spun at 1500 *g* for more than 4 min. The Echo 525 liquid handler (Labcyte) was used to dispense tagmentation reagents (Illumina, FC-131–1096) at nanoliter scales, with plate and liquid types detailed in the following steps. Thorough mixing and spinning of each plate after every dispense and incubation period is crucial to maximizing assay performance. Nuclei were lysed in 200 nL (384PP_SPHigh) of freshly prepared Tx lysis buffer (1.36 AU/mL protease diluted 1:9 in 5% Tween 20, 0.5% Triton X-100, and 30 mM Tris pH 8.0). Lysis thermocycler settings were programmed as 55°C 10 min, 75°C 15 min, 4°C ∞, lid = 80°C, vol = 1 μL. After lysis, 600 nL tagmentation reaction mixture (TD: ATM 2:1, 384PP-Plus_GPSA) was dispensed per well. The ACT reaction settings on the thermocycler were 55°C, 5 min, 4°C ∞, lid = 60°C, vol = 1 μL. The ACT reaction was neutralized with 200 nL (384PP_SPHigh) of NT buffer for 5 min at room temperature. The final polymerase chain reaction (PCR) included 1.11 μM N7XX (5′-CAAGCAGAAGACGGCATACGAGAT**<u>XXXXXXXX</u>**GTCTCGTGGGCTCGG-3′) and S5XX (5′-AATGATACGGCGACCACCGAGATCTACAC**<u>XXXXXXXX</u>**TCGTCGGCAGCGTC-3′) primers (384PP_AQBP) in 2× HiFi HotStart ReadyMix (Roche, KK2602, 6RES_GPSA). Dual barcode sequences in primers are denoted by “**<u>XXXXXXXX</u>**.” Unique dual barcode combinations for each well of a 384-well plate were achieved by dispensing 16 unique N7XX barcodes across each row and 24 unique S5XX barcodes across each column. The PCR settings were 72°C 3 min, 98°C 30 s, then 15 to 18 cycles of 98°C 10 s, 63°C 30 s, 72°C 30 s, followed by 72°C 5 min, 4°C ∞, lid = 105°C, vol = 6 μL. ACT performance was evaluated by fluorometer (Qubit) and TapeStation (Agilent) from selected cell libraries. Final libraries were pooled and purified with 1.8× AMPure XP beads. The final libraries were sequenced at 50 single-read cycles with dual barcodes on the Illumina NextSeq 2000 P3.

### Single nuclei 10x Genomics Chromium Single Cell 3′ protocols

We FACS-sorted nuclear suspensions (in PBS with 1% BSA and RNase inhibitor) and manually counted and concentrated the nuclei within the recommended range of 300 to 700 cells/μL for loading at a volume to recover 3000 to 8000 cells for snRNA-seq. Single-nuclei capture, barcoding, and library preparation followed 10x Genomics Chromium Single Cell 3′ protocols (CG000183, v3.1). Four libraries were pooled to give a final concentration of 10 nM, and the pooled samples were further tested by quantitative PCR for final concentration before submission for sequencing with the Illumina NextSeq 2000 sequencer using the P3 100 cycles flowcell. The samples were sequenced with 28 cycles for read 1, 8 cycles for i7 index, and 91 cycles for read 2 through the Advanced Technology Genomics Core at the University of Texas MD Anderson Cancer Center. Reads from single cells were demultiplexed and aligned using the human GRCh38-2020-A genome reference using the 10x Genomics Cell Ranger 6.1 pipeline for snRNA-seq.

### snRNA-seq and scRNA-seq preprocessing and clustering

The R package Seurat (v4.3.0)^3^ was used for single-cell data preprocessing and clustering. For patient snRNA or scRNA-seq data, cells with mitochondrial gene content above 15% and expressing fewer than 200 genes were excluded. The raw count matrix was normalized using the “LogNormalize” method with a scale factor of 10,000 and then scaled across cells. Principal component analysis was performed on all the genes. HBC samples profiled in two batches were integrated using canonical correlation analysis^4^. Clustering was based on the first 15 principal components and visualized by UMAP. Annotation of tumor and normal cell clusters was based on established pediatric liver cancer markers^5,6^ and the automated annotation tool SingleR (v2.0.0)^7^. Because we focused on tumor composition, only high-confidence tumor cell clusters were included in further analyses. Cell clusters with ambiguous tumor or normal assignments were excluded. HB17 cell clusters were integrated by sample using the standard Seurat package based on integration anchor identification (FindIntegrationAnchors function). When analyzing scRNA-seq profiles of PDX samples, we classified the xenograft-derived sequence read data for further analyses using Xenome^8^, which effectively handles a mix of reads from both the host and the graft. Cells from PDXs with mitochondrial gene content higher than 15% and those expressing fewer than 500 genes or more than 4000 genes were excluded to eliminate poorly or inconsistently profiled cells.

### snDNA-seq analysis

AneuFinder^9^ R package (v1.24.0) was used for snDNA-seq data preprocessing, quality control, and detecting aneuploidy. We used this tool to align reads to the GRCh38 reference genome, partition the genome into 1-Mb non-overlapping bins per cell, and apply a hidden Markov model to the binned data for unbiased detection of CNVs and aneuploidy.

### Inferring CNVs from snRNA-seq

We used the inferCNV R package (v1.8.1) to infer CNVs for each tumor cell from snRNA-seq profiles of TLTX18, TLTX23, TLTX25, and TLTX33. A total of 3189 non-cancer hepatocytes from pediatric adjacent normal liver samples were used as the reference for CNV inference. To reduce false-positive detections, only CNVs that were supported by at least one additional DNA profile of the same patient, either snDNA-seq, WES, or Affymetrix OncoScan, were included in further analyses.

To define copy number states of each cell for detected CNV regions, we used two alternative criteria, in which the first assigned CNVs for more cells, while the second was more restrictive and aimed to eliminate potential contamination between genetic subtypes: (1) The inferCNV package applied a hidden Markov model to predict the CNV states. To reduce false-positive detections, CNVs supported by more than 7 of 10 runs of the hidden Markov model were used to define the CNV patterns for each cell. (2) For copy number gain, the average inferred copy number value across the chromosome regions of interest greater than an upper cutoff was defined as *copy number gain*, while less than a lower cutoff was defined as *copy number neutral*. The lower cutoff was set at the 50% quantile of the inferred copy number value of normal reference cells for each CNV region. For TLTX23, TLTX25, and TLTX33, the upper cutoff was set at the 95% quantile of the inferred copy number value of normal reference cells for each CNV region. For TLTX18, which had more tumor cells profiled, the upper cutoff was set at the 99% quantile. Similarly, for copy number loss, the average inferred copy number value across the chromosome regions of interest less than a lower cutoff was defined as *copy number loss*, while greater than an upper cutoff was defined as *copy number neutral*. The upper cutoff was set at the 50% quantile of the inferred copy number value of normal reference cells for each CNV region. For TLTX23, TLTX25, and TLTX33, the lower cutoff was set at the 5% quantile of the inferred copy number value of normal reference cells for each CNV region. For TLTX18, the lower cutoff was set at the 1% quantile.

### Whole exome and targeted sequencing and analysis

A total of 40 tumor samples, including two biopsies of TLTX23 and TLTX36, and 12 matched normal samples were collected with consent from 12 patients from Texas Children’s Hospital with approval from the institutional review board (H13999). Because only WES profiles of one pre-treatment biopsy and five post-treatment resection regions were available to support the discovery of most mutations in TLTX36, we mandated that each mutation be identified in at least two regions. The exon regions of these 52 DNA samples were captured using Agilent SureSelect and sequenced. Quality control was performed on sequence reads with FastQC^10^, and sequence reads were trimmed with Trimmomatic^11^. To verify key mutation and copy number alterations calls, we used the Texas Children’s Hospital’s Solid Tumor Comprehensive Panel, or a mean coverage of 345x of 124 genes, as previously described^12^. Somatic variant discovery was performed with the Genome Analysis Toolkit (v4.1.9.0)^13^. The quality-controlled reads were aligned to the human GRCh38 reference genome^14^ with the BWA-MEM algorithm in the Burrows-Wheeler Aligner^15^ using options --TRAILING 20, --MINLEN 25, and PE, and the BAM files were indexed using Samtools^16^.

To detect somatic variants, the two modes of Mutect2^17^, single tumor sample and jointly multiple tumor samples, with the panel of 12 normal samples, were used to call somatic variants. Single-nucleotide variants (SNVs) and small insertions and deletions (indels) were filtered with PASS tags by GATK FilterMutectCalls with default parameters and subsequently annotated by ANNOVAR (October 4, 2019, release) with refGene reference^18^. The results of these 2 modes were intersected to extract the final SNV and INDEL files for subsequent analysis. Mutations in select genes were also called using the Texas Children’s Pediatric Solid Tumor Cancer Mutation Panel.

To detect CNVs in WES data, FACETS (v0.6.2)^19^ was used to calculate the total copy number, minor copy number, and cancer purity based on SNV read counts from tumor-normal BAM files, with GRCh38p7_GATK_00-common_all as the reference. The cellular fraction (CF) of CNV (copy number alteration) estimated by FACETS can be transformed into a cancer cell fraction (CCF) using the formula CCF = CF/p, where p denotes tumor purity^20^.

### Inferring CNVs by Affymetrix OncoScan

Genome-wide copy numbers were estimated using DNA isolated from formalin-fixed paraffin-embedded (FFPE) tumor tissue using the Affymetrix OncoScan FFPE Assay Kit and OncoScan Console 1.3, and data were analyzed by Affymetrix ChAS 3.1 and OncoScan Nexus Copy Number 7.5 after Burrows-Wheeler Aligner alignment to GRCh38, as previously reported^21^. Affymetrix OncoScan detects copy number alterations at a 50-kb resolution and was extensively validated in our laboratory for clinical use.

### RNA expression profiling by NanoString

We profiled the gene expression of each FFPE tissue region using a custom NanoString nCounter panel (Table S21), partially based on the PanCancer Pathways Panel and HB-specific genes (Table S22), as previously reported^21^. HB-specific genes were selected based on differential expression and risk-predictive potential observed in previously reported HB expression profiling efforts and to represent key hepatocarcinogenesis pathways. Multiplexed measurements of gene expression through digital readouts of the relative abundance of mRNA transcripts were performed as follows: Hybridization of RNA to fluorescent reporter probes and capture probes, purification of the target/probe complexes using nCounter SPRINT cartridges, and imaging and analysis using nCounter SPRINT Profiler. Gene expression profiles across samples were normalized to equate positive-control probes, and expression estimates were z-score-normalized independently for each probe.

### HB17 Culturing and drug-response assays

The HB17 patient-derived cell line was cultured in media composed of 50% Minimum Essential Media (MEM) and 50% Hepatocyte Basal Media (HBM, Lonza, Walkerville, MD). The cells were incubated at 37°C in an atmosphere containing 5% CO2. Cisplatin and ICG-001 were administered at specified doses and time points to achieve a response with minimal toxicity (Figure S4). Cell viability was evaluated using 3-(4,5-dimethylthiazol-2-yl)-2,5-diphenyltetrazolium bromide (MTT) assays. IC values were determined by GraphPad Prism (GraphPad Software, Inc., La Jolla, CA) employing a nonlinear regression model of log(inhibitor) versus normalized response with a variable slope. RNA was extracted from the treated cells for qRT-PCR experiments to measure the expression levels of *DKK1* and *LIN28B*. We used the well-established β-catenin/TCF reporter system^22^. Comparing the activity profiles of pTOPFLASH and pFOPFLASH reporters in HEPG2 cells—an HBC cell line with high β-catenin activity. The reporter pTOPFLASH contains 3 copies of the β-catenin/TCF binding site CCTTTGATC, while pFOPFLASH contains the mutant binding site CCTTTGGCC. Thus, in HEPG2, pTOPFLASH is highly active while pFOPFLASH shows relatively lower activity. Inhibition of β-catenin activity reduces its differential activity. DKK1 is a well-known inhibitor of WNT-signaling and β-catenin activity. As expected, we found that DKK1 silencing significantly increased β-catenin/TCF activity. In contrast, silencing β-catenin or treating with ICG-001 or WNT-C59 reduced β-catenin/TCF activity. Importantly, co-silencing DKK1 partially rescued inhibition by ICG-001 and WNT-C59. Treated cells were also profiled by scRNA-seq.

### Estimation of cancer cell fractions for scDNA-seq

Variants that met the following criteria were analyzed using PyClone-VI (--num-clusters 20 --density beta-binomial --num-restarts 100)^20,23,24^: (1) The exonic function of mutations is neither a synonymous SNV nor unknown, following the annotations from the GRCh38 reference genome; (2) the coverage of mutations is at least 10 reads in all samples from a patient; and (3) the total copy number of the segment overlapping the mutations is not equal to zero. We ensured that variants with zero copy number were analyzed by adjusting the total copy number input to 2 if the copy number was zero and if the cancer cell fraction (CCF) of the CNV was less than 1.

SNVs with exonic function, i.e., non-synonymous SNV, startloss, stopgain, and stoploss, analyzed in PyClone-VI were converted to browser extensible data (BED) files to generate pileup files for scDNA and to variant call format (VCF) files for scRNA by Samtools mpileup. GRCh38.d1.vd1.fa and refdata-gex-GRCh38-2020-A (10x Genomics) were used in mpileup as references in scDNA and scRNA, respectively. The pileup files of each cell were used to extract reference counts and alternative counts. VCFs were used to check whether WES SNVs were identified in the scRNA BAM files before extracting a single-cell variant. The VarTrix coverage mode from 10x Genomics was used to extract single-cell reference counts and alternative counts for scDNA-seq.

### Reconstructing evolutionary trees from WES and single-cell resolution profiles

TLTX36 phylogeny was reconstructed based on the WES profiles of six tumor regions, a tumor-adjacent region, and a blood sample; TLTX36 A1, included in Figure 5A, is the diagnostic biopsy, while the rest are resection samples. Phylogeny inference by Chimaera followed previously described protocols^24^. For TLTX18 and TLTX25, the phylogenetic trees were reconstructed based on observations from their snDNA-seq profiles (Figures S7A and S7C), assuming that tumor cells evolve continuously and each CNV arises only once. Tumor cells from the snRNA-seq profiles with the corresponding inferred CNV patterns were then assigned to each node in the phylogenetic trees. We used criterion (1) for Inferring CNVs from snRNA-seq when defining CNV patterns. Any cells with contradictory node assignments based on the CNVs from snRNA-seq inference criterion (2) were excluded from further analysis. For TLTX23 and TLTX33, the phylogenetic trees were reconstructed based on inferred CNVs from their snRNA-seq profiles. Using criterion (2) with a more stringent cutoff (upper cutoff at the 99% quantile for copy gain and lower cutoff at the 1% quantile for copy loss), we tested the sequence of CNV pairs of interest based on the number of cells supporting which CNV occurred first. If there were significantly more cells with CNV A and without CNV B than cells with CNV B and without CNV A, then CNV A was considered to occur before CNV B (p<0.01, hypergeometric). CNV pairs that could not be ordered were not distinguished and were included in the same node. Cells were assigned to the established nodes based on criterion (2). Finally, predicted ancestral relations between genetic subclones were tested to ensure the criteria were significant and non-overlapping (Figure S8). Mutations inferred from snRNA-seq data supported by at least one additional DNA profile from the same patient—either by snDNA-seq or WES—were mapped to their comprising nodes in our phylogenetic trees.

### HB-HCC subtype classification

We identified HB and HCC biomarker sets using a 2-step process. First, differential expression analysis comparing snRNA-seq profiles of HB and HCC resections identified genes that were differentially expressed between HB and HCC cells. The samples included 2375 HB and 1468 HCC cells that passed quality control (Figure S2A). We identified 1427 differentially expressed genes, including 484 upregulated in HB and 943 upregulated in HCC. Then, to avoid sampling bias and ensure the generalizability of the gene set, we refined the gene set using two independent single-cell datasets. For HB, we compared 1918 HB cells with 1792 adjacent normal hepatocytes using a published scRNA-seq dataset^25^ and identified 45 of the 484 HB genes as upregulated in HB cells relative to adjacent normal hepatocytes. Similarly, a comparison of 1487 HCC cells and 6170 adjacent normal cells from two pediatric AGAR-trial patients with HCC^26^ found that 265 of 943 of the HCC genes were upregulated in HCC cells relative to adjacent normal hepatocytes. The resulting gene sets comprised the HB and HCC biomarker sets. In total, classifier training included two sets of normal (tumor-adjacent) cells derived from AGAR HCC samples (6,170 cells) and eight normal samples corresponding to the HB samples from Song et al. (1,792 cells). It also included snRNA-seq profiles of one HB and one HCC resection from our institution, three HB samples from Song et al., and two AGAR HCC samples. Note that no genes met the criteria of being both significantly downregulated in HCC relative to adjacent-normal and HB. This is largely because variability in normal cell expression reduced the likelihood of identifying genes that were simultaneously downregulated in HCC vs. HB and relative to normal. While twenty-six genes were downregulated in HB vs. HCC and normal samples, we elected not to include this direction. These genes account for <10% of the genes that are differentially expressed in HB vs. HCC and upregulated in cancer.

Cancer cell-type evaluations of profiled cells and nuclei samples were performed by comparing the mean normalized expression of the HB and HCC biomarker sets. For single-cell resolution profiles, raw count matrices were log-normalized with Seurat^3^, and the expression values for HB- and HCC-specific genes were min-max normalized across cells. Raw HB and HCC scores were obtained by summing the normalized expression values of HB- and HCC-specific genes within cells. Finally, raw HB and HCC scores of tumor cells were transformed into z-scores using the means and standard deviations of raw HB and HCC scores of adjacent-normal cells. For the high-risk HB and 5 HBC resection samples, z-score transformations were based on adjacent normal cells in 5 HBCs. For the 2 AGAR-trial HBC samples, z-score transformations were based on adjacent normal cells within these 2 samples.

The cell-type classification criteria were as follows: Cells with an HB z-score greater than 2 and an HCC z-score less than 2 or a z-score difference of at least 2 between HB and HCC were classified as HB cells. Cells with an HB z-score less than 2 and an HCC z-score greater than 2 or a z-score difference of at least 2 between HB and HCC were classified as HCC cells. Cells with both HB and HCC z-scores greater than 2 were classified as HBC cells. Tumor cells that showed no significant upregulation (z-scores less than 2) of either HB or HCC genes were considered unassigned tumor cell types and were excluded from further analysis. The proportion of cells excluded in each sample was as follows. TLTX01: 25.7%; TLTX11: 4.3%; TLTX18: 2.6%; TLTX23: 4.0%; TLTX25: 0.5%; TLTX33: 5.9%; AGAR HBC: 4.1%. The visualization in Figure 2A shows evaluations of random samples (100 cells). When analyzing bulk profiles, evaluations were performed based on the available biomarkers—5 HB- and 8 HCC-specific genes by NanoString nCounter and 42 HB- and 245 HCC-specific genes by bulk RNA-seq. The expression values for HB- and HCC-specific genes were min-max normalized across samples. Raw HB and HCC scores were obtained by summing the normalized expression values of HB- and HCC-specific genes within each sample. Like the methods described for single-cell resolution profiles, the raw scores were transformed into z-scores based on the means and standard deviations of the raw scores of the normal samples.

### Liver cell embryonic differentiation inference

We trained a classifier to predict embryonic liver differentiation stages using scRNA-seq profiles of human embryonic and fetal liver samples^27^. This dataset includes profiles of nine developing liver samples taken from embryos from 5 weeks (5W) to 19 weeks post-conception (19W) (Figure S3A). Cell-type annotation followed the outline provided by Wang et al^27^. We included 19,557 developing liver cells from the hepatoblast-hepatocyte lineage. We trained a random forest model to distinguish between 5W (early differentiation) and 19W (late differentiation) cells, selecting 20 tree-classifiers based on performance in training: Increasing the number of classifiers beyond 20 failed to result in significant training error improvement (Figure S3D). A total of 811 differentially expressed genes were used as predictive features. Here, given a profile, each classification tree produces a binary decision, <DIFFERENTIATED> or <UNDIFFERENTIATED>, and the differentiation score is the normalized count of <UNDIFFERENTIATED> calls, from 0 to 1. Namely, the predicted differentiation stage was quantified by a differentiation score as the proportion of <UNDIFFERENTIATED> votes among the 20 trees. To compare the proportion of undifferentiated cells in each tumor, we set a differentiation score cutoff based on its 95% quantile in profiles of 11,480 normal hepatocyte cells from a tumor-adjacent sample, producing a differentiation score cutoff of 0.4 (Figure S3E). To address modality differences and improve the generalizability of the trained model on bulk RNA profiles, each profile was transformed into ranks, and the classifier evaluated gene ranks in each sample, pseudobulk, or cell profiles independently. Cells from intermediate time points (6W-16W) were used to evaluate classification quality, and independent testing was performed on an embryonic mouse liver scRNA-seq dataset derived from samples taken from 11 (E11) to 17.5 (E17.5) embryonic days post-conception with 12,393 developing liver cells from the hepatoblast-hepatocyte lineage by Wang et al^27^. The results suggest significant quantitative predictive ability for human liver cells (Figure S3B, p=8E−3, hypergeometric distribution) and mouse liver cells (Figure S3C, p=7E−3, hypergeometric distribution).

### Cisplatin-resistance prediction

Hirsh et al. profiled RNA and evaluated cisplatin resistance in nine pediatric liver cancer cell lines^28^. We identified 37 genes whose expression profiles were individually significantly predictive of cisplatin resistance in these lines (0.01% quantile by rank). This set included 20 genes whose expression correlated with cisplatin resistance (*AGPAT3*, *BRPF3*, *DENND5B*, *DTX2*, *FBXO6*, *FOXA1*, *GRTP1*, *IQCH-AS1*, *KLHL9*, *LINC00235*, *NFATC2*, *NFX1*, *PIK3C2A*, *RNPEP*, *SERPINB1*, *TAF8*, *THRB*, *UBAP1*, *UHRF2*, *ZNF417*), and 17 genes whose expression was negatively correlated with cisplatin resistance (*ANKRD34A*, *APEX2*, *ASB4*, *BMERB1*, *CA11*, *CAMTA1-IT1*, *CCDC74B*, *CERCAM*, *FAM216A*, *HMOX2*, *IL27RA*, *MAP4K1*, *MCTS1*, *METTL27*, *PRICKLE3*, *TBL3*, *UBE2A*). We computed the cisplatin-resistance score as the ratio between the mean expression profiles of genes in the two sets. When evaluating snRNA-seq and scRNA-seq data, the score was calculated based on pseudobulk transformation.

We evaluated our cisplatin-resistance score on the training data (Figure S6A, p=2E−4), on four HB organoids profiled and treated by Song et al.^25^ (Figure S6B, p<0.04, hypergeometric distribution), on low- and high-risk HBs^6^ (Figure S6C, p<0.01), on our reference HB and HCC samples (Figure S6D), and on an HB PDX before and after treatment with cisplatin (Figure S6E). When evaluating the cisplatin-resistance score for our HBC clusters, each cluster shown in Figure 2C for each patient was evaluated independently. The resistance scores were then normalized to ensure comparability by calculating the ratio of each resistance score to the resistance score of the corresponding patient’s normal cells.

### Evaluation of HBC proliferation vs. quiescence

We estimated the proliferative activity of HBC tumor cells across differentiation stages using cell cycle S-related genes identified by Whitfield et al^29^. To produce Figure S3I, HBC cells were categorized into early, medium, and late differentiation stages based on differentiation score cutoffs at 0.2 and 0.45. A proliferation score was calculated for each differentiation stage in the five patients with HBC by evaluating the mean of the pseudobulk expression values across the cell cycle S-related gene set and adjusting for the number of cells in each differentiation stage for each patient. To ensure comparability among the five patients with HBC, each proliferation score was converted to a ratio relative to the proliferation score of the early differentiation stage within each patient.

### Differential gene expression analysis

When looking for differentially expressed genes—including during classifier construction—we used the “FindMarkers” function in Seurat, including genes that were detected in a minimum fraction of 25% of the cells in either of the two tested populations. Genes with a log2-fold change of greater than 0.5 and an adjusted p-value of less than 0.01 (Bonferroni correction) were considered significantly expressed.

### Gene set enrichment analysis

We performed gene set enrichment analysis (GSEA) to calculate pathway enrichment in single-cell clusters using the R package fgsea (v1.24.0)^30^. Gene sets were ranked based on fold change between the cell clusters so they could be compared, and the number of permutations was set at 10,000. Gene sets for pathways were obtained from the “C2” collection of the MSigDB database. In Figure 2G, GSEA was used to compare each HBC tumor cluster and adjacent normal cells. For visualization, pathway scores were z-score transformed across the 10 HBC tumor clusters. Figure 3G shows the −log_10_(adjusted p) values, with positive and negative signs determined by upregulation or downregulation relative to adjacent normal cells.

### Estimating miRNA activity and data processing inequality from snRNA-seq data

We used miReact^31^ based on predicted miRNA targets^32,33^ to estimate *let-7*-family^34^ miRNA activity in cells with more than 1000 mapped unique molecular identifiers. Normalized mutual information (MI) was calculated between (i) estimated *let-7* activity and LIN28B expression, (ii) cell differentiation scores and LIN28B expression levels, and (iii) cell differentiation scores and estimated *let-7*-family miRNA activity. To evaluate MI significance and normalize MI, the expression values of LIN28B, estimated *let-7* activity, and cell differentiation scores were shuffled 1000 times, and the normalized MI calculated from the shuffled triplets was used to fit a generalized extreme value distribution. Furthermore, the differences were calculated between the smallest normalized MI values: One between *let-7* activity and LIN28B, and the other between cell differentiation scores and LIN28B expression levels, compared with the normalized MI between *let-7*activity and cell differentiation scores; p-values were estimated using a generalized extreme value distribution fit to the differences from shuffled data. The normalized MI was also calculated between (i) the expression levels of *let-7* targets and LIN28B, (ii) cell differentiation scores and LIN28B expression levels, and (iii) cell differentiation scores and the expression levels of let-7 targets. The same analysis was conducted for differentiation-related non-*let-7* target genes. A chi-squared test was used to examine whether the distribution of triplet pairs having the smallest normalized MI between cell differentiation scores and target genes differed significantly from that of differentiation-related non-*let-7* target genes. To evaluate the data processing inequality, the 3 MI values were compared across 223 verified *let-7*-family targets. For all but one target, the MI between the cell differentiation scores and the estimated *let-7*-family miRNA activity was the smallest of the three. Assuming that each comparison is as likely to be greater than the other, we can conclude that the relationship between cell differentiation scores and *let-7*-family miRNA activity is indirect at p<1E−13 (binomial distribution).

---

**Figure S1.**
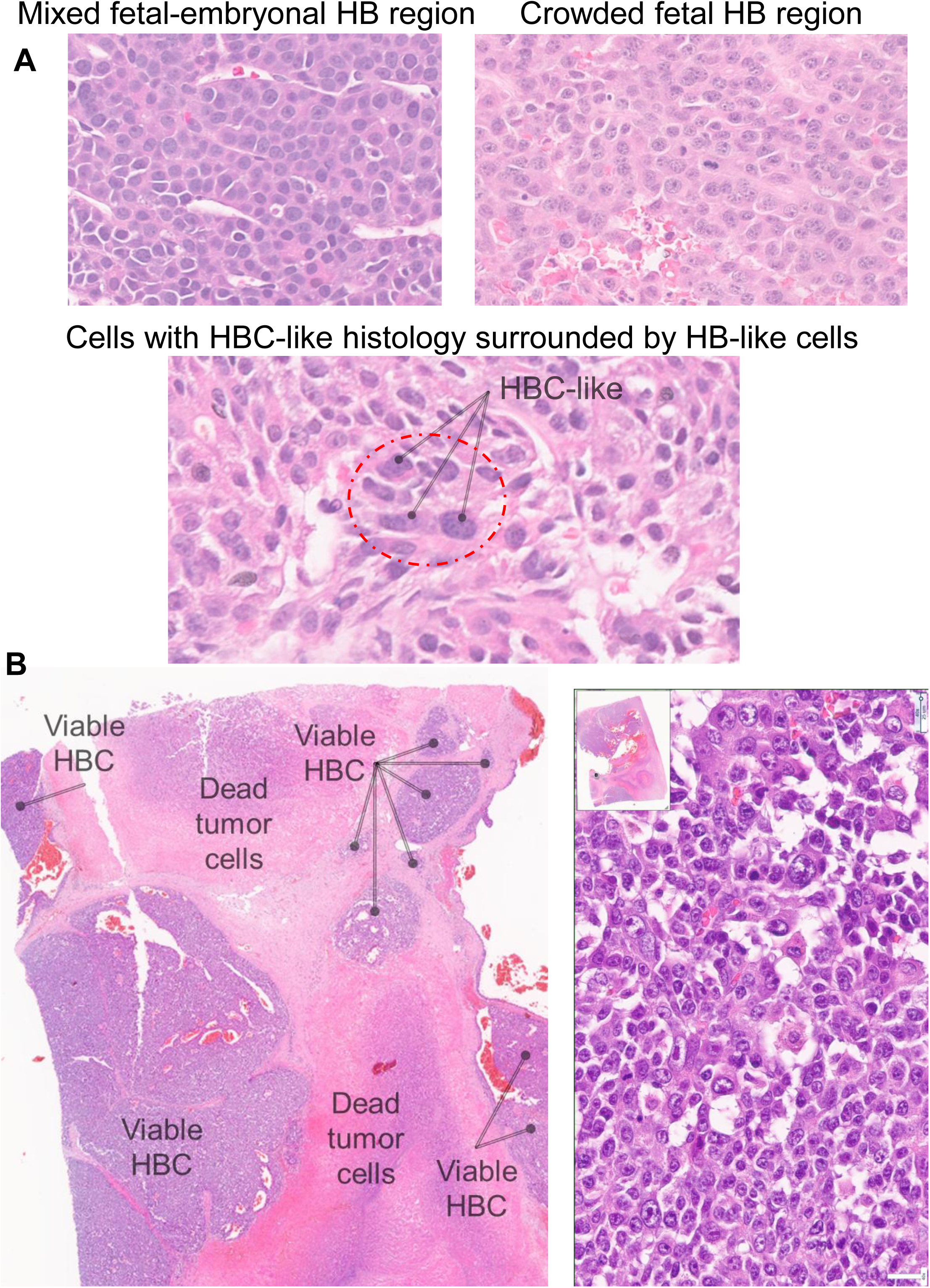
Representative histopathology of pre- and post-treatment HBC. Pre-treatment HBCs are composed of HB, HBC, and HCC cells. (**A**) Multiple regions of the pre-treatment biopsy of the equivocal HBC TLTX16 reveal cells with HB-like histology at variable embryonic differentiation stages (embryonal and fetal) and cells with HBC-like histology that are enveloped in regions that mainly comprise cells with HB-like histology. (**B**) The post-treatment resection (4 cycles of chemotherapy, including cisplatin and vincristine-irinotecan) of TLTX16 reveals extensive necrotic regions adjacent to areas primarily containing viable cells with HBC-like histology.

**Figure S2.**
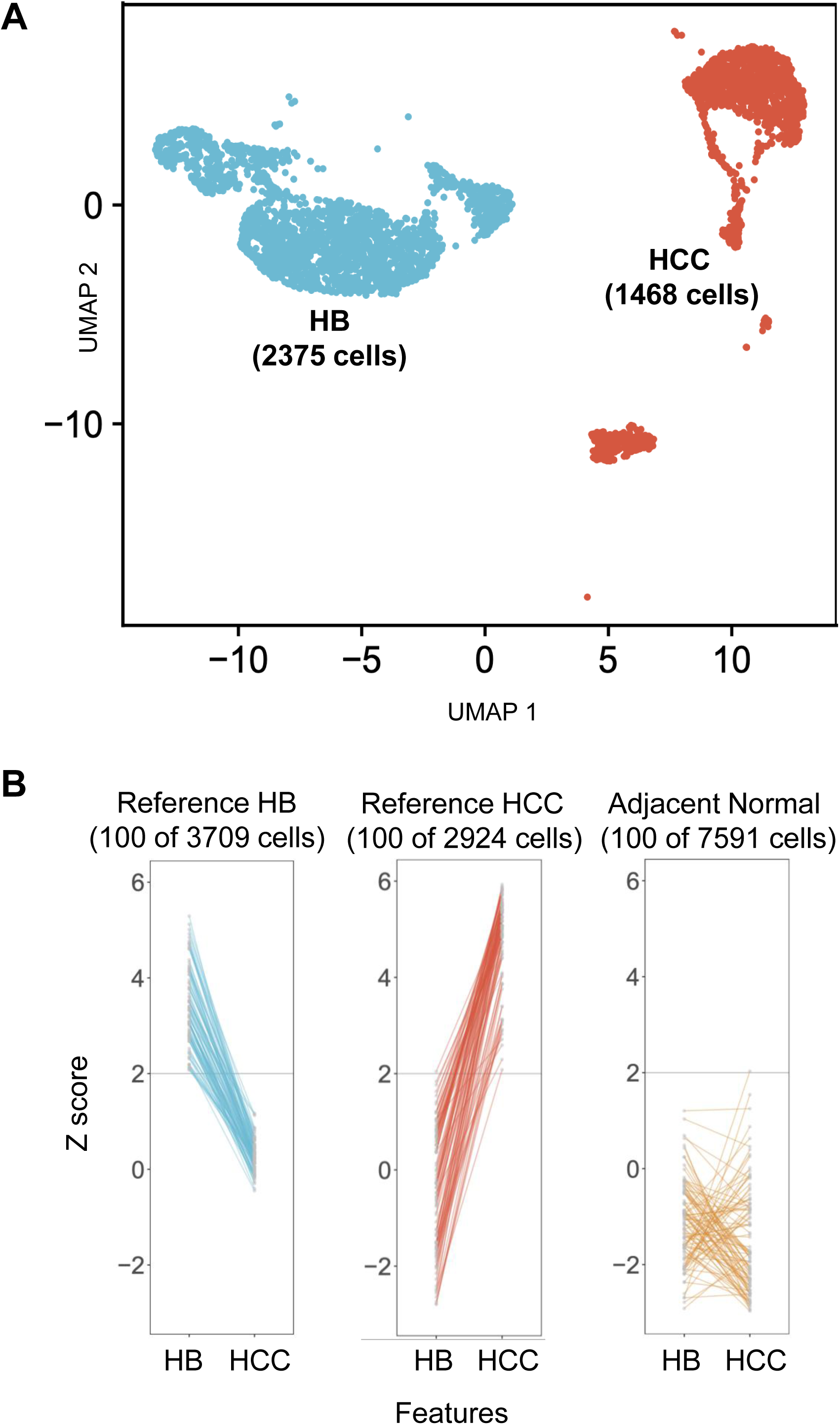
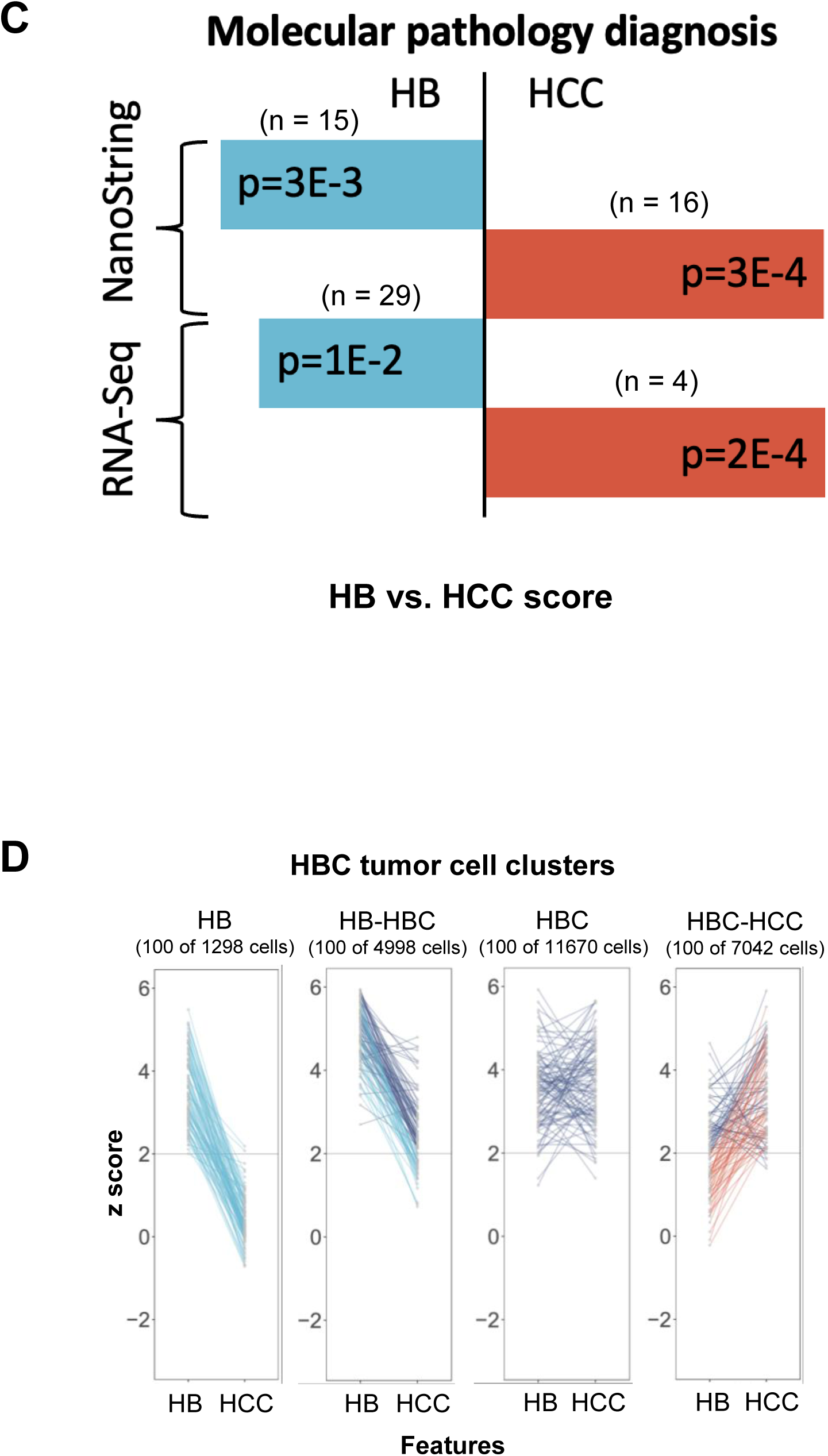
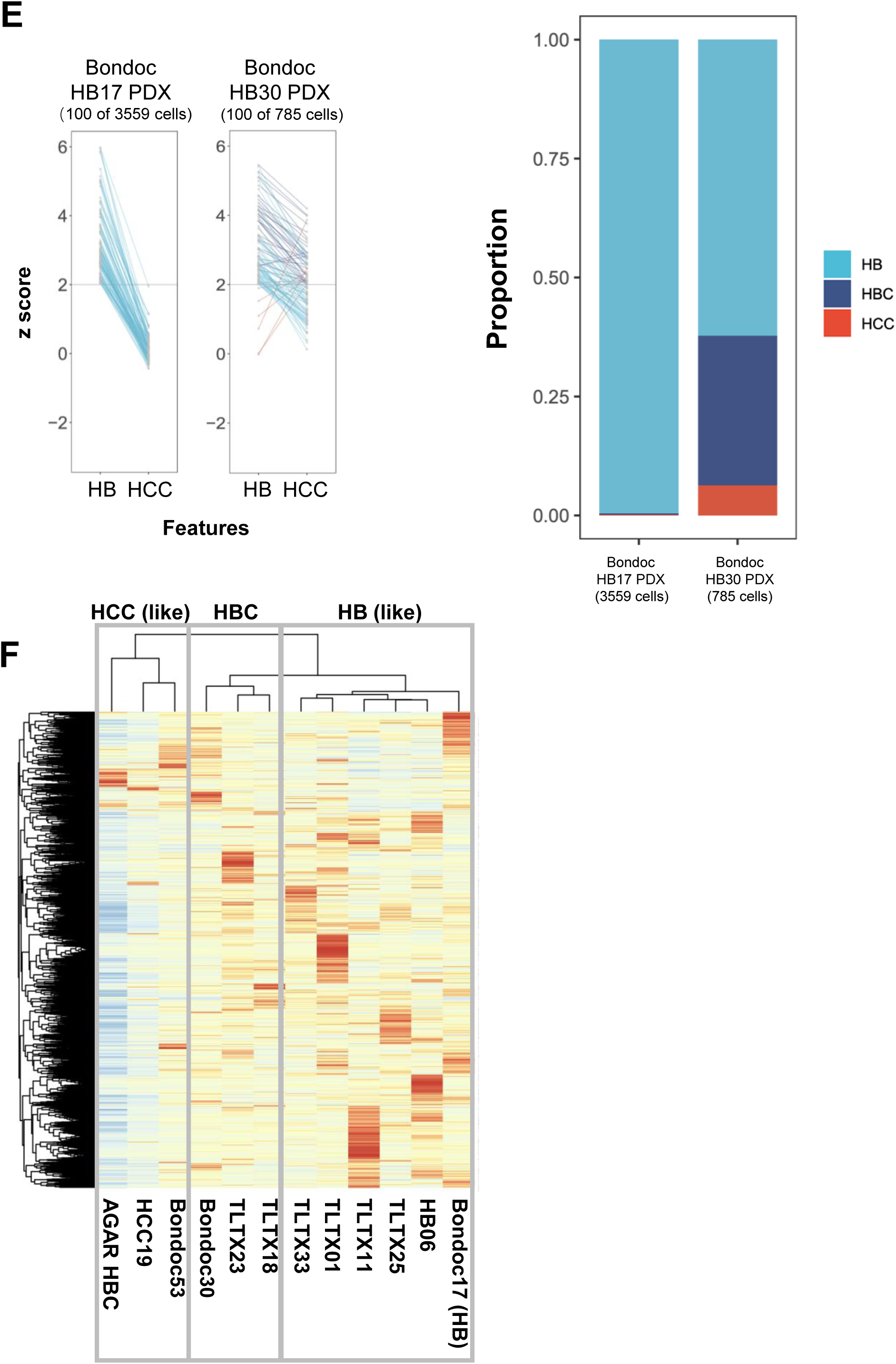
Evaluation of HB and HCC biomarker sets. Our HB and HCC biomarker sets were derived from single-cell resolution profiles of HBs and HCCs, as reported by Song et al. (2022), samples from the AGAR trial by Steffin et al. (2024), and additional profiles, encompassing 2375 HB and 1468 HCC cells, respectively. We refer to these profiles—including snRNA-seq profiles of 2375 HB and 1468 HCC cells—as our reference profiles. (**A**) The UMAP of our reference snRNA-seq profiles of one HB and one HCC demonstrates a perfect separation between HB (light blue) and HCC (red) cells using unbiased clustering. (**B**) HB and HCC biomarker z-scores of cells represented in our reference profiles from our HB and HCC samples, Song et al.—HB cells from patients 1,4, and 6, AGAR-trial HCCs; adjacent normal cells shown (orange) were derived from our reference HB and HCC, and in our 5 HBC samples—these samples were not used during method training. Z-scores greater than 2 indicated significant upregulation of HB or HCC biomarker gene sets. (**C**) HB and HCC biomarker gene set z-scores using bulk RNA-seq profiles of 29 HBs, 4 HCCs, and 6 fetal samples from patients without cancer, alongside NanoString nCounter profiles of 15 HB, 16 HCC, and 9 fetal samples from patients without cancer. The NanoString panel includes only 5 of 45 HB and 8 of 265 HCC biomarker genes; p-values denote the significance of the separation between HB and HCC z-scores. (**D**) We classified HBC-profile clusters—in Figure 2C— into HBs, HB-HBCs, HBCs (purple), and HBC-HCCs. Random samples from each classification are displayed here. Namely, HB clusters predominantly contain cells with z-scores greater than 2 for HB biomarkers and less than 2 for HCC biomarkers. HBC clusters contain cells with z-scores greater than 2 for both HB and HCC biomarkers. HB-HBC and HBC-HCC clusters contain mixtures of cell types. Z-scores greater than 2 indicate significant upregulation of biomarker genes. (**E**) HB and HCC biomarker z-scores and predicted cell type proportions of cells from 2 PDX profiled by Bondoc et al (2021). (**F**) Unsupervised hierarchical clustering of 1 HCC, 8 HBC, and 3 HB pseudobulk samples.

**Figure S3.**
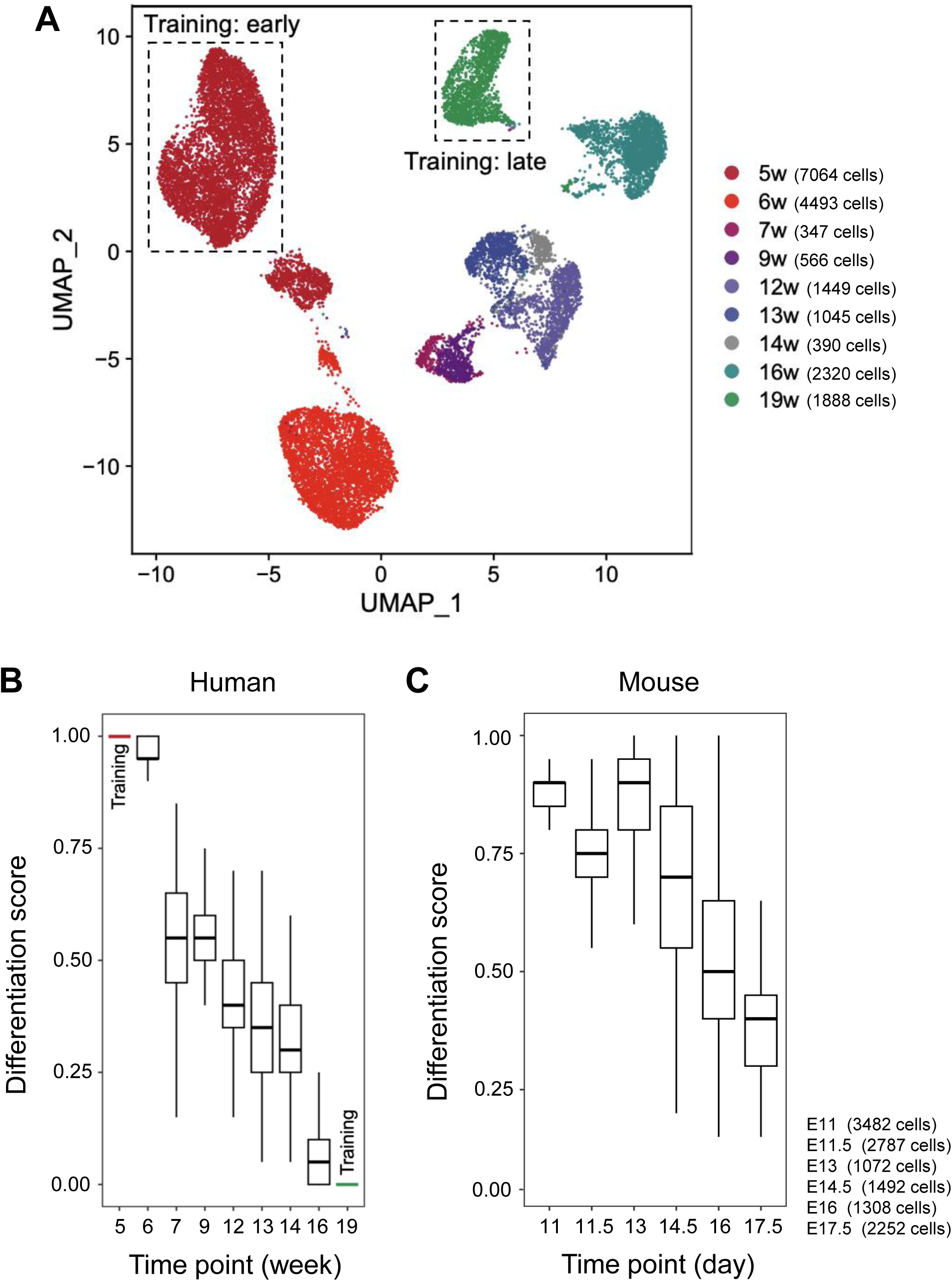
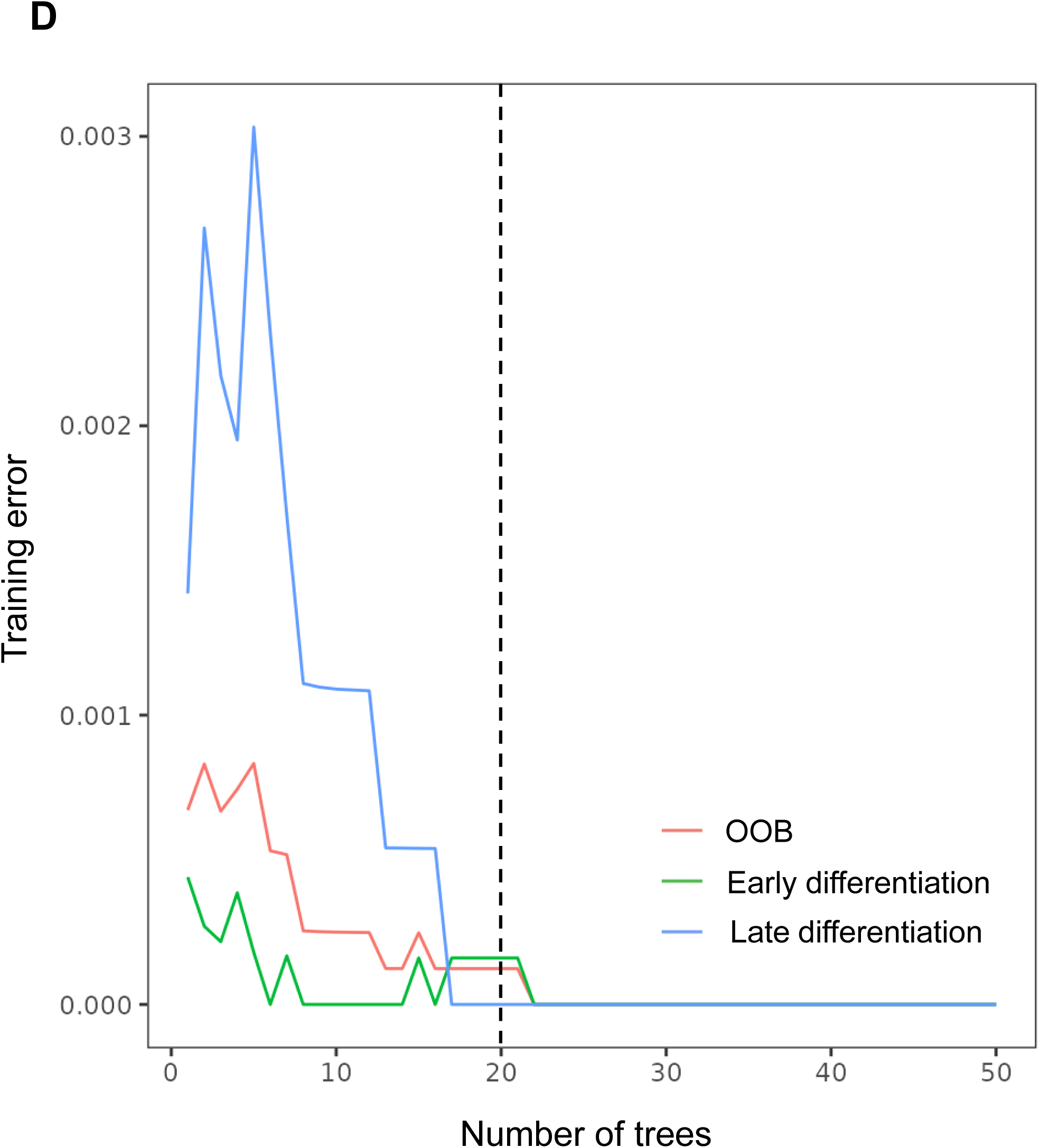
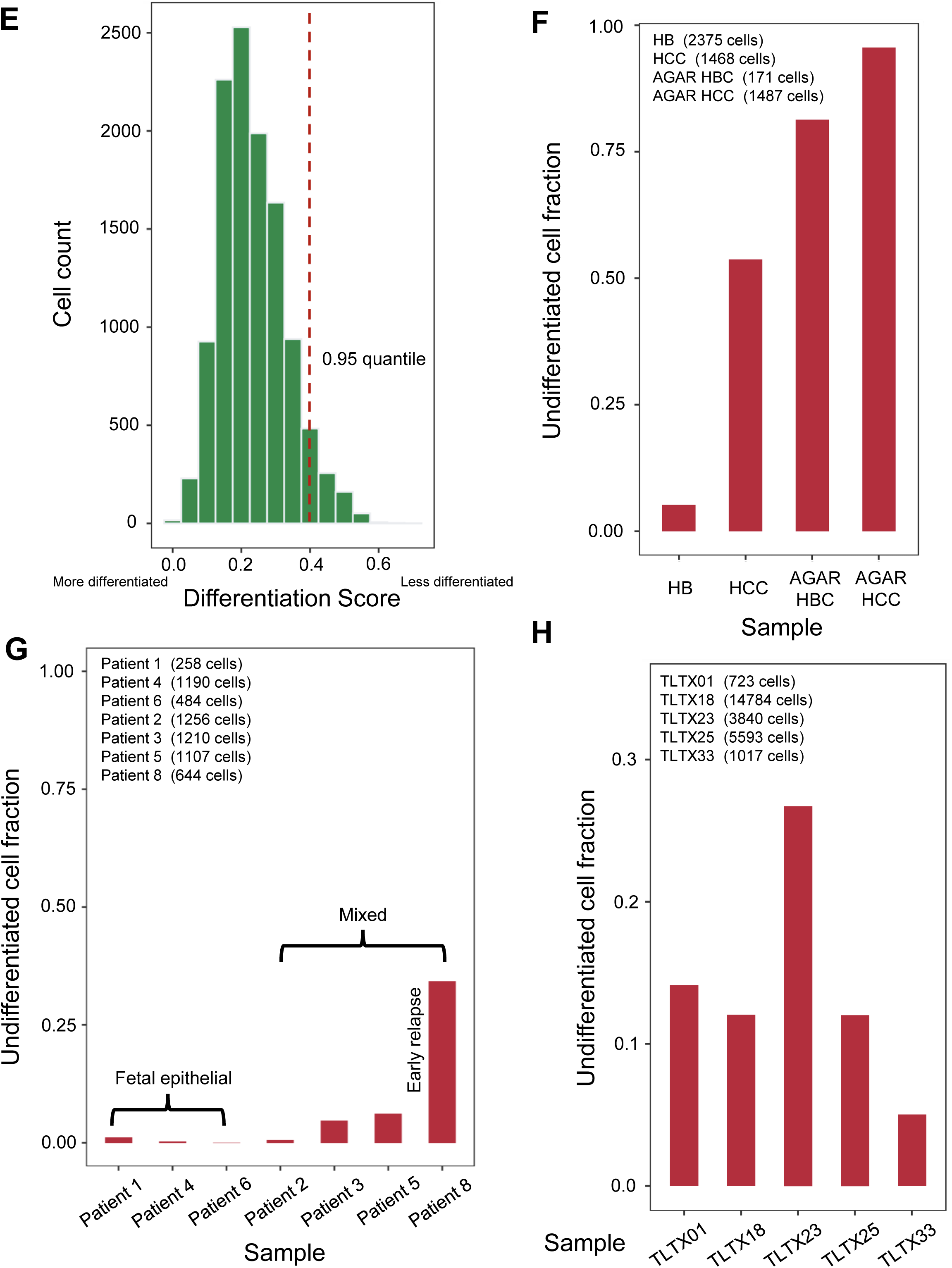
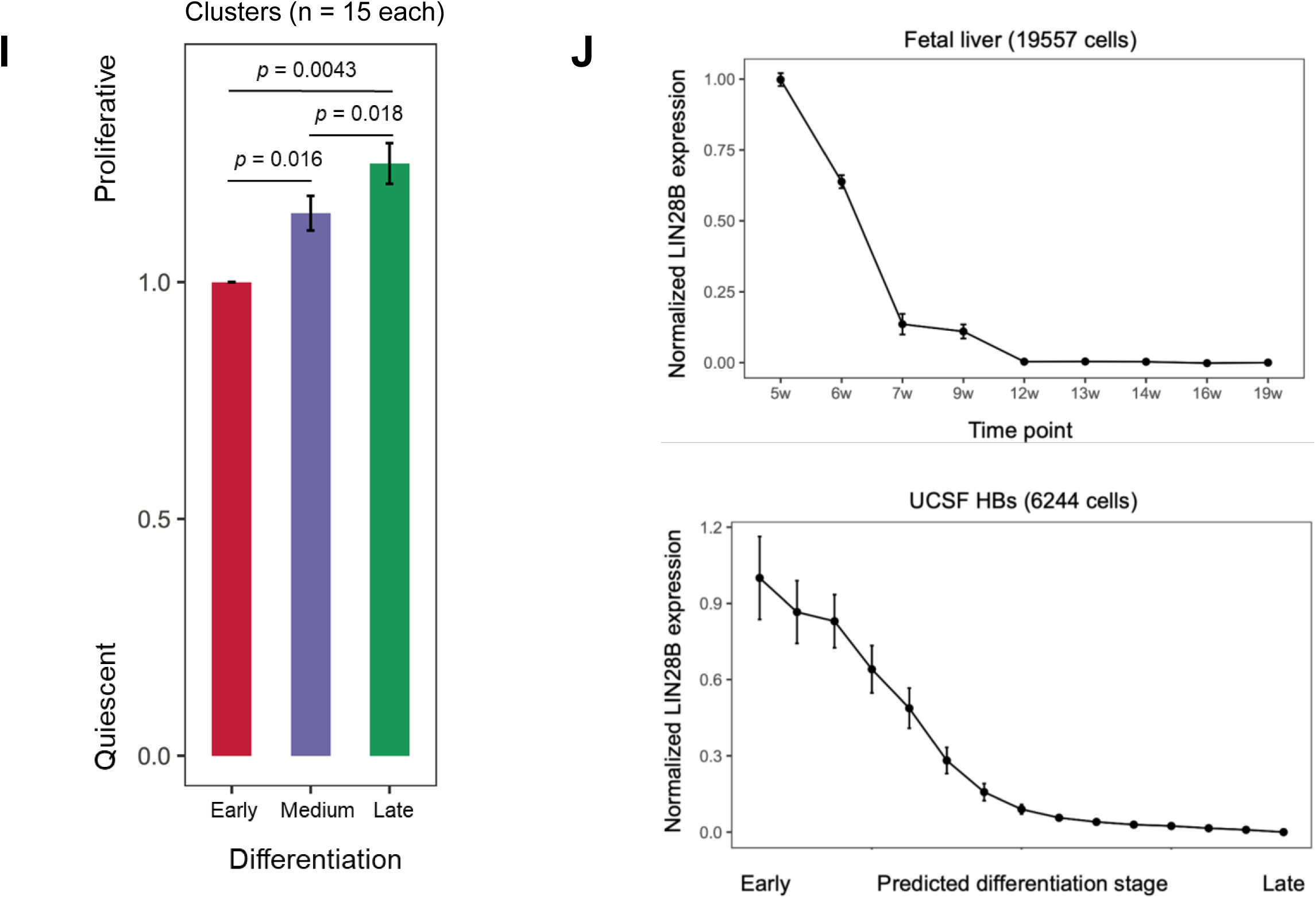
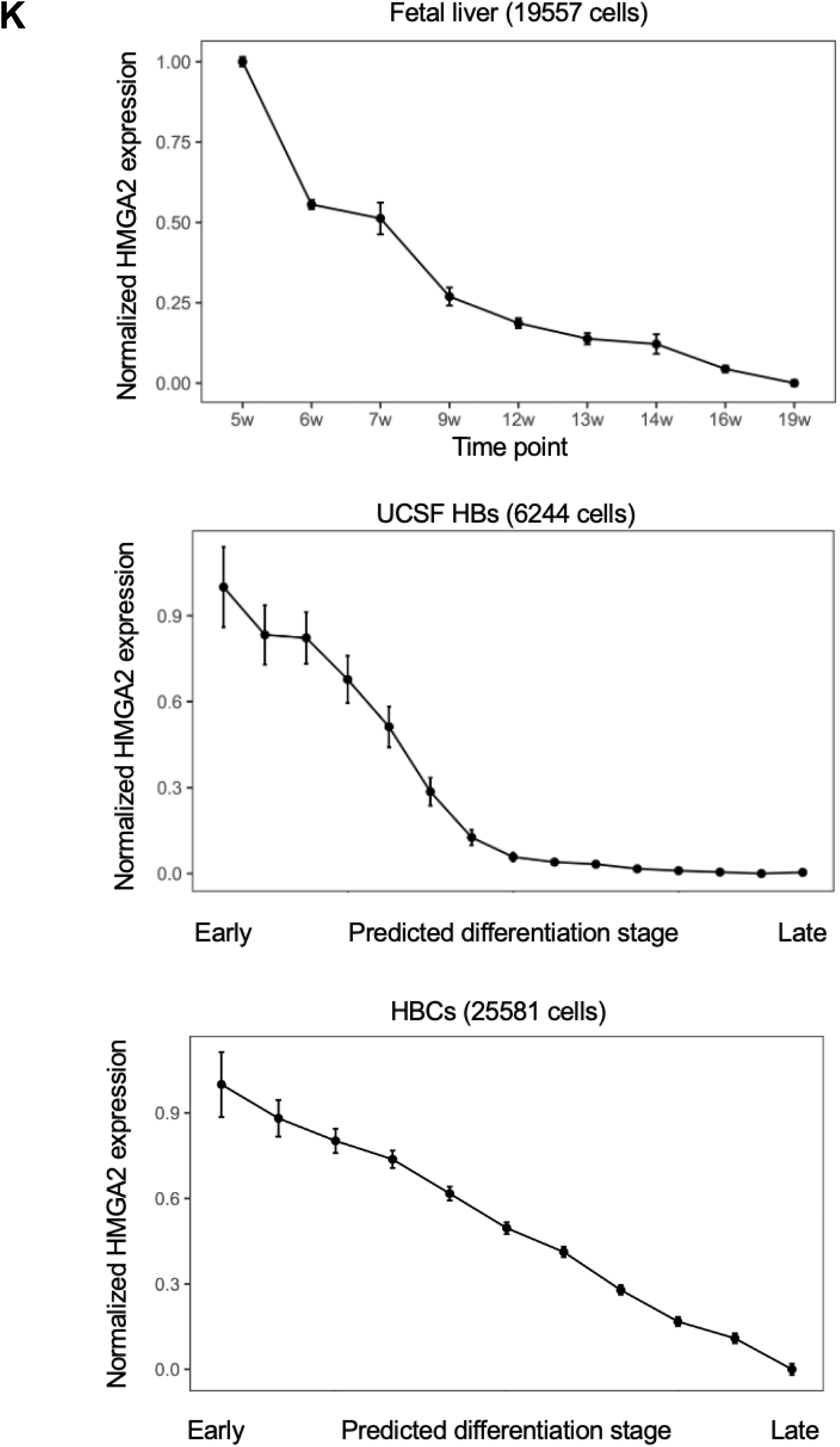
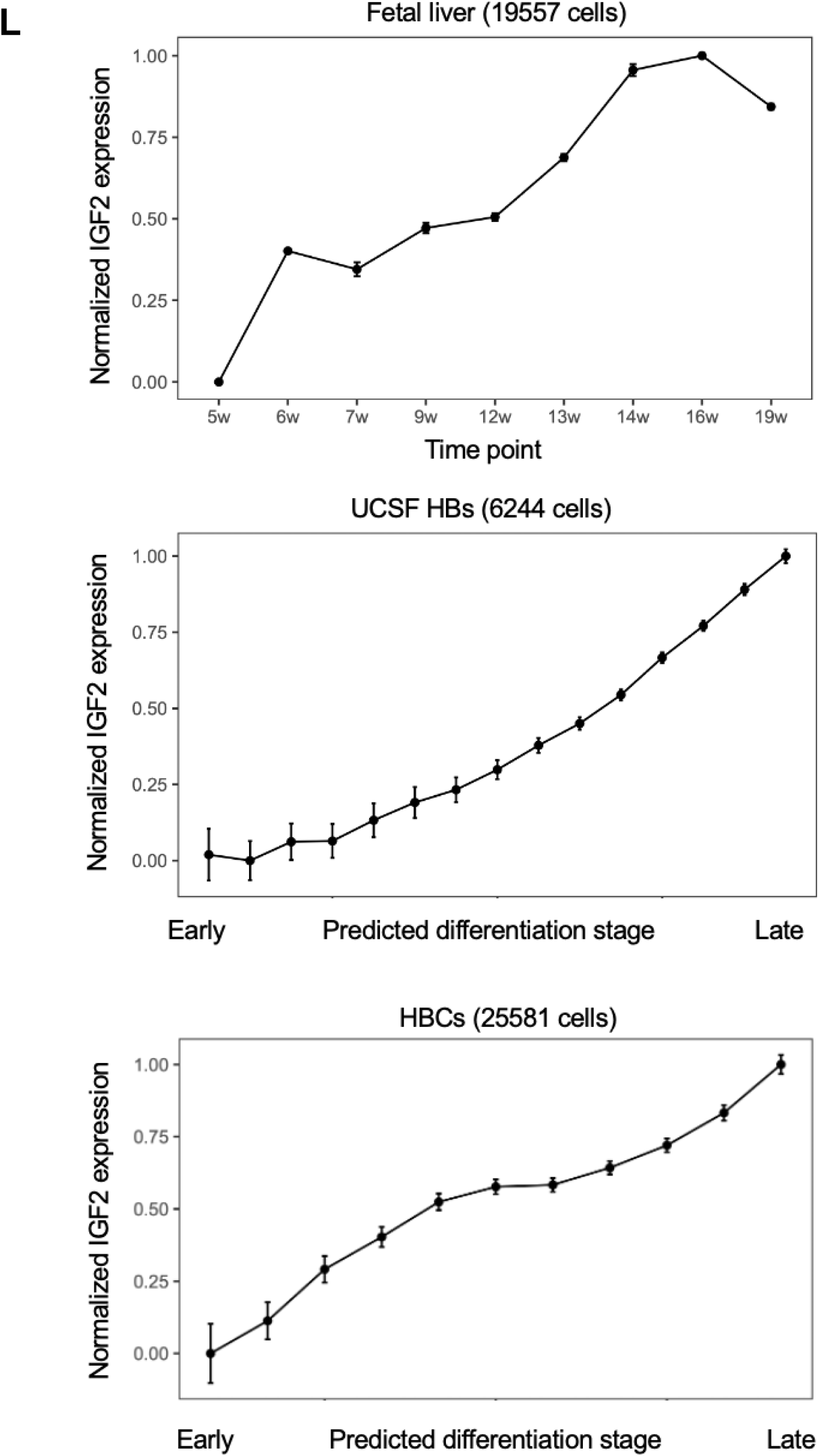
Liver cell differentiation inference. We used profiles of embryonic liver cells by Wang et al. (2020), spanning 5 to 19 weeks post-conception, to infer liver cell differentiation. Specifically, we developed a classifier based on profiles from cells at weeks 5 and 19 post-conception, while profiles of cells taken from embryos 6 to 16 weeks post-conception were used for evaluation. The classifier produces a differentiation score for each cell profile. (**A**) UMAP of 19,557 developing human liver cells from 5 to 19 weeks post-conception. (**B**) Differentiation scores for human embryonic liver cells from 5 to 19 weeks post-conception. (**C**) Differentiation scores of mouse embryonic liver cells from 11 to 17.5 days post-conception. (**D**) Differentiation inference was based on a random forest differentiation prediction classifier that could be trained with any given number of classification trees. We opted for a 20-tree classifier because increasing the number of trees to more than 20 did not significantly reduce the training error. Here, we show the training error as a function of the number of trees; OOB, out-of-bag error. (**E**) To compare the proportion of embryonically undifferentiated cells in each tumor, we set a differentiation score cutoff based on its 95% quantile in profiles of 11,480 normal hepatocyte cells from a tumor-adjacent sample, producing a differentiation score cutoff of 0.4; higher scores indicate undifferentiated embryonic cell stages, while lower scores indicate differentiated stages. (**F**) Inferred undifferentiated cell fractions of our HB reference sample, HCC reference sample, 2 AGAR-trial HBC samples, and 2 AGAR-trial HCC samples. All differences are significant, and the results suggest that these pediatric HCC samples are less embryonically differentiated than HBs, while cancer cells remaining after AGAR-trial treatments are predominantly undifferentiated. (**G**) Inferred embryonically undifferentiated cell fractions of 7 HB samples by Song et al. Higher-risk samples are significantly less embryonically differentiated. (**H**) Inferred undifferentiated cell fractions of 5 HBC samples. TLTX23, characterized by predominant HCC and HBC composition, is less embryonically differentiated. (**I**) Normalized proliferation score of early (differentiation score ≤ 0.2), intermediate (0.2 < differentiation score < 0.45), and late (differentiation score ≥ 0.45) differentiated tumor cells in 5 HBC samples; p-values are based on the paired sample t-test. (**J**) Normalized expression of LIN28B in the human fetal liver during embryonic development (top) and in HBs across predicted embryonic differentiation stages (bottom). Normalized expression profiles of (**K**) HMGA2 and (**L**) IGF2 in human fetal liver development, HBs profiled by Song et al. (2022), and HBCs across predicted embryonic differentiation stages; S.E.M. is shown.

**Figure S4.**
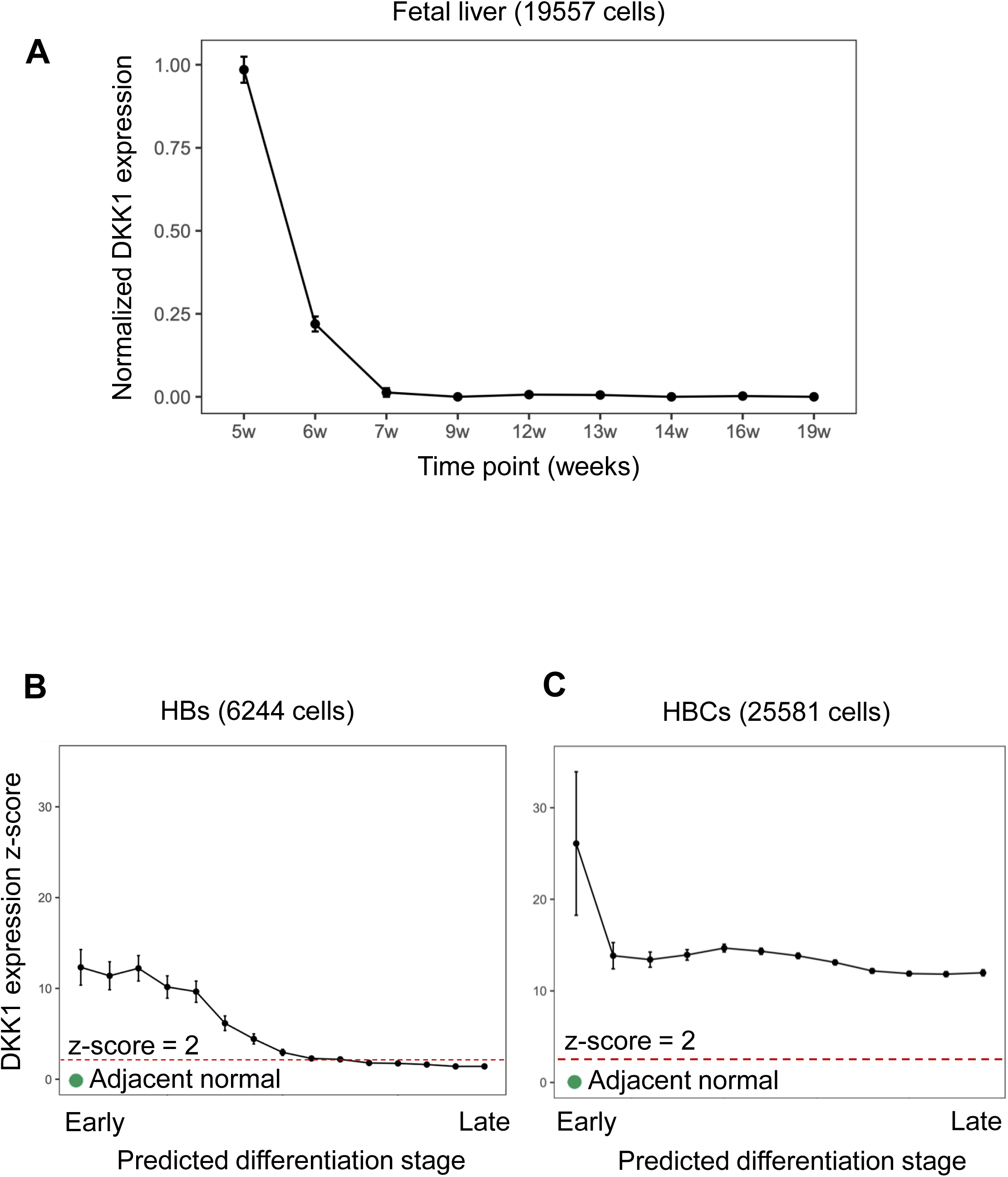
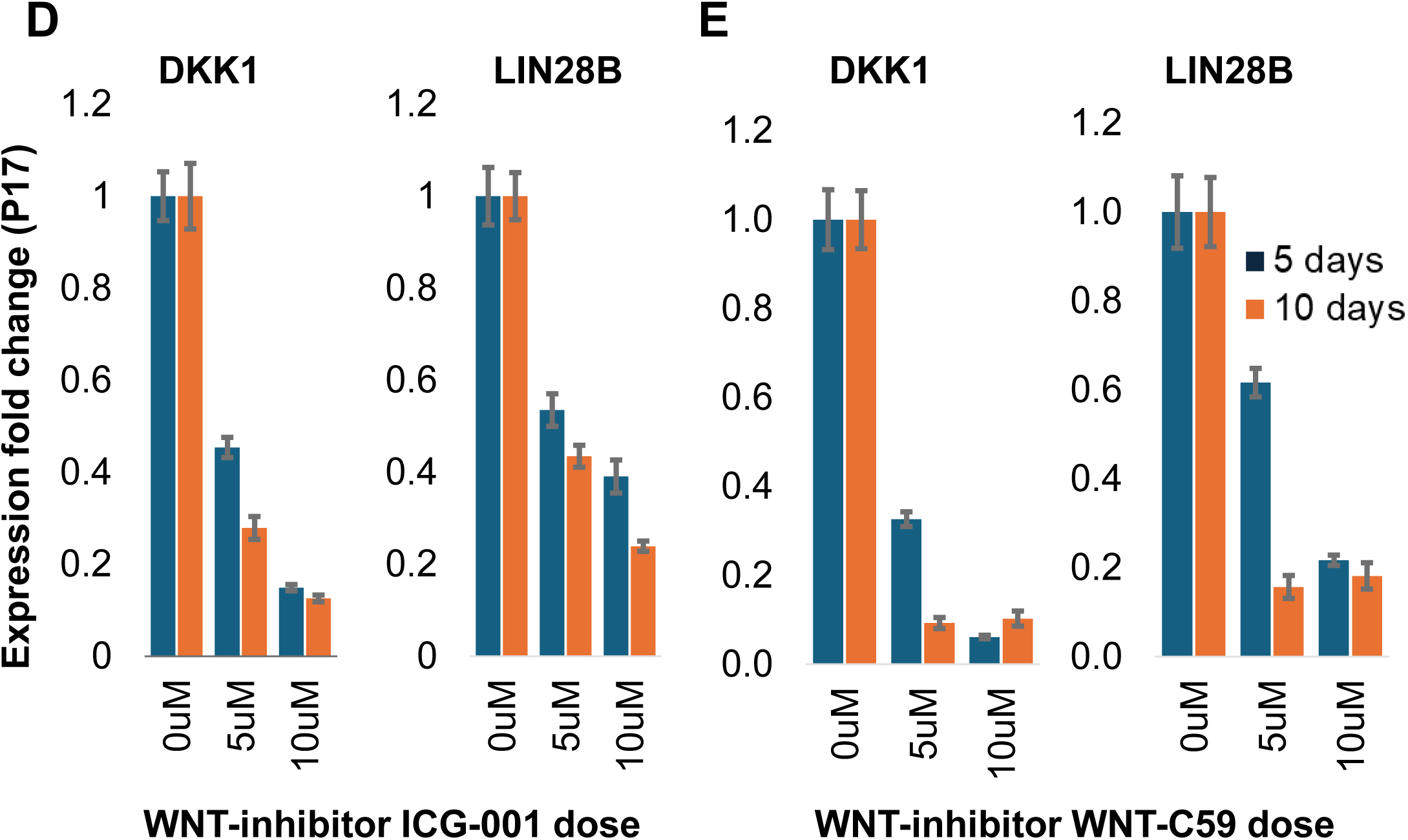
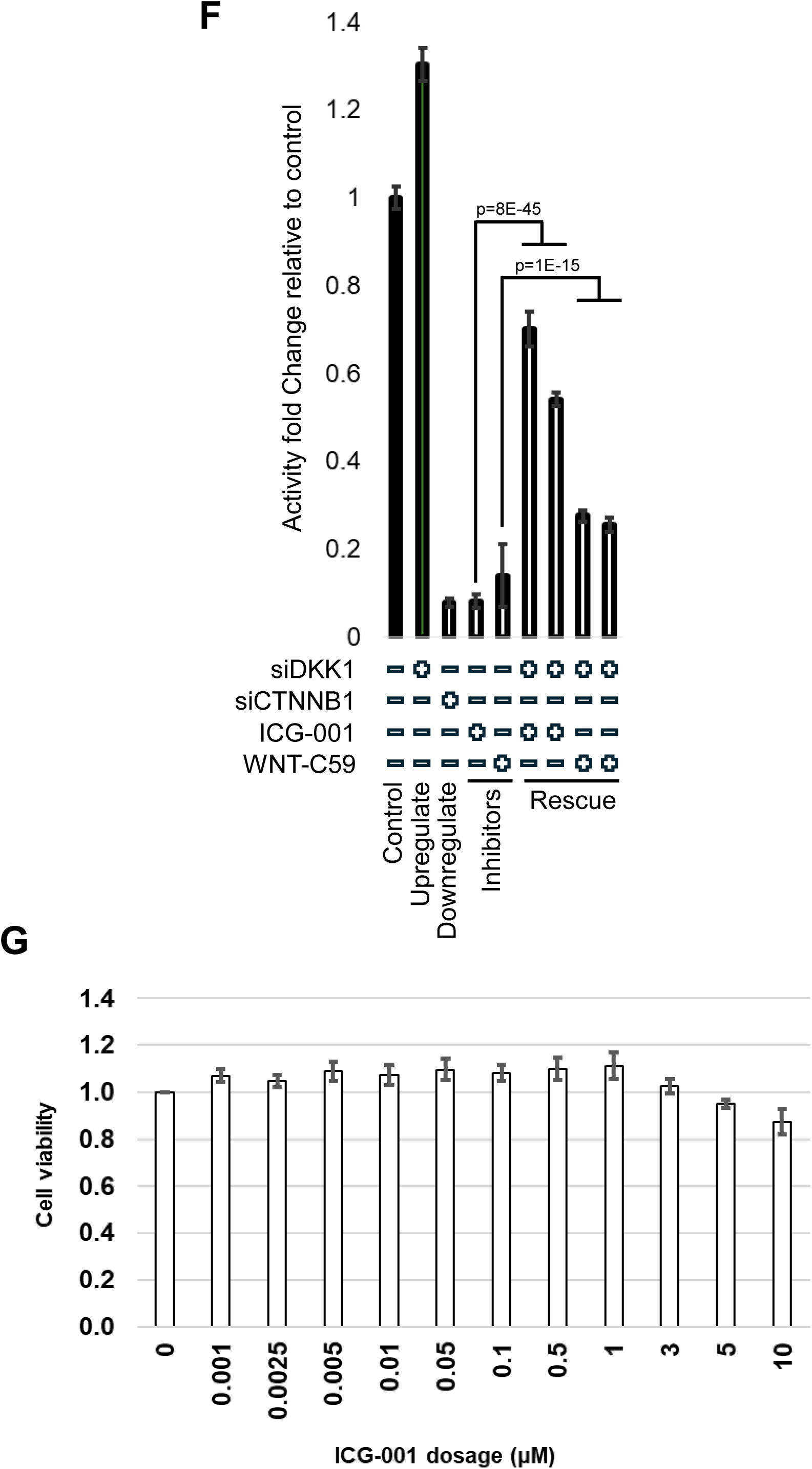
WNT-signaling inhibition decreases DKK1 and LIN28B expression. DKK1 is a widely used indicator of WNT-signaling activity, while LIN28 indicates liver cell differentiation (Sumazin et al., 2017). (**A**) Normalized expression of DKK1 in the human fetal liver during development suggests reduced WNT-signaling activity as a function of liver differentiation; S.E.M. is shown. (**B-C**) Normalized expression of DKK1 in HBs profiled by Song et al. (B) and HBCs (C) across predicted embryonic differentiation stages; S.E.M. is shown. (**D-E**) In vitro treatment with the WNT-inhibitor ICG-001 (D) and WNT-C59 (E) for 5 or 10 days at variable dosages reduced DKK1 and LIN28B expression in the patient-derived cell line HB17; 5 biological replicates, S.E.M. is shown. (F) A comparison of the activity profiles of pTOPFLASH and pFOPFLASH β-catenin/TCF reporters^22^ in HEPG2 cells suggested that inhibition of β-catenin activity by silencing β-catenin with RNAi or treating with the WNT inhibitors ICG-001 or WNT-C59 reduced differential activity, siDKK1 increased β-catenin/TCF activity, and co-silencing DKK1 partially rescued inhibition by ICG-001 and WNT-C59. (G) HB17 cell viability at variable dosages of ICG-001.

**Figure S5.**
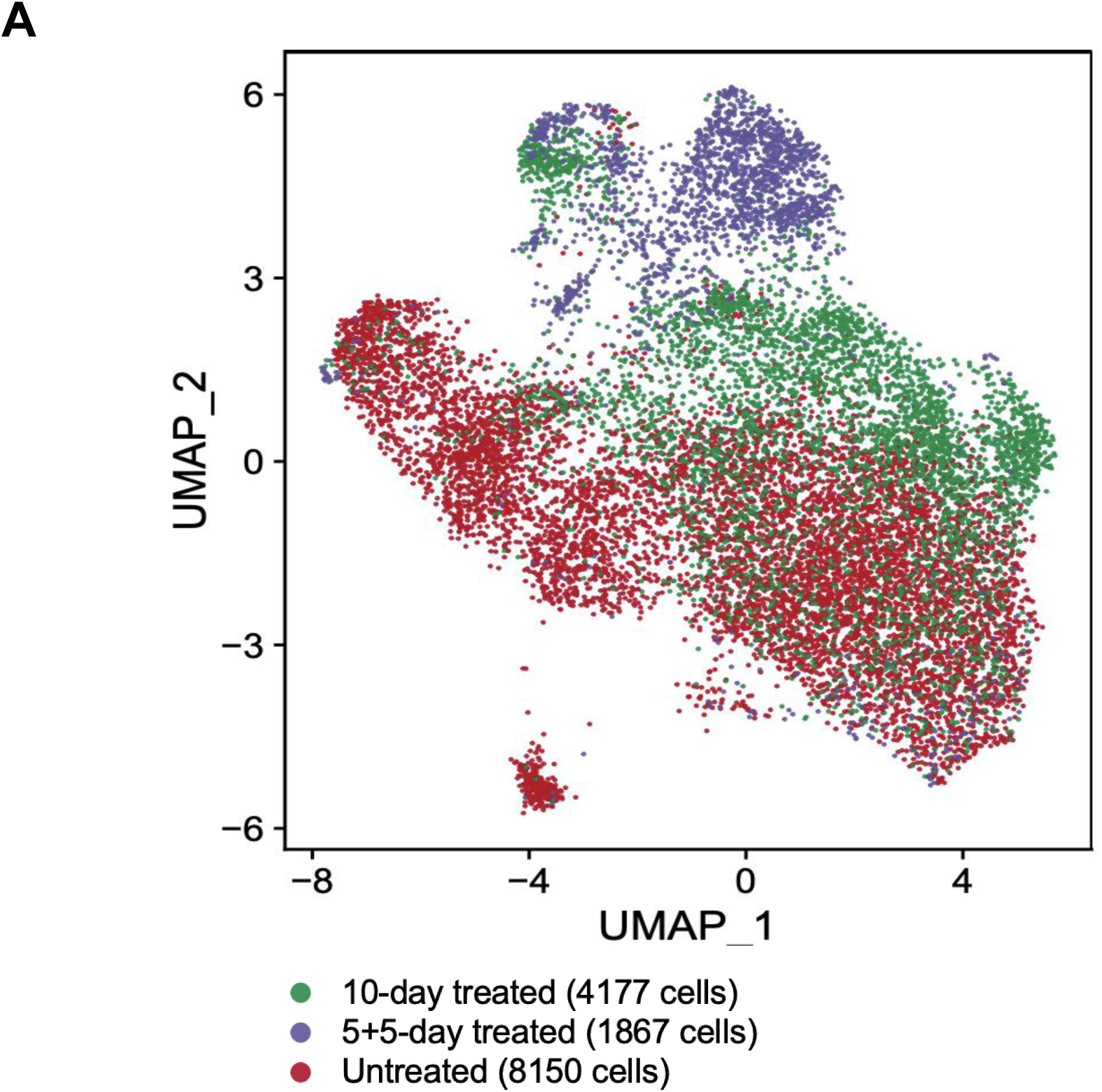
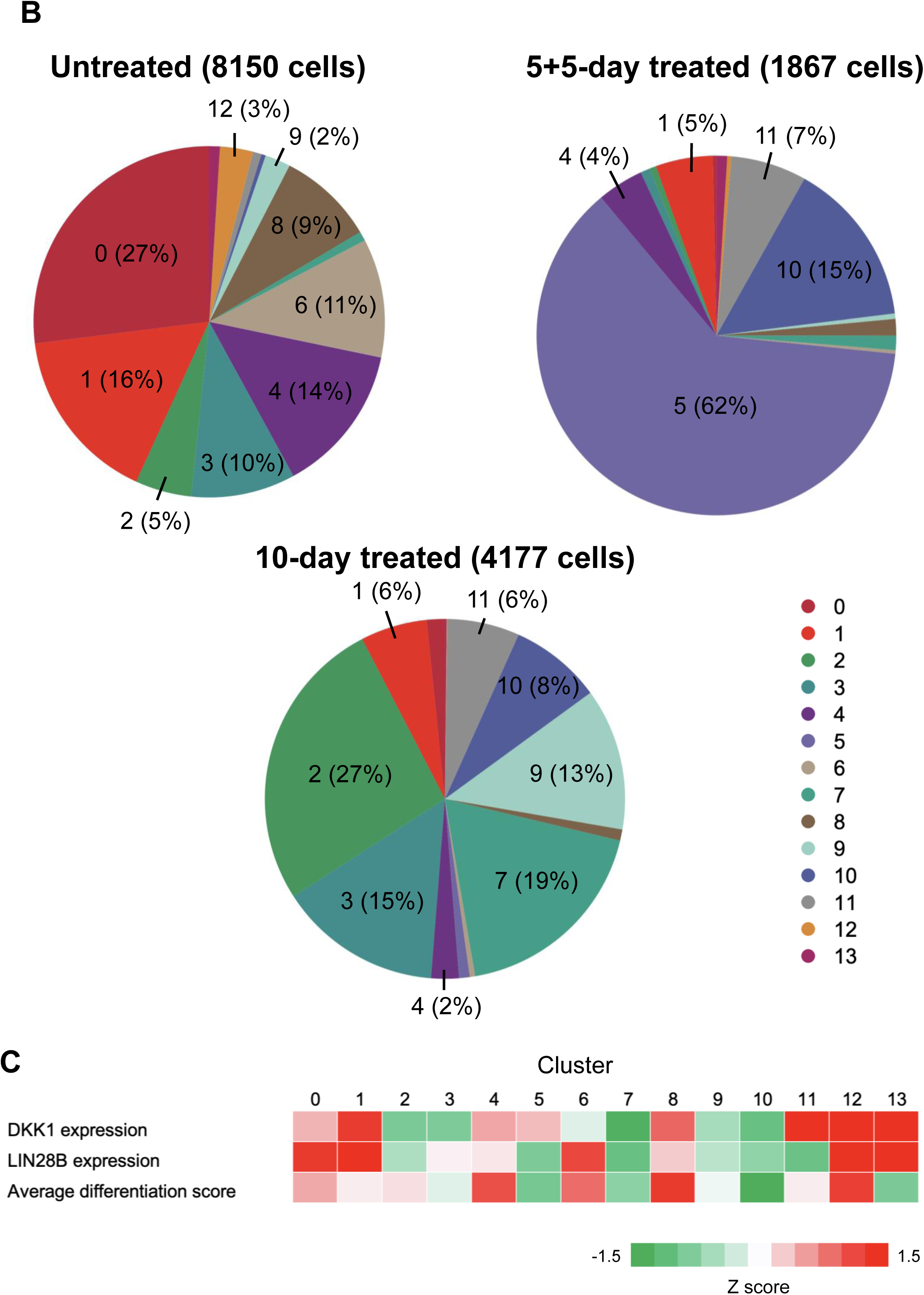
WNT-signaling inhibition promotes cancer cell differentiation in vitro. We treated the patient-derived cell line HB17 with the WNT-inhibitor ICG-001 at 5 μM for 10 days, comparing WNT-signaling activity, cell differentiation, and resistance to cisplatin, the standard-of-care chemotherapy. All cells were cultured for 10 days: either untreated, treated for 5 days with the inhibitor and the following 5 days without the inhibitor (5+5), or treated for 10 days with the inhibitor. Cells were then profiled by scRNA-seq. (**A**) UMAP of 14,194 HB17 cells grouped by 5 μM ICG-001 treatment for 5+5 or 10 days. (**B**) The composition of cell clusters (from Figure 4D) in the integrated scRNA-seq profiles identifies clusters with low and high DKK1 (WNT-signaling) and LIN28B (differentiation) expression. (**C**) Mean expression of DKK1 and LIN28B and the mean predicted differentiation score in each cell cluster after z-score transformation (mean and standard deviation based on cluster 0-11 only, due to limited cell counts in clusters 12 and 13); red and green indicate high and low values, respectively.

**Figure S6.**
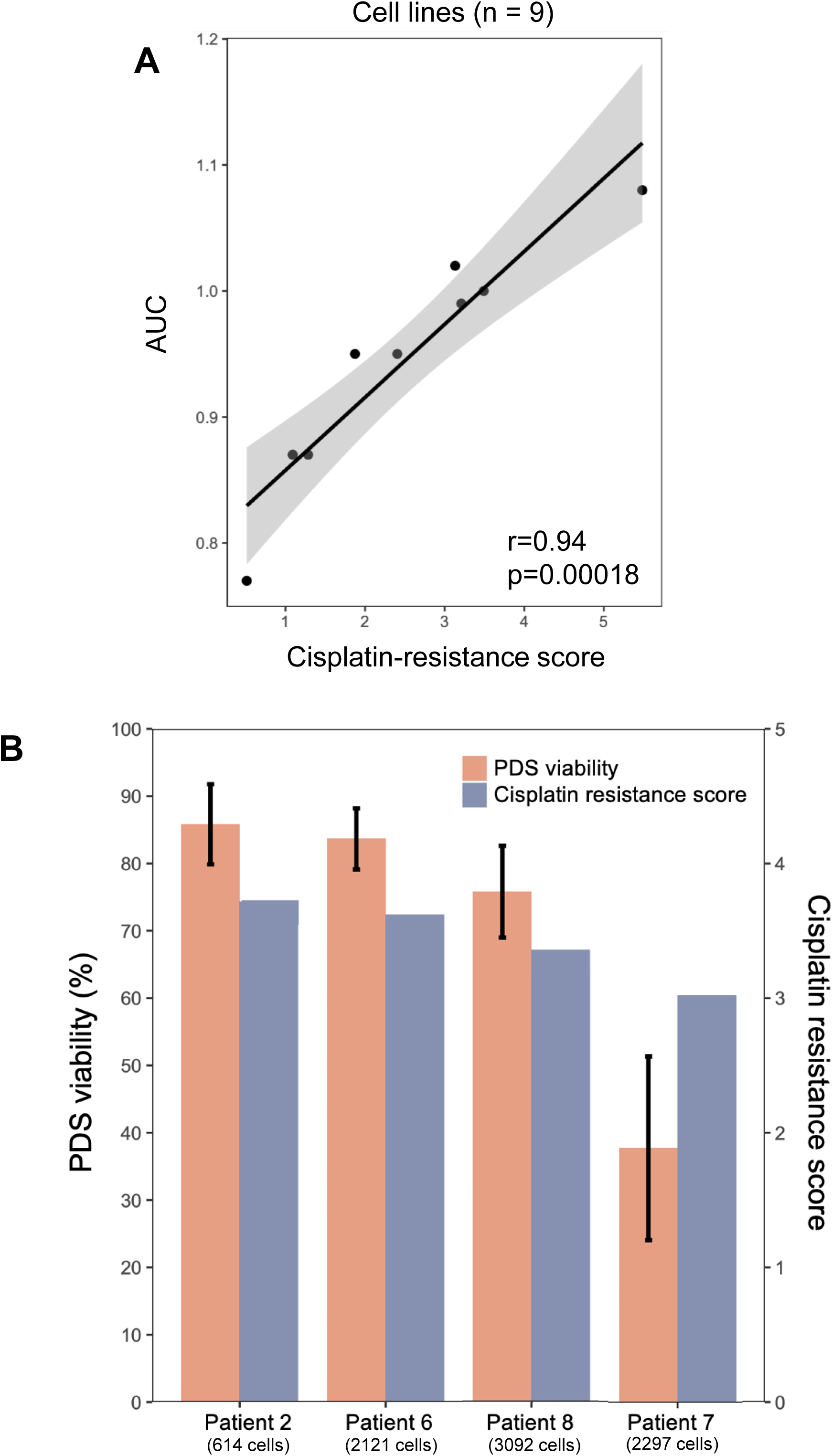
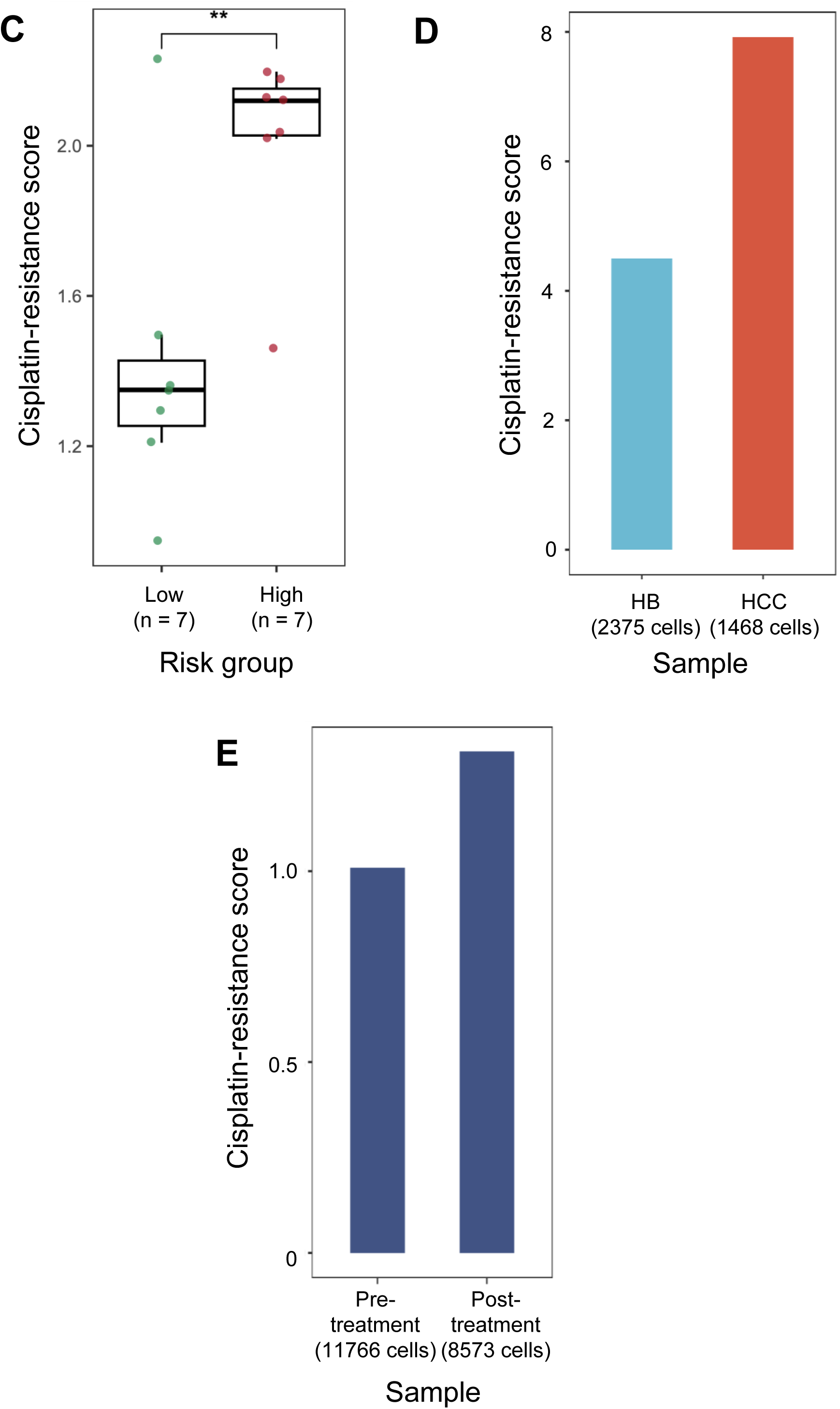
Gene expression-based scores predict resistance to cisplatin. (**A**) Correlation between predicted cisplatin-resistance scores and cisplatin drug test results, measured with the area under the curve (AUC) in nine pediatric cancer cell lines as reported by Hirsch et al (2021). Pearson correlation coefficient and associated p-value are shown. (**B**) Predicted cisplatin-resistance scores of 4 HB patient-derived organoids (PDOs) profiled and treated by Song et al. (2022), with their cell viability measurements after cisplatin treatment. Cisplatin-resistance inference was trained on bulk RNA-seq profiles and evaluated on either bulk RNA-seq or pseudobulk of scRNA-seq profiles; S.E.M. is shown. (**C**) Predicted cisplatin-resistance scores in low-risk and high-risk HB cohorts profiled by bulk RNA-seq; **: p<0.01. (**D**) Predicted cisplatin-resistance scores of our reference HB and HCC samples. (**E**) Predicted cisplatin-resistance scores of a PDX before and after cisplatin treatment profiled by scRNA-seq.

**Figure S7.**
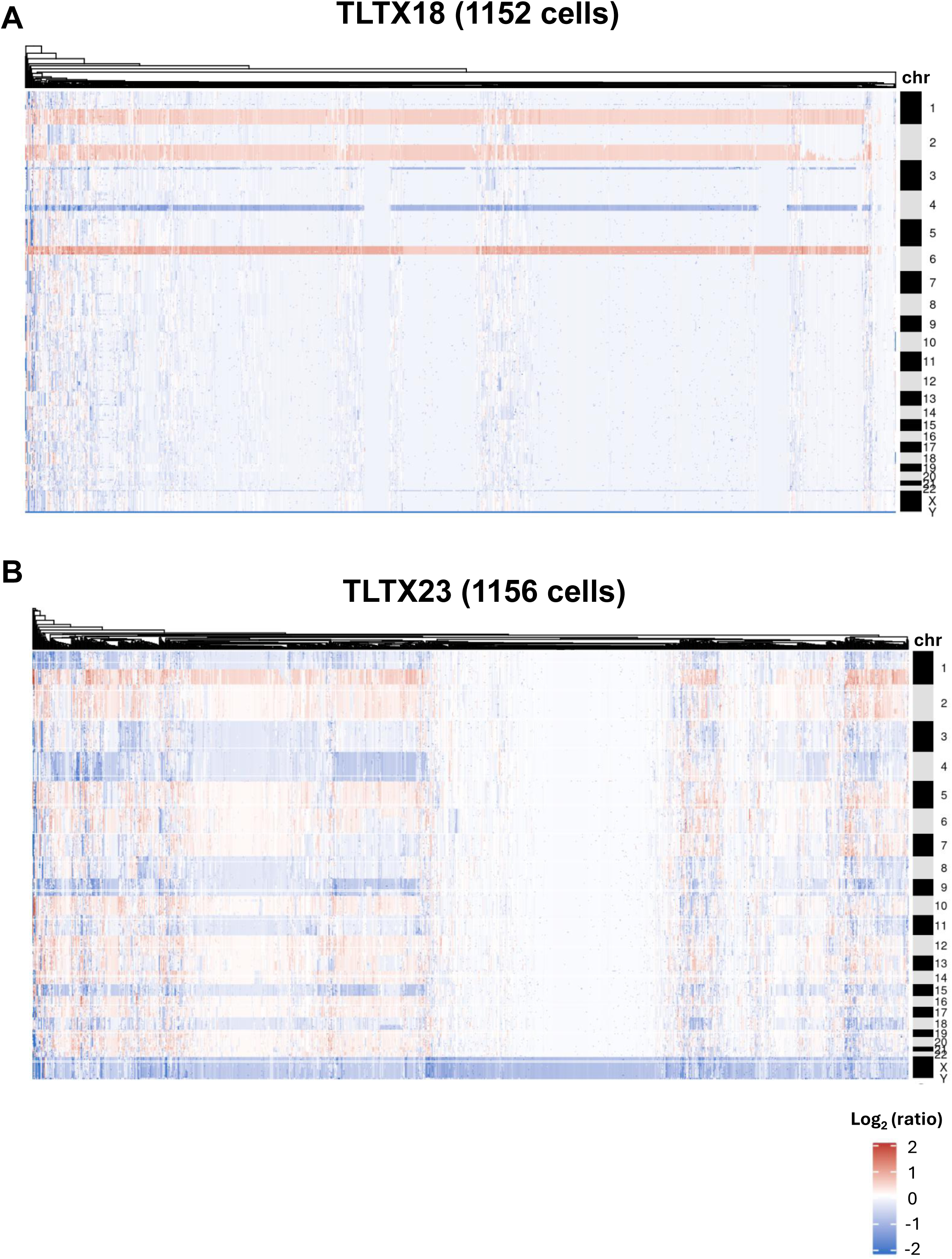
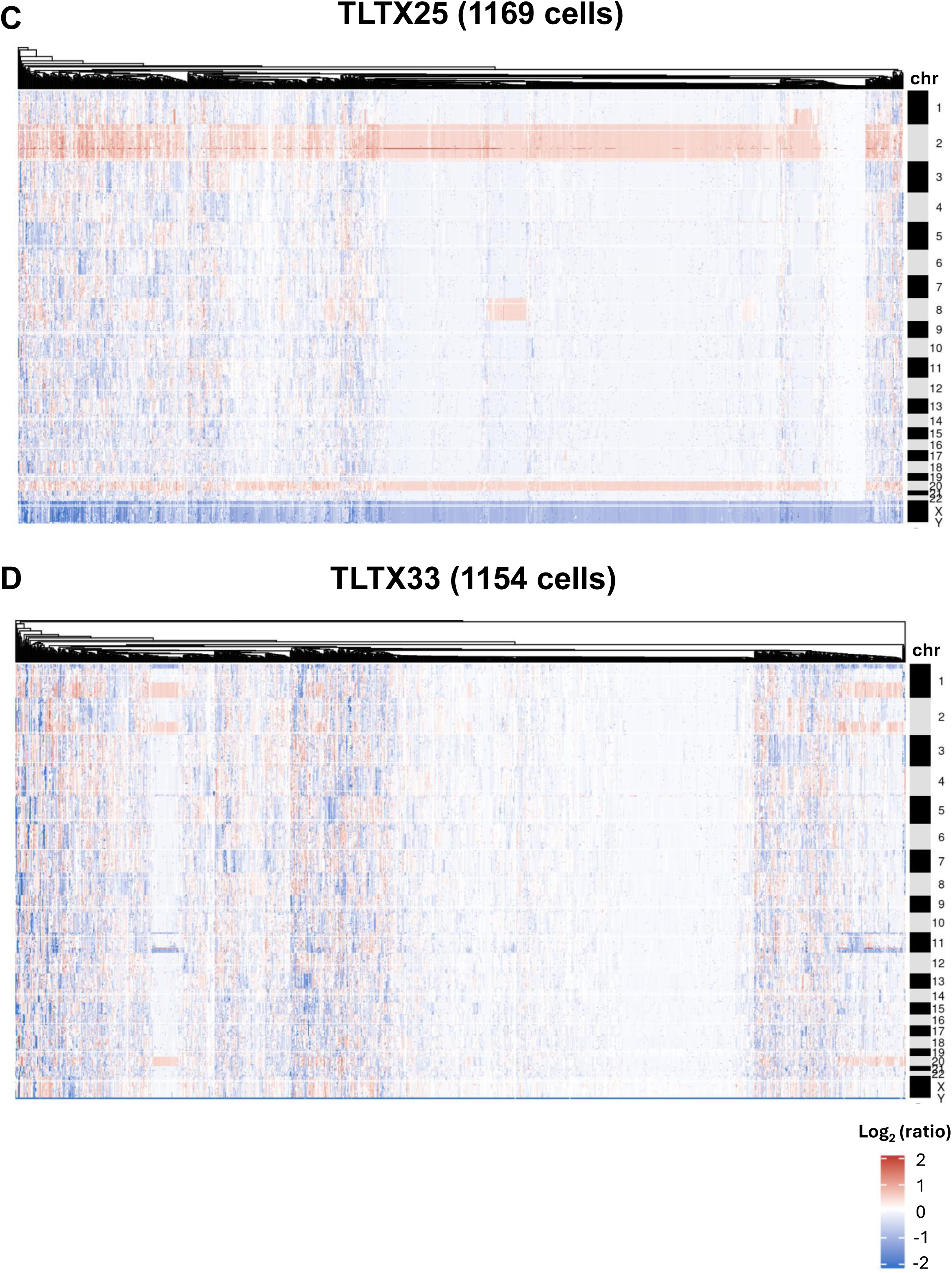
Biclustering of cells and their copy number alterations. The results of the biclusterings of the snDNA-seq profiles of (**A**) TLTX18, (**B**) TLTX23, (**C**) TLTX25, and (**D**) TLTX33.

**Figure S8.**
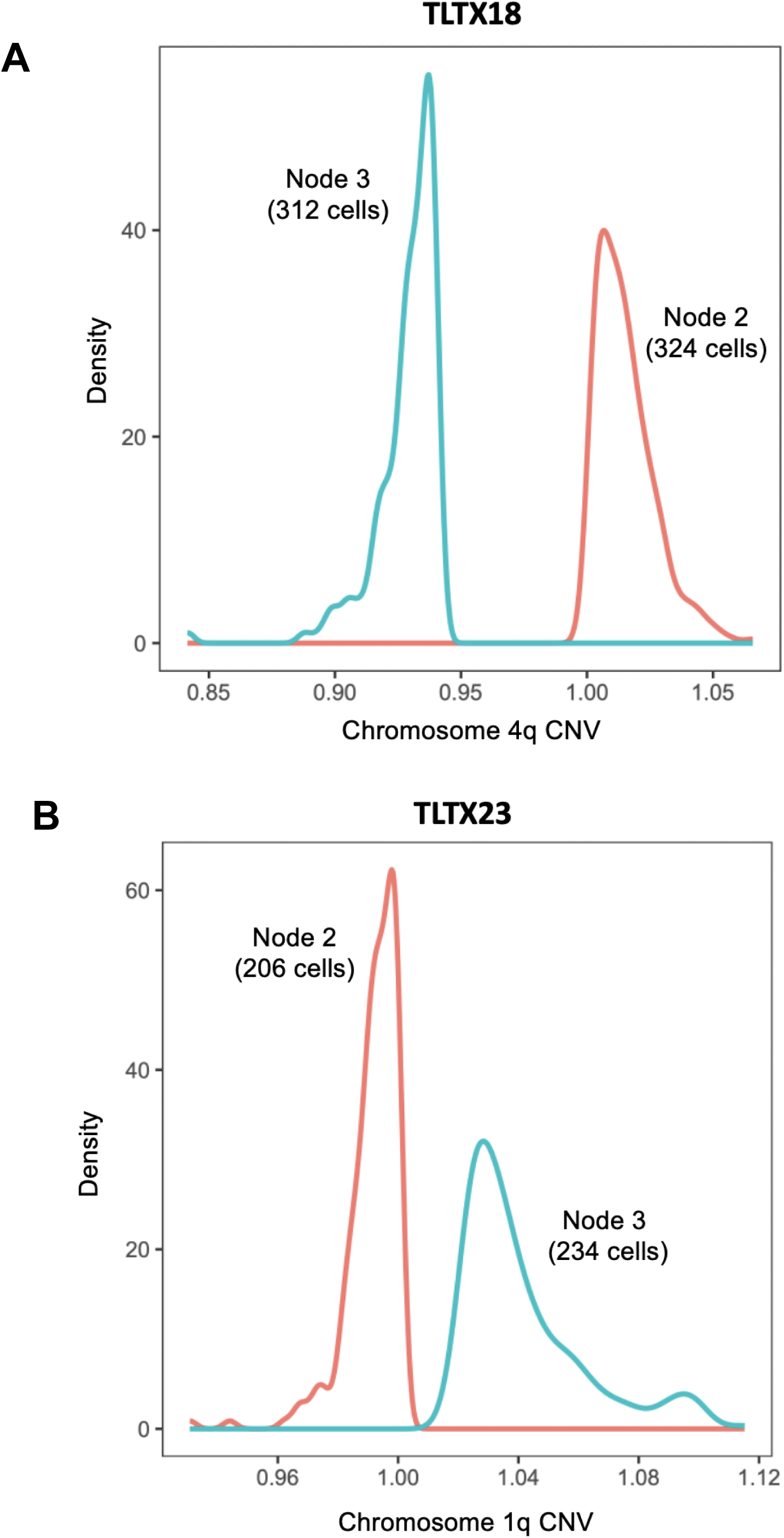
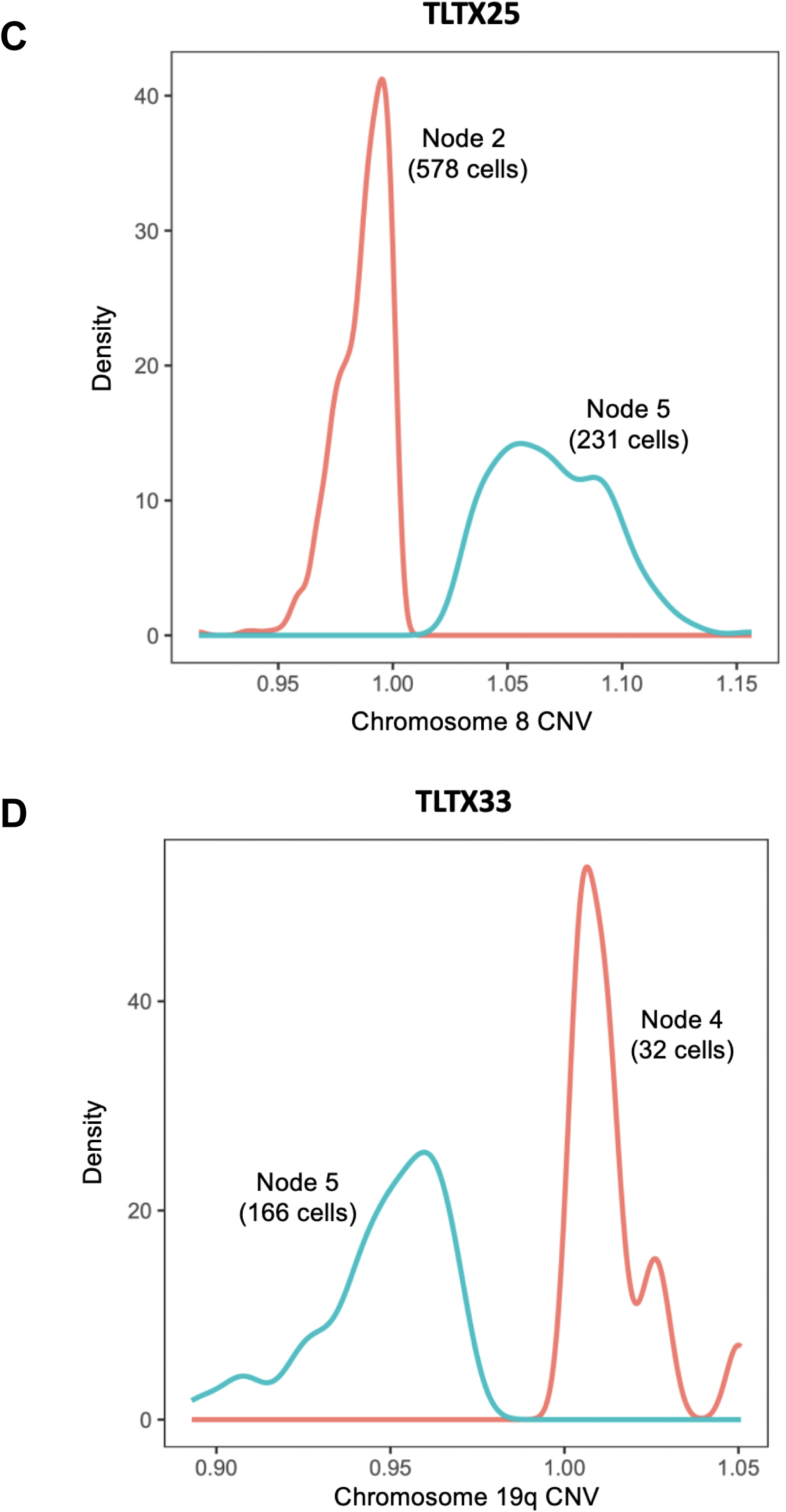
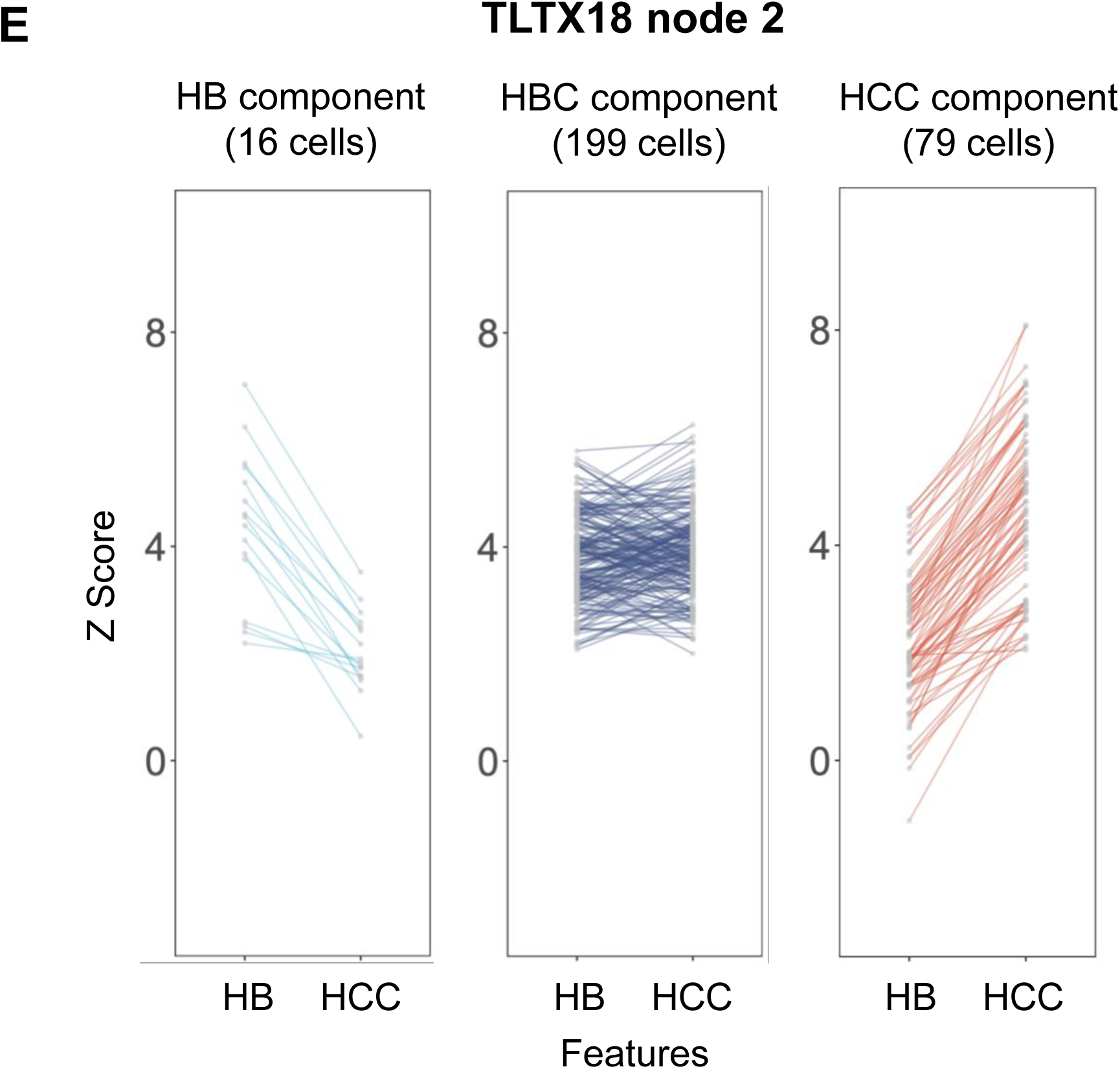
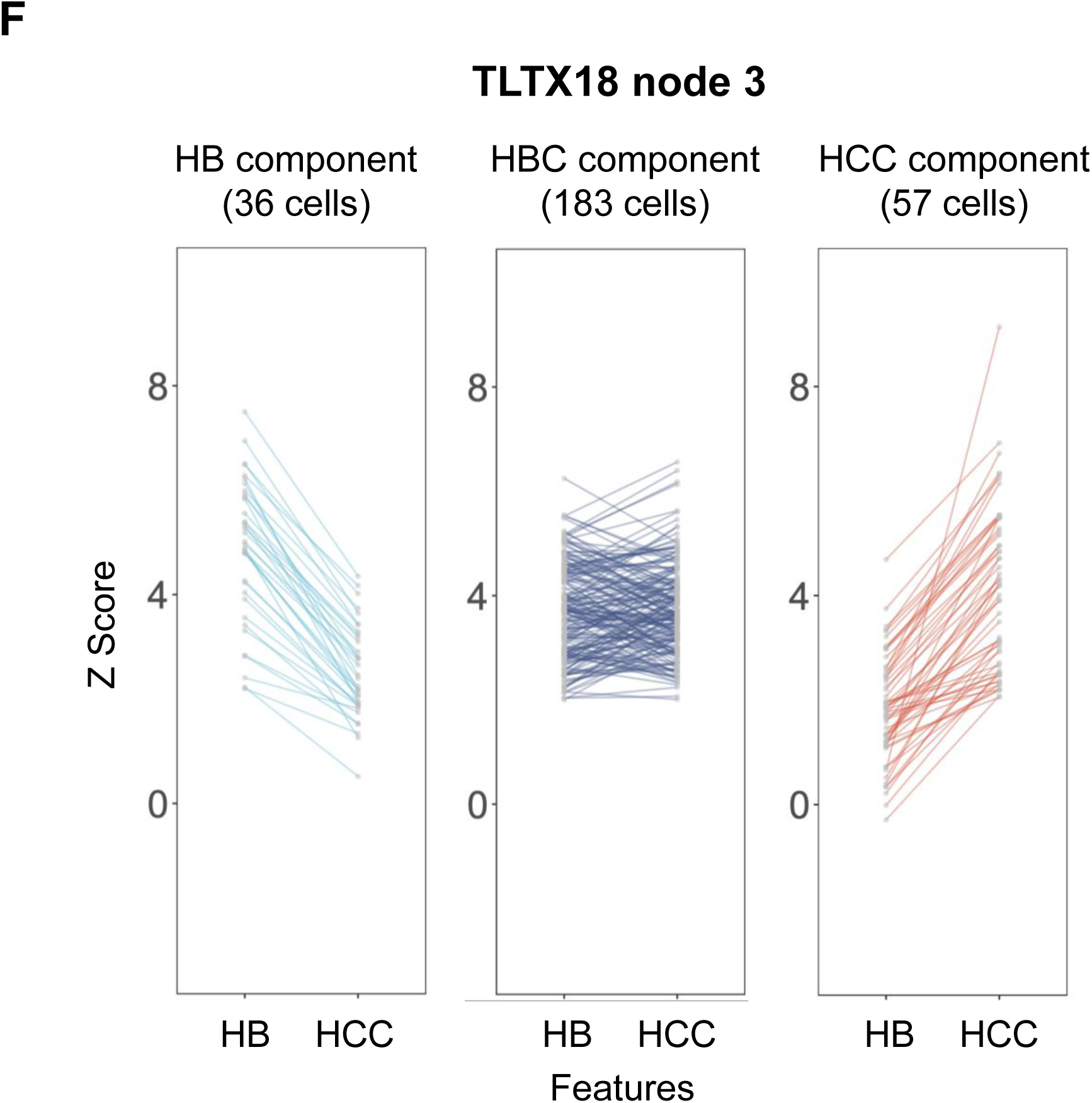
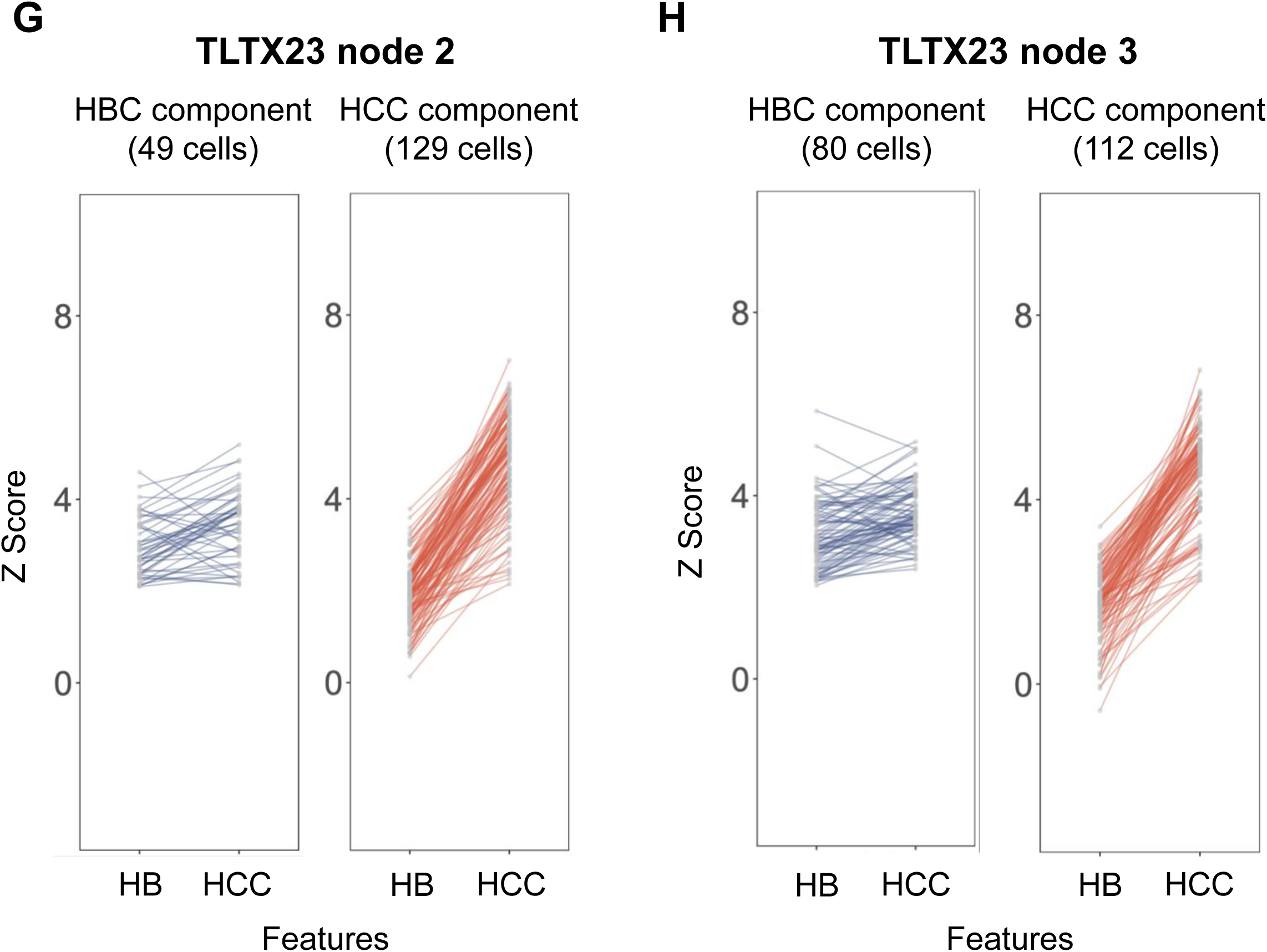
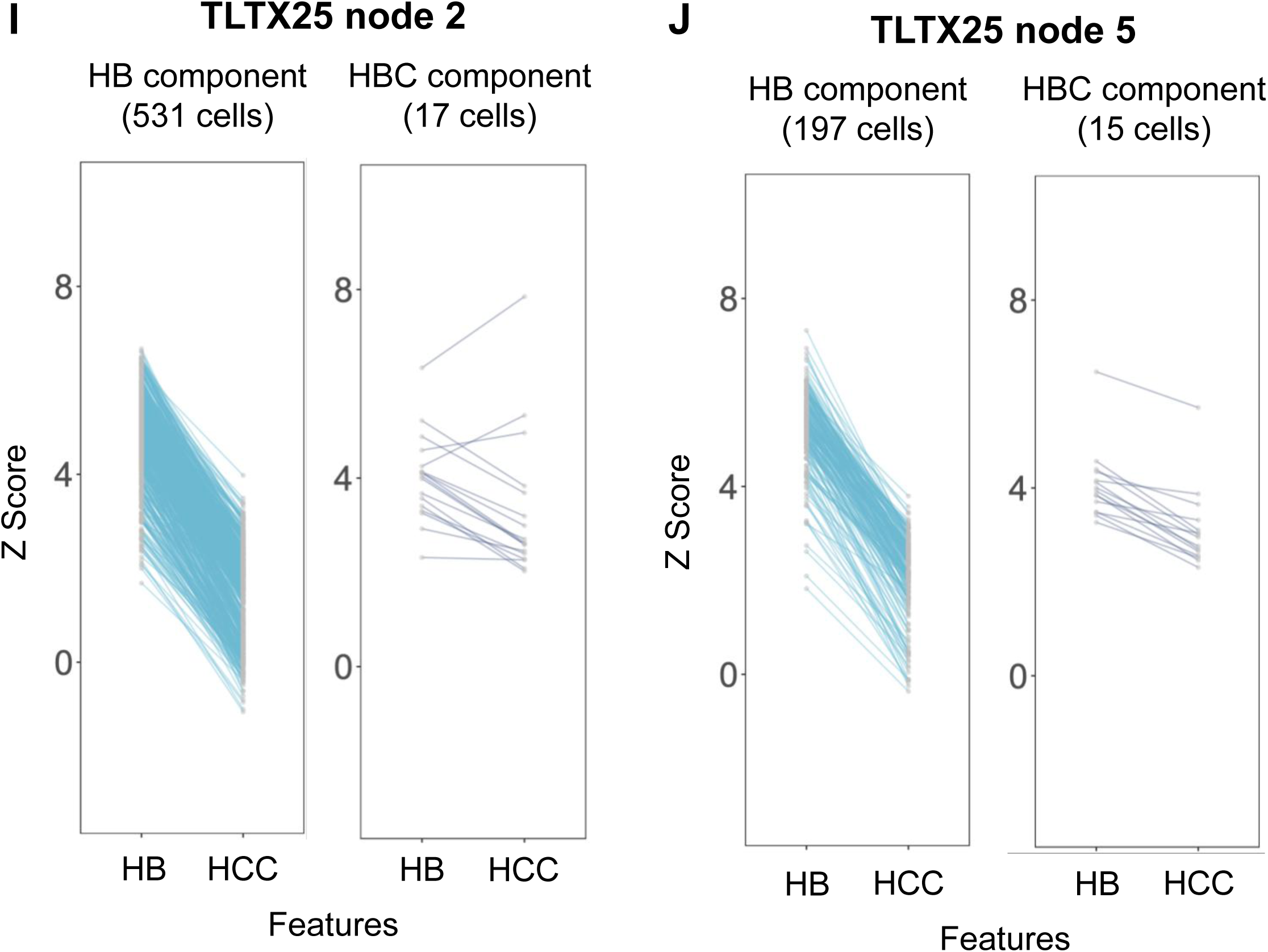
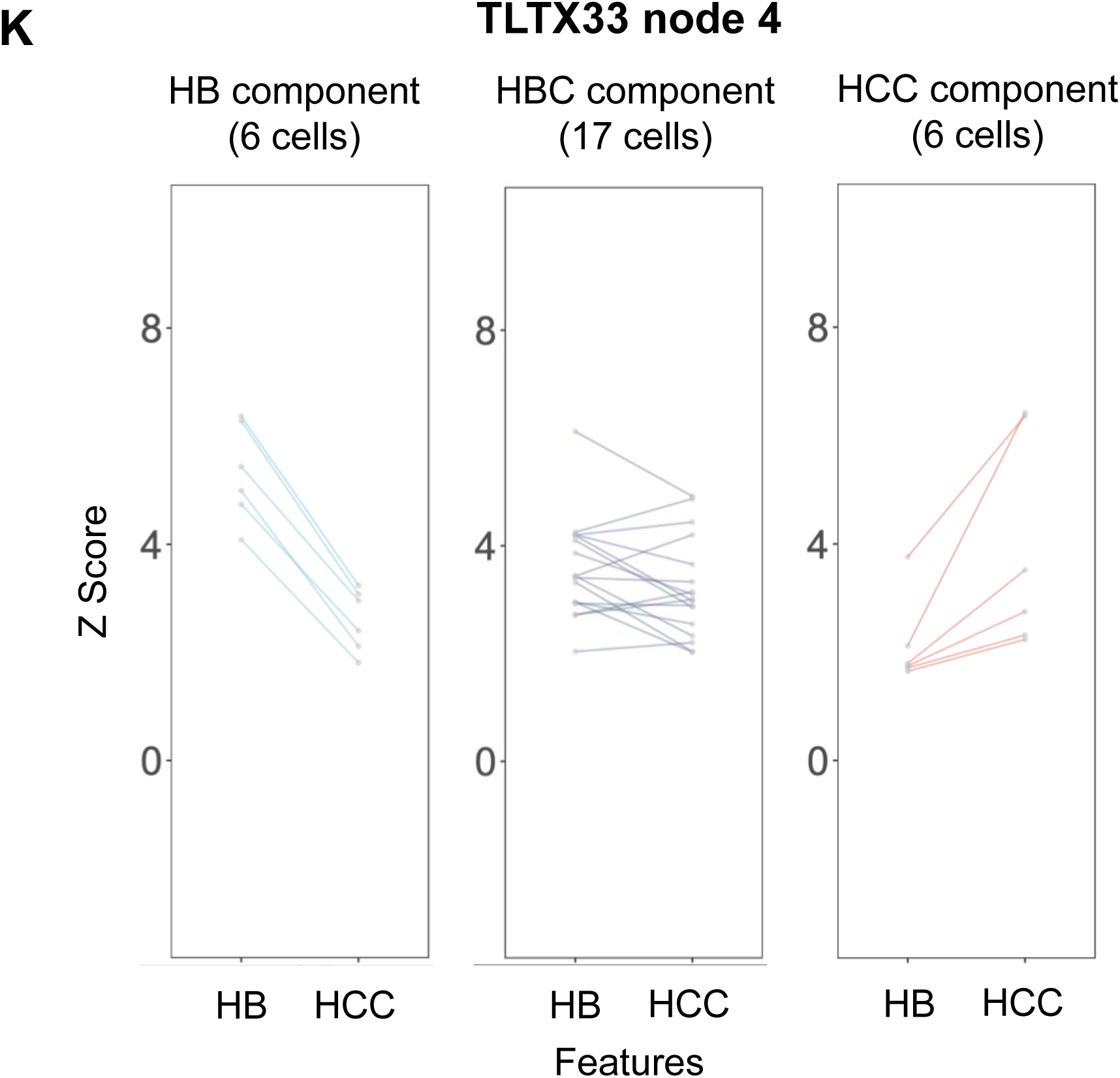
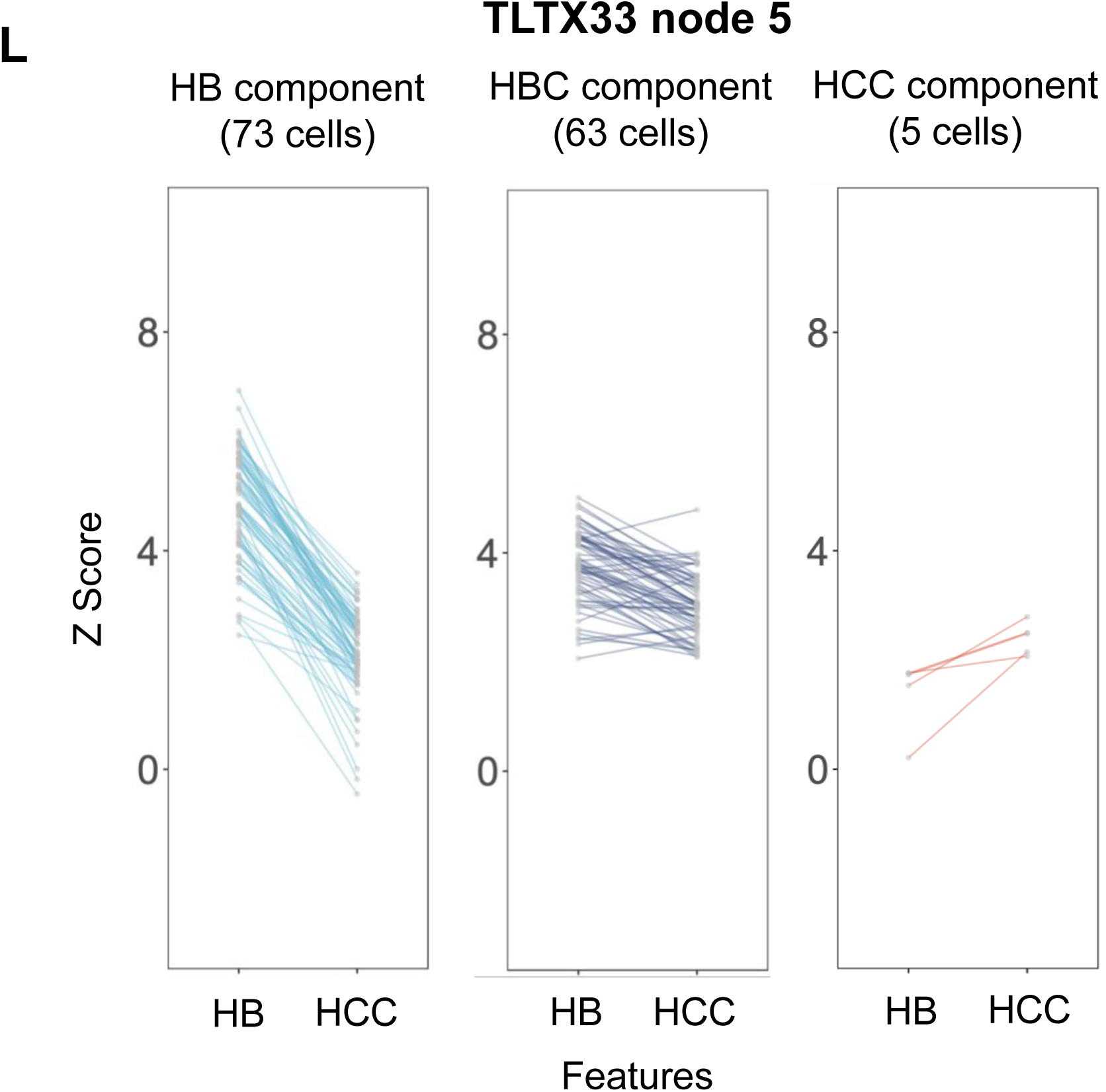
Distinct evaluations of genetic alterations across HBC phylogeny nodes. (**A-D**) Distributions of the inferCNV scores—mean normalized expression estimates across all genes in the region—in the queried regions in cells from nodes in phylogenies that are presented in Figure 5. (**A**) TLTX18 nodes 2 and 3 inferCNV scores for chromosome 4q, which was identified as lost in node 3 cells. (**B**) TLTX23 nodes 2 and 3 inferCNV scores for chromosome 1q, which was identified as gained in node 3 cells. (**C**) TLTX25 nodes 2 and 5 inferCNV scores for chromosome 8, which was identified as gained in node 5 cells. (**D**) TLTX33 nodes 4 and 5 inferCNV scores for chromosome 19q, which was identified as lost in node 5 cells. (**E-L**) Cell-type composition of genetically distinct nodes. (**E**) TLTX18 node 2 and (**F**) node 3 were both composed of HB, HBC, and HCC cells. (**G**) TLTX23 node 2 and (**H**) node 3 were both composed of HBC and HCC cells. (**I**) TLTX25 node 2 and (**J**) node 5 were both composed of HB and HBC cells. (**K**) TLTX33 node 4 and (**L**) node 5 were both composed of HB, HBC, and HCC cells.

**Figure S9.**
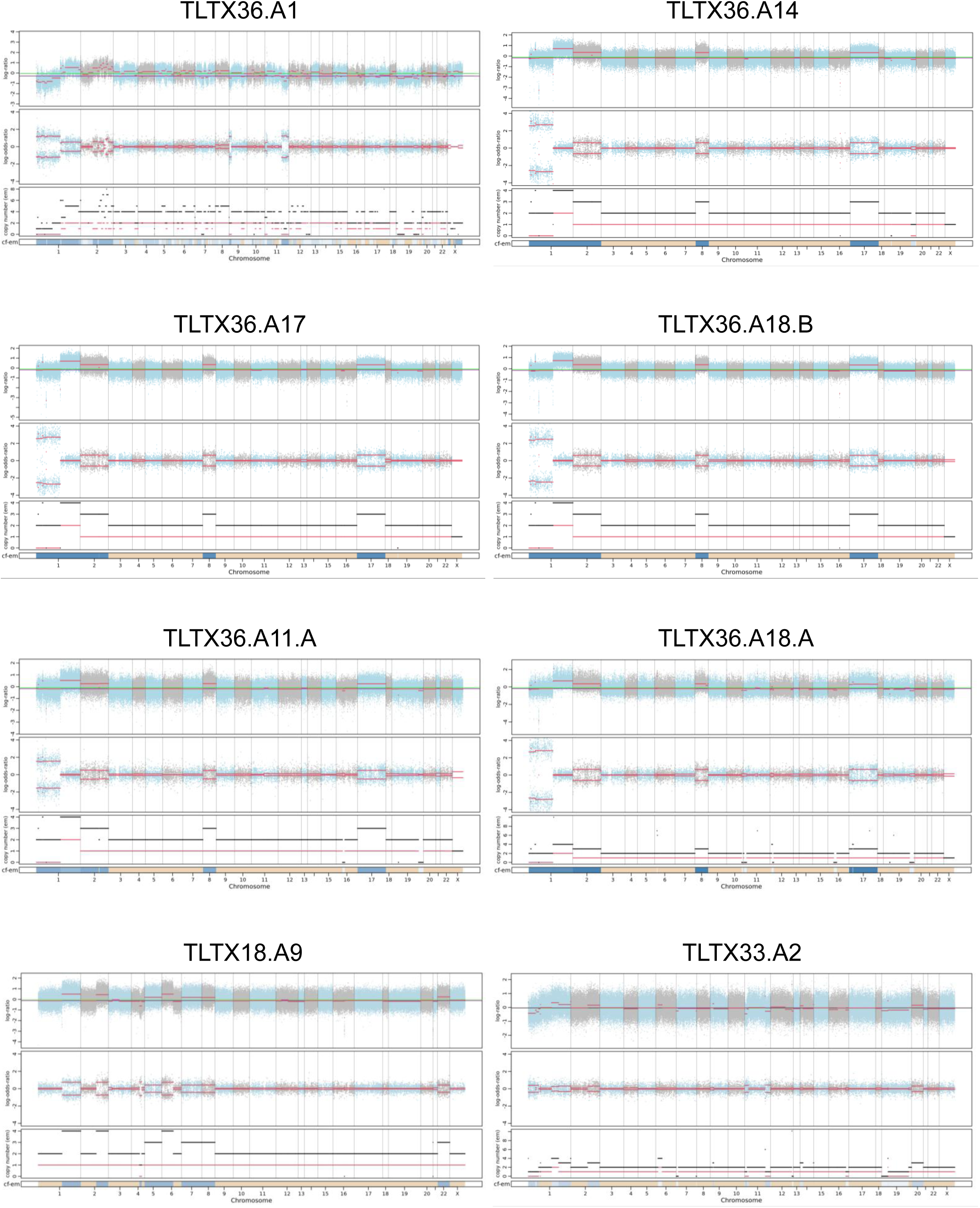
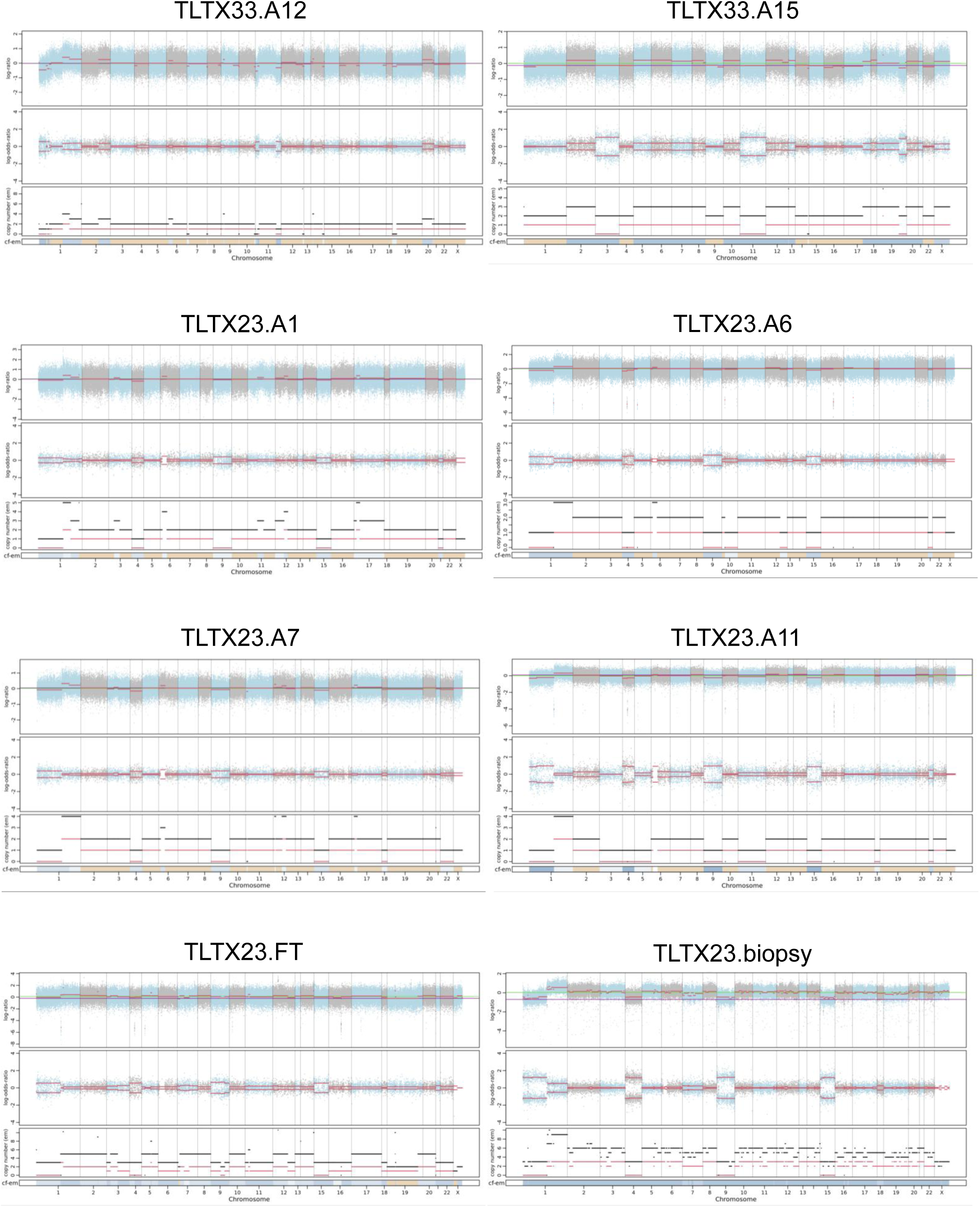
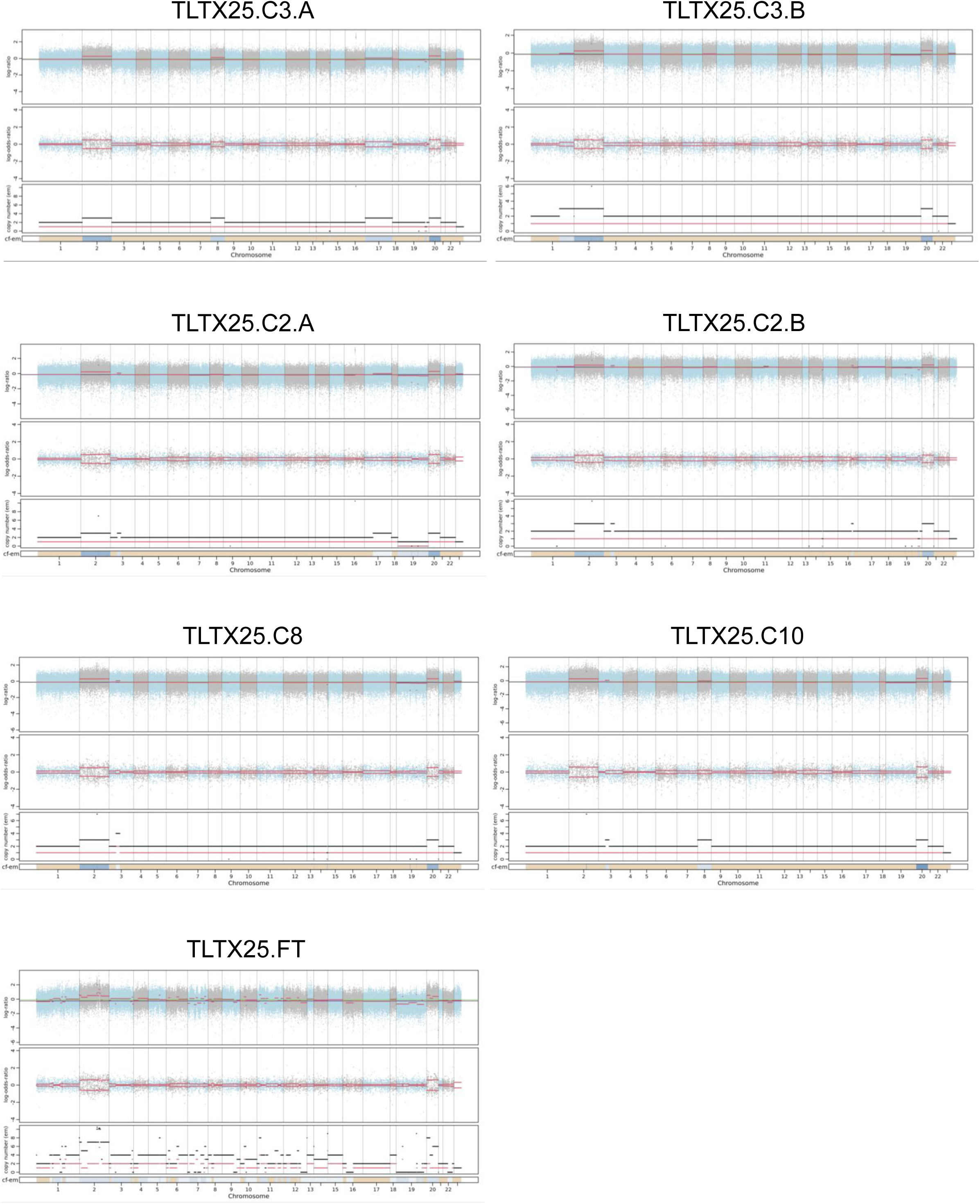
CNV calls from multi-region WES data. CNV calls from WES profiles of 6 tumor regions of TLTX36, 1 tumor region of TLTX18, 3 tumor regions of TLTX33, 6 tumor regions of TLTX23, and 7 tumor regions of TLTX25.

**Figure S10.**
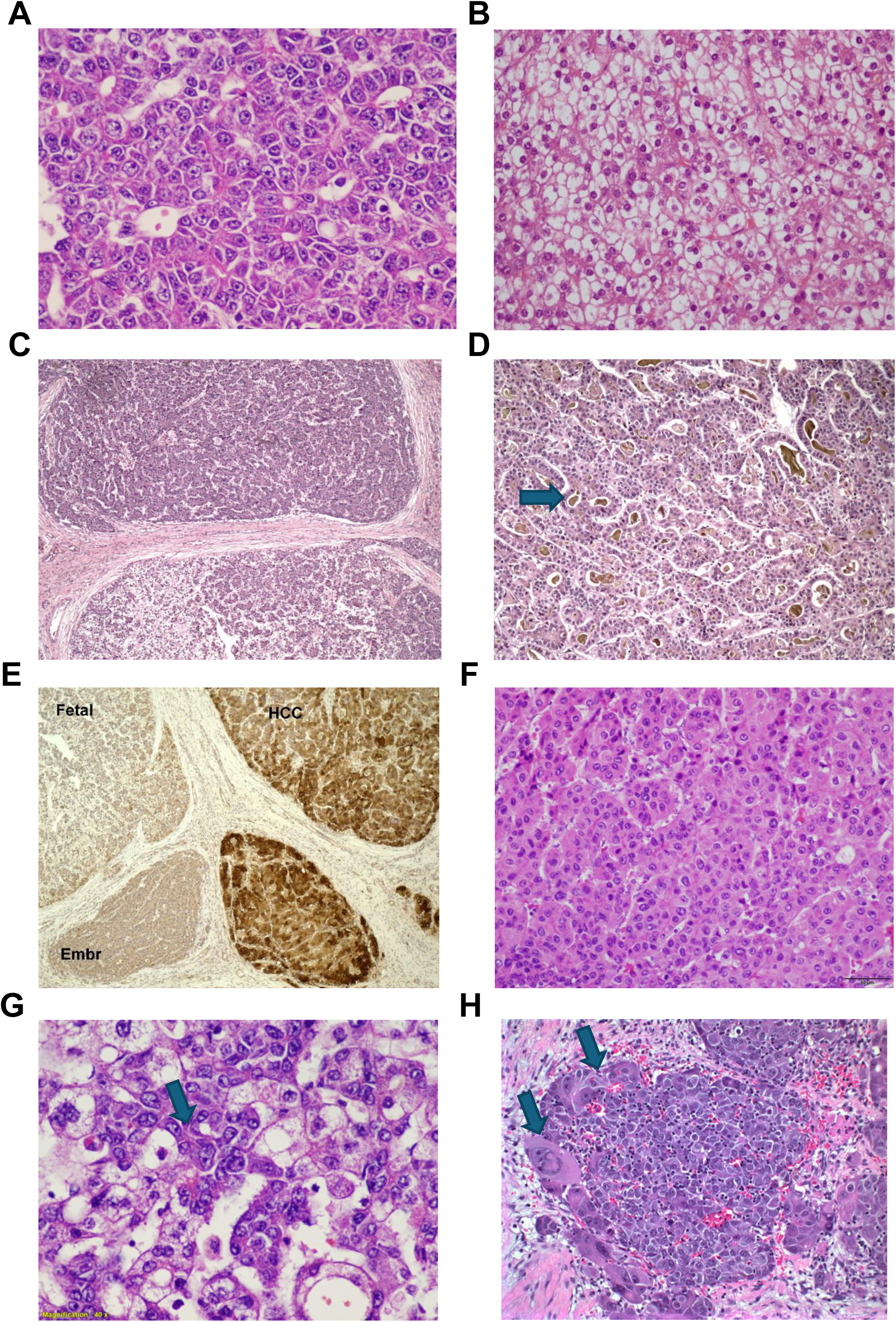
(**A**) Epithelial hepatoblastoma, embryonal pattern, focal pseudo-glandular formation. (**B**) Epithelial hepatoblastoma, fetal pattern cells with clear cytoplasm and small round nuclei. (**C**) HCN-NOS with a biphasic pattern; HB fetal and embryonal on the bottom, and HCC on top. (**D**) The HCC component of the same case (C) shows focal bile production (pointed by the arrow). (**E**) Glutamine synthetase immunohistochemistry of the same tumor, showing strong positivity in HCC areas and negative in HB fetal and embryonal areas. (**F**) Pre-treatment samples of an HCN-NOS diagnosed in a 7-year-old boy carrying a TERT promoter mutation; notice the focal macrotrabecular pattern and increased cellular pleomorphism. (**G**) Focal atypia and mild pleomorphism (arrow) in HBC post-chemotherapy. (**H**) Marked pleomorphism and frank anaplasia (arrows) in an HCN-NOS relapse.

## SUPPLEMENTARY TABLES

**Supplementary Table 1.** Clinical information of our HBC cohort and the Sumazin et al. (2017) HB cohort with published patient outcomes data.

**Supplementary Table 2.** 45 HB- and 265 HCC-specific genes were identified from our reference HB and HCC snRNA-seq sample. For each gene, the mean log2-fold changes, the proportion of HB and HCC cells expressing the gene, and the adjusted p-values are shown.

**Supplementary Table 3.** Evaluation of HB and HCC biomarker sets in reference samples—including HB, HCC, and HBC-adjacent samples—profiled at single-cell resolutions. HB and HCC biomarker z-scores of each cell are shown.

**Supplementary Table 4.** Evaluation of HB and HCC biomarker sets in bulk samples. HB and HCC biomarker z-scores of each bulk sample profiled by RNA-seq or the NanoString nCounter assay.

**Supplementary Table 5.** HB and HCC biomarker z-scores of each cell and predicted cell-type composition for 1 high-risk HB resection, 5 HBC resections, and 2 AGAR-trial HBC biopsies. HB and HCC biomarker z-scores of each cell are shown.

**Supplementary Table 6.** Hierarchical clustering of the expression values of 45 HB- and 265 HCC-specific genes in our reference HB and HCC samples and 5 HBC resections.

**Supplementary Table 7.** Selected pathways enriched in the 10 HBC tumor clusters. Adjusted p-values and normalized enrichment scores (NES) by GSEA are shown.

**Supplementary Table 8.** Evaluation of liver cell differentiation prediction in human and mouse embryonic liver cells. Predicted differentiation scores of each cell are shown.

**Supplementary Table 9.** Predicted differentiation scores of each cell for the tumor-adjacent sample, reference HB and HCC samples, AGAR-trial HBC and HCC samples, 7 HB samples by Song et al., and 5 HBC resections.

**Supplementary Table 10.** Predicted differentiation scores of each sample for pediatric and adult liver cancer and non-cancer bulk samples.

**Supplementary Table 11.** Estimated MI between differentiation scores, the expression profile of LIN28B, and the expression profiles of *let-7*miRNA targets, for each verified target. This comparison included all cells with non-zero LIN28B mRNA counts. To evaluate data processing inequality (DPI), we counted the number of verified targets where MI (differentiationScore, *let-7*target expression) was smaller than both MI (differentiationScore, LIN28B expression) and MI (LIN28B expression, *let-7*target expression). This relationship was true for 222 of the 223 *let-7* targets.

**Supplementary Table 12.** Selected pathways enriched in differentiated and undifferentiated tumor cells. Adjusted p-values and normalized enrichment scores (NES) by GSEA are shown.

**Supplementary Table 13.** Expression of DKK1 and LIN28B by quantitative PCR in wildtype untreated HB17 cells and in ICG-001-treated HB17 cells for 5 or 10 days.

**Supplementary Table 14.** Predicted differentiation scores of each cell for wildtype untreated HB17 cells and ICG-001-treated cells for 5+5 or 10 days.

**Supplementary Table 15.** The composition of cell clusters in the integrated scRNA-seq profiles of HB17 cells.

**Supplementary Table 16.** Predicted cisplatin-resistance scores of each patient in an HB cohort profiled by bulk RNA-seq.

**Supplementary Table 17.** Predicted cisplatin-resistance scores based on pseudobulk profiles of HB17 cell clusters and HBC tumor clusters.

**Supplementary Table 18.** Relative HB17 cell viability after low-dose treatment with cisplatin, ICG-001, or both in three biological replicates per condition.

**Supplementary Table 19.** Predicted differentiation scores and histological examinations of each region for TLTX36. Predicted differentiation scores and cancer cell type regarding cancer maps for TLTX18, TLTX23, TLTX25, and TLTX33 as shown in Figure 5.

**Supplementary Table 20.** Mutation calls from WES, snDNA-seq, and snRNA-seq regarding cancer maps for TLTX36, TLTX18, TLTX23, TLTX25, and TLTX33 as shown in Figure 5.

**Supplementary Table 21.** The identities of probes used in NanoString assays.

**Supplementary Table 22.** NanoString expression profiles of 15 HB, 34 HBC, 16 HCC, and 9 normal bulk samples, in addition to NanoString expression profiles of six tumor regions and one adjacent normal region from TLTX36.

## Notes

Funding declaration: This work was partially supported by the European Union Horizon 2020 award H2020-826121, which supported scientist salaries; the Cancer Prevention and Research Institute of Texas awards RP180674, RP190160, RP250580, and RP230120 funded profiling, salaries, and cancer model development; the National Institutes of Health awards S10OD032189, P30CA125123, R01CA258866, R01CA282467, R21CA286257, T32GM136554, and UH3CA271227 funded salaries, profiling, reagents, and the Single Cell Genomics Core; and the Helis Research Program by the Helis Medical Research Foundation funded salaries and cancer model development. The content is solely the responsibility of the authors and does not necessarily represent the official views of the funding agencies. This manuscript was prepared with the assistance of a science writer, Ariel M Lyons-Warren, MD, PhD.

### Competing Interest Statement

The authors have declared no competing interest.

### Summary of Updates

Updated experiments and edited the main document. Added experiments include Figures S4 and S10

## REFERENCES

[1] Hadzic N, Finegold MJ. Liver neoplasia in children. Clinics in liver disease 2011;15:443–462.

[2] López-Terrada D, Alaggio R, de Dávila MT, Czauderna P, Hiyama E, Katzenstein H, et al. Towards an international pediatric liver tumor consensus classification: proceedings of the Los Angeles COG liver tumors symposium. Modern Pathology 2014;27:472–491.

[3] Prokurat A, Kluge P, Kościesza A, Perek D, Kappeler A, Zimmermann A. Transitional liver cell tumors (TLCT) in older children and adolescents: a novel group of aggressive hepatic tumors expressing beta-catenin. Medical and pediatric oncology 2002;39:510–518.

[4] Eichenmuller M, Trippel F, Kreuder M, Beck A, Schwarzmayr T, Haberle B, et al. The genomic landscape of hepatoblastoma and their progenies with HCC-like features. J Hepatol 2014;61:1312–1320.

[5] Sumazin P, Peters TL, Sarabia SF, Kim HR, Urbicain M, Hollingsworth EF, et al. Hepatoblastomas with carcinoma features represent a biological spectrum of aggressive neoplasms in children and young adults. Journal of Hepatology 2022;77:1026–1037.

[6] Shrivastava N, O’Neill AF. Update in Solid Tumors of Childhood. Update in Pediatrics: Springer; 2024. p. 629–662.

[7] Czauderna P, Lopez-Terrada D, Hiyama E, Häberle B, Malogolowkin MH, Meyers RL. Hepatoblastoma state of the art: pathology, genetics, risk stratification, and chemotherapy. Current opinion in pediatrics 2014;26:19–28.

[8] Lau CS, Mahendraraj K, Chamberlain RS. Hepatocellular Carcinoma in the Pediatric Population: A Population Based Clinical Outcomes Study Involving 257 Patients from the Surveillance, Epidemiology, and End Result (SEER) Database (1973-2011). HPB Surg 2015;2015:670728.

[9] Espinoza AF, Patel R, Patel KR, Badachhap AA, Whitlock R, Srivastava RK, et al. A Novel Treatment Strategy Utilizing Panobinostat for High-Risk and Treatment-Refractory Hepatoblastoma. Journal of Hepatology 2024;80:610–621.

[10] Czauderna P, Haeberle B, Hiyama E, Rangaswami A, Krailo M, Maibach R, et al. The Children’s Hepatic tumors International Collaboration (CHIC): Novel global rare tumor database yields new prognostic factors in hepatoblastoma and becomes a research model. European journal of cancer 2016;52:92–101.

[11] Cairo S, Armengol C, Maibach R, Häberle B, Becker K, Carrillo-Reixach J, et al. A combined clinical and biological risk classification improves prediction of outcome in hepatoblastoma patients. European Journal of Cancer 2020;141:30–39.

[12] Cairo S, Armengol C, De Reyniès A, Wei Y, Thomas E, Renard CA, et al. Hepatic stem-like phenotype and interplay of Wnt/beta-catenin and Myc signaling in aggressive childhood liver cancer. Cancer Cell 2008;14:471–484.

[13] Sumazin P, Chen Y, Treviño LR, Sarabia SF, Hampton OA, Patel K, et al. Genomic analysis of hepatoblastoma identifies distinct molecular and prognostic subgroups. Hepatology 2017;65:104–121.

[14] Nagae G, Yamamoto S, Fujita M, Fujita T, Nonaka A, Umeda T, et al. Genetic and epigenetic basis of hepatoblastoma diversity. Nature Communications 2021;12:5423.

[15] Hirsch TZ, Pilet J, Morcrette G, Roehrig A, Monteiro BJE, Molina L, et al. Integrated Genomic Analysis Identifies Driver Genes and Cisplatin-Resistant Progenitor Phenotype in Pediatric Liver Cancer. Cancer discovery 2021;11:2524–2543.

[16] Bondoc A, Glaser K, Jin K, Lake C, Cairo S, Geller J, et al. Identification of distinct tumor cell populations and key genetic mechanisms through single cell sequencing in hepatoblastoma. Commun Biol 2021;4:1049.

[17] Steffin D, Ghatwai N, Montalbano A, Rathi P, Courtney AN, Arnett AB, et al. Interleukin-15-armoured GPC3 CAR T cells for patients with solid cancers. Nature 2025;637:940–946.

[18] Boster JM, Superina R, Mazariegos GV, Tiao GM, Roach JP, Lovell MA, et al. Predictors of survival following liver transplantation for pediatric hepatoblastoma and hepatocellular carcinoma: Experience from the Society of Pediatric Liver Transplantation (SPLIT). Am J Transplant 2022;22:1396–1408.

[19] Wang X, Yang L, Wang Y-C, Xu Z-R, Feng Y, Zhang J, et al. Comparative analysis of cell lineage differentiation during hepatogenesis in humans and mice at the single-cell transcriptome level. Cell Research 2020;30:1109–1126.

[20] Song H, Bucher S, Rosenberg K, Tsui M, Burhan D, Hoffman D, et al. Single-cell analysis of hepatoblastoma identifies tumor signatures that predict chemotherapy susceptibility using patient-specific tumor spheroids. Nature communications 2022;13:4878.

[21] TCGA. Comprehensive and Integrative Genomic Characterization of Hepatocellular Carcinoma. Cell 2017;169:1327–1341.e1323.

[22] Talukdar S, Bhoopathi P, Emdad L, Das S, Sarkar D, Fisher PB. Dormancy and cancer stem cells: An enigma for cancer therapeutic targeting. Advances in cancer research 2019;141:43–84.

[23] Nguyen LH, Robinton DA, Seligson MT, Wu L, Li L, Rakheja D, et al. Lin28b is sufficient to drive liver cancer and necessary for its maintenance in murine models. Cancer Cell 2014;26:248–261.

[24] Piskounova E, Polytarchou C, Thornton JE, LaPierre RJ, Pothoulakis C, Hagan JP, et al. Lin28A and Lin28B inhibit let-7 microRNA biogenesis by distinct mechanisms. Cell 2011;147:1066–1079.

[25] Ustianenko D, Chiu H-S, Treiber T, Weyn-Vanhentenryck SM, Treiber N, Meister G, et al. LIN28 selectively modulates a subclass of let-7 microRNAs. Molecular cell 2018;71:271–283. e275.

[26] Viswanathan SR, Daley GQ, Gregory RI. Selective blockade of microRNA processing by Lin28. Science 2008;320:97–100.

[27] Takashima Y, Terada M, Udono M, Miura S, Yamamoto J, Suzuki A. Suppression of lethal-7b and miR-125a/b Maturation by Lin28b Enables Maintenance of Stem Cell Properties in Hepatoblasts. Hepatology 2016;64:245–260.

[28] Armengol C, Cairo S, Fabre M, Buendia M. Wnt signaling and hepatocarcinogenesis: the hepatoblastoma model. The international journal of biochemistry & cell biology 2011;43:265–270.

[29] Nelson WJ, Nusse R. Convergence of Wnt, beta-catenin, and cadherin pathways. Science 2004;303:1483–1487.

[30] Thompson MD, Monga SP. WNT/beta-catenin signaling in liver health and disease. Hepatology 2007;45:1298–1305.

[31] Emami KH, Nguyen C, Ma H, Kim DH, Jeong KW, Eguchi M, et al. A small molecule inhibitor of beta-catenin/CREB-binding protein transcription [corrected]. Proc Natl Acad Sci U S A 2004;101:12682–12687.

[32] Saeki I, Ida K, Kurihara S, Watanabe K, Mori M, Hishiki T, et al. Successful treatment of young childhood standard-risk hepatoblastoma with cisplatin monotherapy using a central review system. Pediatric blood & cancer 2024;71:e31255.

[33] Meyers RL, Maibach R, Hiyama E, Häberle B, Krailo M, Rangaswami A, et al. Risk-stratified staging in paediatric hepatoblastoma: a unified analysis from the Children’s Hepatic tumors International Collaboration. The Lancet Oncology 2017;18:122–131.

[34] Manica M, Kim HR, Mathis R, Chouvarine P, Rutishauser D, De Vargas Roditi L, et al. Inferring clonal composition from multiple tumor biopsies. NPJ systems biology and applications 2020;6:27.

[35] Navin N, Kendall J, Troge J, Andrews P, Rodgers L, McIndoo J, et al. Tumour evolution inferred by single-cell sequencing. Nature 2011;472:90–94.

[36] Wang Y, Waters J, Leung ML, Unruh A, Roh W, Shi X, et al. Clonal evolution in breast cancer revealed by single nucleus genome sequencing. Nature 2014;512:155–160.

[37] Parsons DW, Janeway KA, Patton DR, Winter CL, Coffey B, Williams PM, et al. Actionable Tumor Alterations and Treatment Protocol Enrollment of Pediatric and Young Adult Patients With Refractory Cancers in the National Cancer Institute-Children’s Oncology Group Pediatric MATCH Trial. Journal of clinical oncology: official journal of the American Society of Clinical Oncology 2022;40:2224–2234.

[38] David MP, Venkatramani R, Lopez-Terrada DH, Roy A, Patil N, Fisher KE. Multimodal molecular analysis of an atypical small cell carcinoma of the ovary, hypercalcemic type. Cold Spring Harb Mol Case Stud 2018;4.

[39] Tirosh I, Izar B, Prakadan SM, Wadsworth MH, 2nd, Treacy D, Trombetta JJ, et al. Dissecting the multicellular ecosystem of metastatic melanoma by single-cell RNA-seq. Science 2016;352:189–196.

[40] Bakker B, Taudt A, Belderbos ME, Porubsky D, Spierings DC, de Jong TV, et al. Single-cell sequencing reveals karyotype heterogeneity in murine and human malignancies. Genome Biol 2016;17:115.

[41] Scollon S, Eldomery MK, Reuther J, Lin FY, Potter SL, Desrosiers L, et al. Clinical and molecular features of pediatric cancer patients with Lynch syndrome. Pediatric blood & cancer 2022;69:e29859.

[42] Woodfield SE, Shi Y, Patel RH, Chen Z, Shah AP, Srivastava RK, et al. MDM4 inhibition: a novel therapeutic strategy to reactivate p53 in hepatoblastoma. Scientific reports 2021;11:2967.

[43] Bissig-Choisat B, Kettlun-Leyton C, Legras XD, Zorman B, Barzi M, Chen LL, et al. Novel patient-derived xenograft and cell line models for therapeutic testing of pediatric liver cancer. J Hepatol 2016;65:325–333.

[44] Whitfield ML, Sherlock G, Saldanha AJ, Murray JI, Ball CA, Alexander KE, et al. Identification of genes periodically expressed in the human cell cycle and their expression in tumors. Mol Biol Cell 2002;13:1977–2000.

## REFERENCES

1 Tirosh, I. et al. Dissecting the multicellular ecosystem of metastatic melanoma by single-cell RNA-seq. Science 352, 189–196, doi:10.1126/science.aad0501 (2016).

2 Minussi, D. C. et al. Breast tumours maintain a reservoir of subclonal diversity during expansion. Nature 592, 302–308, doi:10.1038/s41586-021-03357-x (2021).

3 Hao, Y. et al. Integrated analysis of multimodal single-cell data. Cell 184, 3573–3587.e3529, doi:10.1016/j.cell.2021.04.048 (2021).

4 Stuart, T. et al. Comprehensive Integration of Single-Cell Data. Cell 177, 1888–1902.e1821, doi:10.1016/j.cell.2019.05.031 (2019).

5 Cairo, S. et al. Hepatic stem-like phenotype and interplay of Wnt/beta-catenin and Myc signaling in aggressive childhood liver cancer. Cancer Cell 14, 471–484, doi:10.1016/j.ccr.2008.11.002 (2008).

6 Sumazin, P. et al. Genomic analysis of hepatoblastoma identifies distinct molecular and prognostic subgroups. Hepatology 65, 104–121, doi:10.1002/hep.28888 (2017).

7 Aran, D. et al. Reference-based analysis of lung single-cell sequencing reveals a transitional profibrotic macrophage. Nat Immunol 20, 163–172, doi:10.1038/s41590-018-0276-y (2019).

8 Conway, T. et al. Xenome--a tool for classifying reads from xenograft samples. Bioinformatics 28, i172–178, doi:10.1093/bioinformatics/bts236 (2012).

9 Bakker, B. et al. Single-cell sequencing reveals karyotype heterogeneity in murine and human malignancies. Genome Biol 17, 115, doi:10.1186/s13059-016-0971-7 (2016).

10 Wingett, S. W. & Andrews, S. FastQ Screen: A tool for multi-genome mapping and quality control. F1000Res 7, 1338, doi:10.12688/f1000research.15931.2 (2018).

11 Bolger, A. M., Lohse, M. & Usadel, B. Trimmomatic: a flexible trimmer for Illumina sequence data. Bioinformatics 30, 2114–2120, doi:10.1093/bioinformatics/btu170 (2014).

12 Scollon, S. et al. Clinical and molecular features of pediatric cancer patients with Lynch syndrome. Pediatric blood & cancer 69, e29859, doi:10.1002/pbc.29859 (2022).

13 McKenna, A. et al. The Genome Analysis Toolkit: a MapReduce framework for analyzing next-generation DNA sequencing data. Genome Res 20, 1297–1303, doi:10.1101/gr.107524.110 (2010).

14 Grossman, R. L. et al. Toward a Shared Vision for Cancer Genomic Data. N Engl J Med 375, 1109–1112, doi:10.1056/NEJMp1607591 (2016).

15 Li, H. Toward better understanding of artifacts in variant calling from high-coverage samples. Bioinformatics 30, 2843–2851, doi:10.1093/bioinformatics/btu356 (2014).

16 Danecek, P. et al. Twelve years of SAMtools and BCFtools. Gigascience 10, doi:10.1093/gigascience/giab008 (2021).

17 Cibulskis, K. et al. Sensitive detection of somatic point mutations in impure and heterogeneous cancer samples. Nat Biotechnol 31, 213–219, doi:10.1038/nbt.2514 (2013).

18 Wang, K., Li, M. & Hakonarson, H. ANNOVAR: functional annotation of genetic variants from high-throughput sequencing data. Nucleic Acids Res 38, e164, doi:10.1093/nar/gkq603 (2010).

19 Shen, R. & Seshan, V. E. FACETS: allele-specific copy number and clonal heterogeneity analysis tool for high-throughput DNA sequencing. Nucleic Acids Res 44, e131, doi:10.1093/nar/gkw520 (2016).

20 Dentro, S. C., Wedge, D. C. & Van Loo, P. Principles of Reconstructing the Subclonal Architecture of Cancers. Cold Spring Harb Perspect Med 7, doi:10.1101/cshperspect.a026625 (2017).

21 Sumazin, P. et al. Hepatoblastomas with carcinoma features represent a biological spectrum of aggressive neoplasms in children and young adults. Journal of Hepatology 77, 1026–1037 (2022).

22 Korinek, V. et al. Constitutive transcriptional activation by a beta-catenin-Tcf complex in APC-/-colon carcinoma. Science 275, 1784–1787, doi:10.1126/science.275.5307.1784 (1997).

23 Gillis, S. & Roth, A. PyClone-VI: scalable inference of clonal population structures using whole genome data. BMC Bioinformatics 21, 571, doi:10.1186/s12859-020-03919-2 (2020).

24 Manica, M. et al. Inferring clonal composition from multiple tumor biopsies. NPJ systems biology and applications 6, 27 (2020).

25 Song, H. et al. Single-cell analysis of hepatoblastoma identifies tumor signatures that predict chemotherapy susceptibility using patient-specific tumor spheroids. Nature communications 13, 4878 (2022).

26 Steffin, D. et al. Interleukin-15-armoured GPC3 CAR T cells for patients with solid cancers. Nature 637, 940–946, doi:10.1038/s41586-024-08261-8 (2025).

27 Wang, X. et al. Comparative analysis of cell lineage differentiation during hepatogenesis in humans and mice at the single-cell transcriptome level. Cell Research 30, 1109–1126 (2020).

28 Hirsch, T. Z. et al. Integrated Genomic Analysis Identifies Driver Genes and Cisplatin-Resistant Progenitor Phenotype in Pediatric Liver Cancer. Cancer discovery 11, 2524–2543, doi:10.1158/2159-8290.Cd-20-1809 (2021).

29 Whitfield, M. L. et al. Identification of genes periodically expressed in the human cell cycle and their expression in tumors. Mol Biol Cell 13, 1977–2000, doi:10.1091/mbc.02-02-0030 (2002).

30 Subramanian, A. et al. Gene set enrichment analysis: a knowledge-based approach for interpreting genome-wide expression profiles. Proceedings of the National Academy of Sciences 102, 15545–15550 (2005).

31 Nielsen, M. M. & Pedersen, J. S. miRNA activity inferred from single cell mRNA expression. Scientific reports 11, 9170, doi:10.1038/s41598-021-88480-5 (2021).

32 Lorenzi, L. et al. The RNA Atlas expands the catalog of human non-coding RNAs. Nature biotechnology 39, 1453–1465 (2021).

33 Chiu, H.-S. et al. Pan-cancer analysis of lncRNA regulation supports their targeting of cancer genes in each tumor context. Cell reports 23, 297–312 (2018).

34 Ustianenko, D. et al. LIN28 selectively modulates a subclass of let-7 microRNAs. Molecular cell 71, 271–283. e275 (2018).

